# Core species and interactions prominent in fish-associated microbiome dynamics

**DOI:** 10.1101/2022.09.11.507499

**Authors:** Daii Yajima, Hiroaki Fujita, Ibuki Hayashi, Genta Shima, Kenta Suzuki, Hirokazu Toju

## Abstract

In aquatic ecosystems, the health of fish depends greatly on the dynamics of microbial community structure in the background environment. Nonetheless, finding microbes with profound impacts on fish’s performance out of thousands of candidate species remains a major challenge. We here show that time-series analyses of microbial population dynamics illuminate core components and structure of fish-associated microbiomes. By targeting eel aquaculture microbiomes as model systems, we reconstructed the population dynamics of 9,605 bacterial and 303 archaeal species/strains across 128 days. Due to the remarkable increase/decrease of constituent microbial populations, the taxonomic compositions of microbiomes changed drastically through time. We then found that some specific microbial taxa showed positive relationship with eels’ activity level even after excluding cofounding effects of environmental parameters (pH and dissolved oxygen level) on population dynamics. In particular, a vitamin B_12_-producing bacteria, *Cetobacterium somerae*, consistently showed strong positive associations with eels’ activity level across the replicate time-series of the five aquaculture tanks. Network theoretical and metabolic modeling analyses further suggested that the highlighted bacterium formed compartments of close microbe-to-microbe interactions with some other bacterial taxa, forming potential core microbiomes with positive impacts on eels. Overall, these results suggest that integration of microbiology, ecological theory, and network science allows us to explore core species and interactions embedded within complex dynamics of fish-associated microbiomes.

---

Microbial communities are essential factors of the life of vertebrates^1–4^, playing key roles in the development and homeostasis of their hosts^5–7^. Gut microbiomes, for example, play key roles in the nutrition and disease prevention of human and other mammal species^8,9^. Such physiological and ecological effects of gut microbes on hosts have been reported as well for fish^6,10,11^. Meanwhile, because fish are continuously exposed to numerous pathogenic and non-pathogenic microbial species in the water, their performance (or fitness) depends not only on gut-associated microbes^6,10^ but also on the microbiomes of the background environment^12–14^. Therefore, finding key microbiome components whose dynamics determine fish’s health or performance is of interdisciplinary interest spanning from microbiology to zoology and environmental science. However, due to the tremendous diversity of bacteria and archaea in aquatic ecosystems^15,16^, exploring such core microbial species associated with fish health remains a challenge.

A starting point for finding fish-health-associated microbes in aquatic ecosystems is to track the dynamics of microbial community compositions. Nonetheless, we still have limited knowledge of the extent to which structure of fish-associated microbiomes change through time. Although time-series data of microbiomes have become available in pioneering projects of human-associated microbes^17,18^, few attempts have been made to monitor microbiomes associated with other animals over tens of time points. Moreover, continuous sampling of fecal samples of targeted vertebrate individuals is generally much harder in aquatic environments than in terrestrial environments. Thus, developing model systems for time-series analyses of microbe–fish ecological interactions is a demanding but essential step for exploring core bacteria and/or archaea out of thousands of candidate species in microbial communities.

Despite the hardship in gaining time-series microbiome samples at the individual level, fish-associated microbiome dynamics can be monitored at the population or community level by sampling environmental water samples^19–21^. Because excrements of fish are released to water, samples of background water are expected to reflect gut microbiomes of fish populations or communities. Furthermore, as individual fish are continuously exposed to the background microbiomes, analyses of water samples provide essential insights into surrounding environmental conditions and potential sources of gut microbiomes^12–14^. In this respect, time-series analyses of aquaculture or aquarium systems offer an ideal opportunity for investigating relationship between microbial community structure, core microbial species, and vertebrate health.

By targeting a recirculating aquaculture system of the Japanese eel (*Anguilla japonica*), we herein integrate microbiology, community ecology, and network science for detecting key species and structure within fish-associated microbiomes. Based on the DNA metabarcoding of prokaryote (bacterial and archaeal) communities for the 128-day time-series, we revealed to what extent the compositions of aquaculture microbiomes fluctuate through time. We then reconstructed the population dynamics (i.e., increase/decrease) of the 9,908 microbial amplicon sequencing variants (ASVs) constituting the aquaculture microbiomes, screening bacteria or archaea whose abundance was tightly linked with the health condition of eels. We then found that several microbial ASVs showed positive associations with eel health consistently across the five replicate aquaculture tanks, even after controlling the effects of their environmental preference (e.g., preference to pH and dissolved oxygen level). With the approaches of network science and metabolic modeling, we further examined potential interactions between the core microbes. Overall, this study illustrates how core species and interactions are detected based on time-series datasets of microbiome dynamics.

## Results

### Microbiome dynamics

Monitoring of microbiome dynamics was conducted targeting the five water tanks of an aquaculture farm of the Japanese eel. In each water tank (diameter = 2.5 m; height = 1 m; volume = 20 m^3^), 1,400–4,300 eel individuals (average weight = 80–130 g) had been kept. The pH and dissolved oxygen (DO) concentrations were recorded for each tank every day. In addition, as a measure of ecosystem-level functions of microbiomes, the health condition of eels was evaluated based on eight criteria, yielding eel activity scores on a scale of 0 to 40 (see Methods). For the analyses of microbiome dynamics, water was sampled from each aquaculture tank every 24 hours during 128 days. By applying a quantitative amplicon sequencing approach for estimating 16S ribosomal RNA gene (16S rRNA) copy concentrations of respective microbes^22,23^, we obtained time-series datasets representing the increase/decrease of 9,605 bacterial and 303 archaeal ASVs representing 618 genera and 325 families (Fig. 1a). Thus, our data offered a novel opportunity to test synchronizations among microbial population dynamics, environmental factors (pH and DO), and vertebrate performance (eel activity level).

**Fig. 1.**
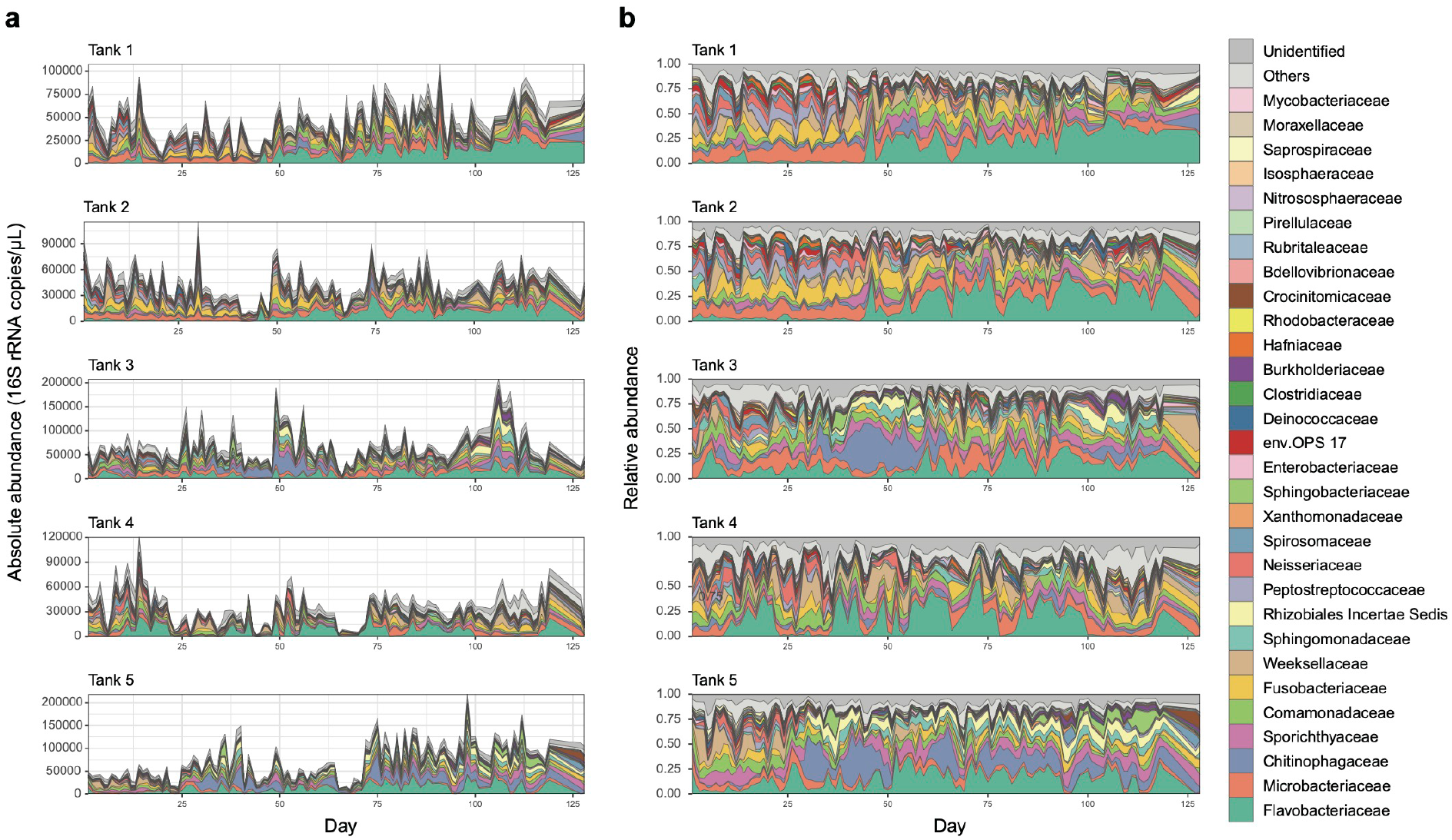
Microbiome dynamics in the eel aquaculture system. **a**, Dynamics of absolute abundance. For each water sample of each aquaculture tank, absolute abundance of prokaryotes was inferred as 16S rRNA gene copy concentration based on the quantitative amplicon sequencing approach with standard DNA gradients. **b**, Dynamics of relative abundance. The time-series of the family-level taxonomic compositions are shown for each aquaculture tank. See Extended Data Figures 1 –3 for phylum-, order-, and genus-level taxonomic compositions.

At the community level, drastic taxonomic turnover was observed in the timeseries of each aquaculture tank (Fig. 1; Extended Data Figs. 1-3). In Tanks 1 and 2, for example, the community structure characterized by the predominance of Fusobacteriaceae and Microbacteriaceae was suddenly altered by a Flavobacteriaceae-dominated state around Day 45 (Fig. 1). Meanwhile, microbiomes of Tanks 3–5 displayed more complex dynamics represented by frequent shifts between Flavobacteriaceae-dominated and Chitinophagaceae-dominated states, although clear classification of community states was difficult (Fig. 1).

A multivariate analysis of the prokaryote community structure indicated that the community state characterized by dominance of Fusobacteriaceae and Microbacteriaceae was associated with high eels’ activity (Fig. 2). In contrast, the Flavobacteriaceae-dominated and Chitinophagaceae-dominated states, which were observed in high-pH conditions, were associated with low eels’ activity (Fig. 2). At the genus level, the high-eel-activity-related state of dominance by Fusobacteriaceae and Microbacteriaceae was characterized by high relative abundance of *Cetobacterium*, which includes species potentially contribute to fish physiological homeostasis^24^. On the other hand, the Flavobacteriaceae-dominated and Chitinophagaceae-dominated states associated with low eels’ activity were represented by *Flavobacterium* and *Edaphobaculum*, respectively (Extended Data Fig. 4). Among the genera, *Flavobacterium* includes fish pathogens^25^, while *Edaphobaculum*^26^ has been poorly investigated in terms of their effects on fish physiology. These results suggest potential impacts of environmental microbiome dynamics on fish health/behavior in aquaculture systems.

**Fig. 2.**
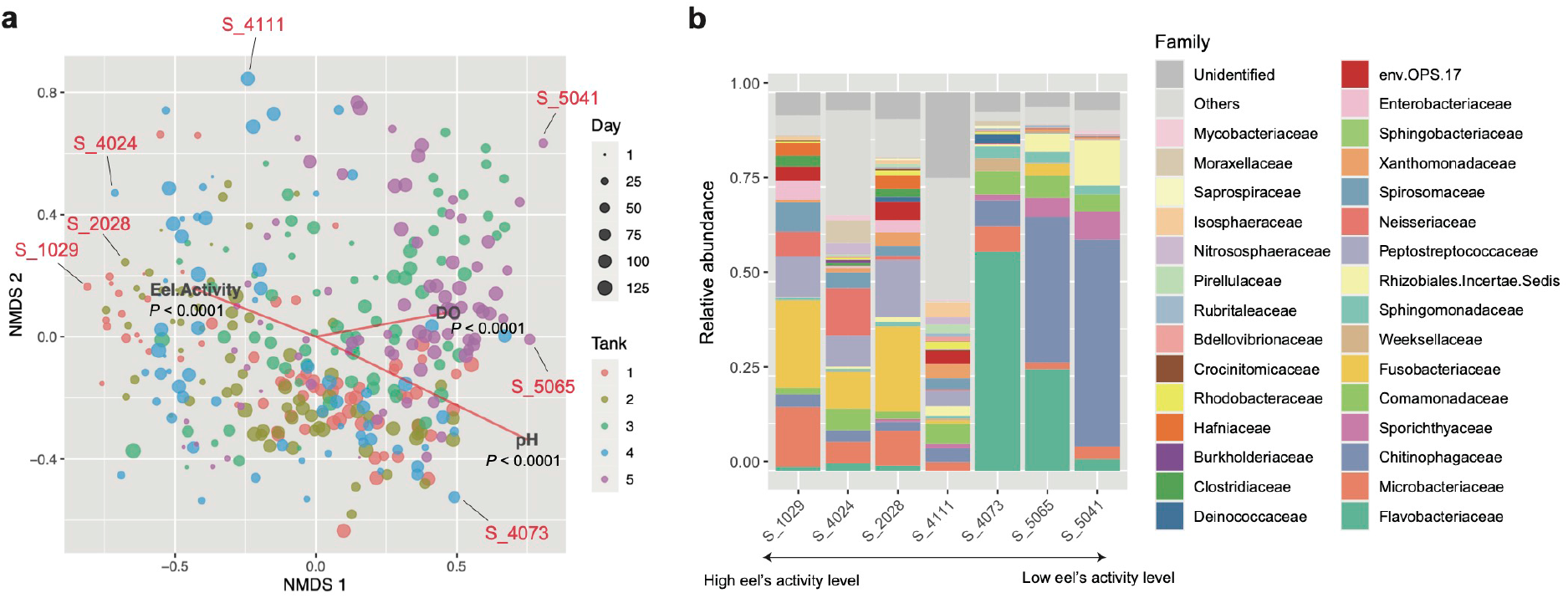
Multivariate analysis of community structure. **a**, Community state space. Community compositions of the samples are plotted on the two-dimensional surface defined with non-metric multidimensional scaling (NMDS). The NMDS was performed based on the Bray-Curtis -diversity of family-level taxonomic compositions. The projections of the data points onto the vectors have maximum correlation with the variables examined (pH, DO, and eels’ activity level). See Extended Data Figure 4 for an additional analysis based on genus-level taxonomic compositions. **b**, Examples of community structure in the NMDS surface. For several points within the NMDS surface (panel **a)**, family-level taxonomic compositions are shown. The example points are ordered along the vector representing high eels’ activity level.

#### Exploring microbes with key roles

We next evaluated how the population dynamics of each microbial ASV were associated with environmental variables and eels’ activity level. Specifically, we examined how population size (absolute abundance) of each ASV varied with pH, DO, and eels’ activity level (Fig. 3a) based on correlation analyses with twin-surrogate permutations^27^ (Fig. 3b-c). The ASVs varied in their environmental preference for pH and DO conditions as well as in their associations with eels’ activity level (Extended Data Fig. 5). We also found that ASVs’ relationship with eels’ activity level displayed tank-dependent complex associations with pH or DO preference (Fig. 3d-e). Thus, for each microbial ASV in each aquaculture tank, we calculated a partial correlation between absolute abundance and eels’ activity scores through the time-series by controlling the effects of pH or DO. Because partial correlation coefficients were consistent between the pH-controlled and DO-controlled calculations (Fig. 3f), the pH-controlled partial correlation coefficients were used in the following analyses.

**Fig. 3.**
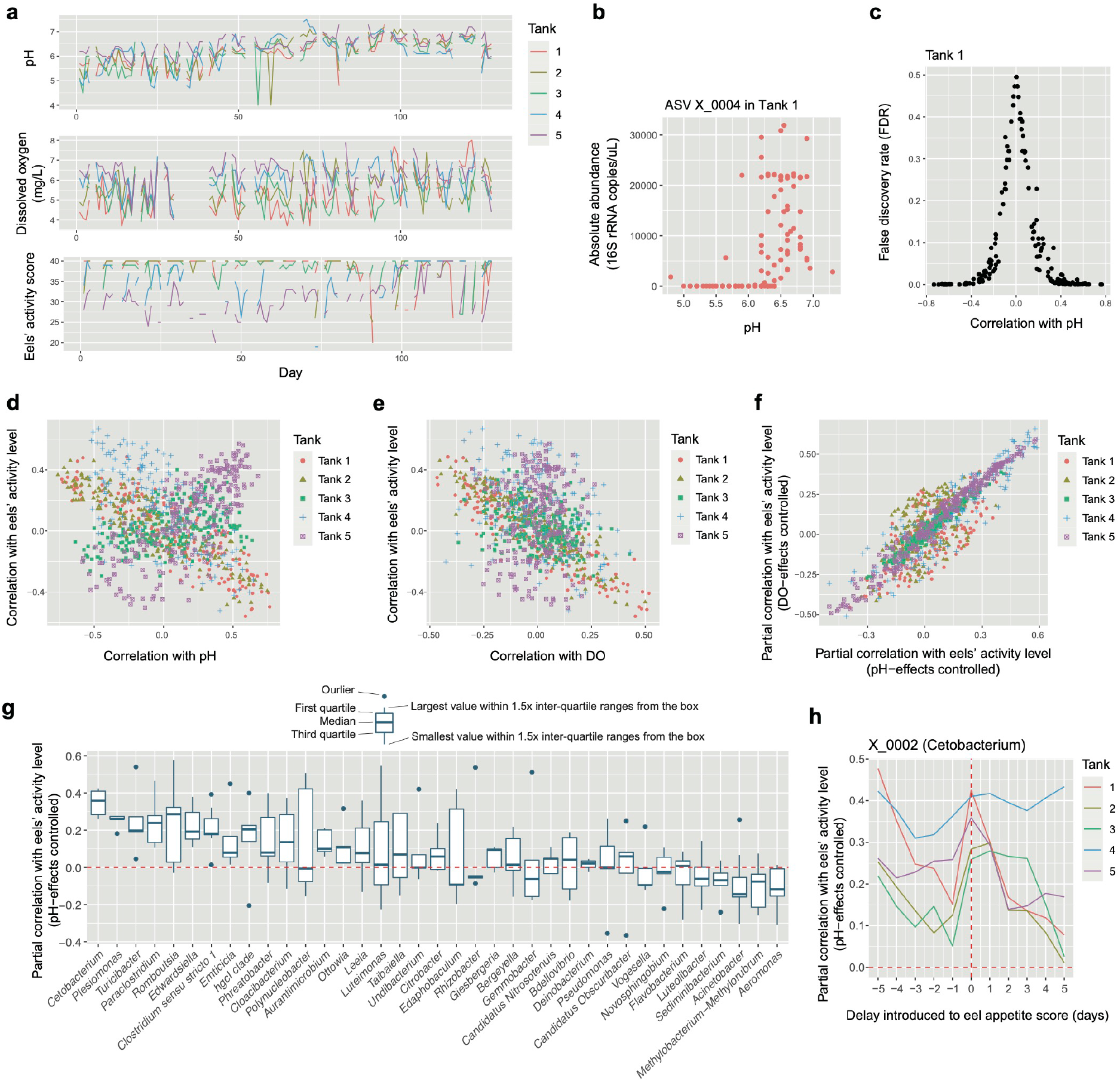
Microbes associated with eels’ activity. **a**, Timeseries of pH, dissolved oxygen (DO) level, and eels’ activity score are shown for each aquaculture tank. **b**, Example of the correlation analysis. For each variable shown in the panel **b**, Spearman’s correlation with the absolute abundance of each ASV in each aquaculture tank was examined. **c**, Randomization analysis of correlation. Significance of correlation coefficients was examined based on a twin-surrogate randomization analysis of time-series data (100,000 permutations). Coefficients less than -0.3 and those larger than 0.3 roughly represent significant negative and positive correlations, respectively. **d**, Each ASV’s correlation with pH and eels’ activity level. **e**, Each ASV’s correlation with DO and eels’ activity level. **f**, Partial correlation with eels’ activity level. To control the effects of pH or DO, partial correlation between absolute abundance and eels’ activity scores was calculated for each ASV in each tank. **g**, Taxonomic comparison of relationship with eels’ activity level. Partial correlation with eels’ activity level is shown for the genera that appeared in all the aquaculture tanks (shown in the decreasing order of mean values). **h**, Time-lag analysis of correlations. In calculating partial correlation between eels’ activity level and the absolute abundance of the *Cetobacterium* ASV (X_0002), defined time-lag was introduced to the eels’ activity variable.

The partial correlation coefficients with eels’ activity level varied greatly depending on prokaryote taxa (Fig. 2e). Nonetheless, ASVs belonging to some bacterial genera showed consistently positive correlation with eels’ activity scores across the five tanks (Fig. 2g; Extended Data Fig. 5). The list of those ASVs included bacteria belonging to the genera *Cetobacterium* (Fusobacteriaceae; Fusobacteriia; ASV ID = X_0002), *Plesiomonas* (Enterobacteriaceae; Gammaproteobacteria; X_0020), *Turicibacter* (Erysipelotrichaceae; Bacilli; X_0041), *Paraclostridium* (Clostridiaceae; Clostridia; X_0014), *Romboutsia* (Peptostreptococcaceae; Clostridia; X_0028), *Edwardsiella* (Hafniaceae; Gammaproteobacteria; X_0027), *Clostridium* (Clostridiaceae; Clostridia; X_0029), and an ASV belonging to Barnesiellaceae (Bacteroidia; X_0064) (Fig. 3g; Extended Data Fig. 5d).

An additional database search of the 16S rRNA sequences suggested that some of the ASVs with positive associations with eels’ activity level belonged to bacterial species with potential physiological impacts on fish. For example, the *Cetobacterium* ASV, which showed strongest positive partial correlation with eels’ activity level, was represented by the 16S rRNA sequences completely matching that of *Cetobacterium somerae* (formerly recognized as “*Bacteroides* type A”) in the NCBI nucleotide database. This *Cetobacterium* species has been known to produce high concentrations of vitamin B_12_ and hence their potential contributions to fish’s physiology have been anticipated. Meanwhile, the *Edwardsiella* ASV listed above was allied to the notorious fish pathogen *E. tarda*^28^, illuminating paradoxical relationships with eels’ health. However, our supplementary phylogenetic analysis based on the *sodB* gene marker^29^ indicated that 95.1 % of *Edwardsiella* bacteria detected in the focal eel aquaculture system belonged to non-pathogenic clades^29,30^ within the genus *Edwardsiella* (Extended Data Fig. 6).

In terms of negative impacts on eels’ activity level, bacteria in the genera *Aeromonas* (Aeromonadaceae; Gammaproteobacteria), *Methylobacterium* (alternatively, *Methylorubrum*; Beijerinckiaceae; Alphaproteobacteria), and *Acinetobactor* (Moraxellaceae; Gammaproteobacteria) were highlighted (Fig. 3g). Among them, *Aeromonas* and *Acinetobactor* have been known to include fish pathogens^31,32^. At the ASV level, an ASV allied to the cvE6 clade within the order Chlamydiales (Chlamydiae; Verrucomicrobiota) showed strongest negative correlation with eels’ activity scores (Extended Data Fig. 5d).

Although the above analysis controlling environmental preferences of respective bacteria allows high-throughput screening for species with potential positive/negative impacts on target biological functions, the simple statistical approach with partial correlation analyses precludes insights into the direction of causality. Specifically, it is important to consider the possibility that high/low abundance of an ASV is a consequence but not a cause of eels’ high/low activity. Therefore, we performed an additional analysis introducing time lags into eels’ activity scores throughout the time-series. We then found that the abundance of the *Cetobacterium* ASV was positively correlated with eels’ activity scores of the next day, while correlations between *Cetobacterium* abundance and past eels’ activity scores were much lower than those with no time lags (Fig. 3h). Meanwhile, high correlation between 5-days-ago eels’ activity level and present-day *Cetobacterium* abundance was observed in some tanks (Tanks 1 and 4; Fig. 3h), illuminating the importance of carefully interpreting the results of the time-series analysis.

#### Networks of interactions

We then reconstructed webs of potential microbe-to-microbe interactions to illuminate microbial groups or interactions positively associated with eels’ health. We first applied the Meinshausen-Bühlmann (MB) method, which was designed to evaluate patterns of coexistence realized by the effects of microbe–microbe interactions as well as those of niche sharing between microbes. For each aquaculture tank, the reconstructed network of microbe–microbe coexistence (Extended Data Figs. 7–8; Supplementary Table 1) was compartmentalized into several modules, which differed in mean partial correlations with eels’ activity scores (Fig. 4). We then found that each of the five networks included a module constituted by the abovementioned *Cetobacterium* ASV and several other ASVs with consistently positive associations with eels’ activity level (Fig. 4; Extended Data Fig. 9). The bacteria consistently formed network modules of coexistence with the *Cetobacterium* ASV were *Plesiomonas* (X_0020), *Turicibacter* (X_0041), *Paraclostridium* (X_0014), *Romboutsia* (X_0028), *Edwardsiella* (X_0027), and *Clostridium* (X_0029) (Fig. 4).

**Fig. 4.**
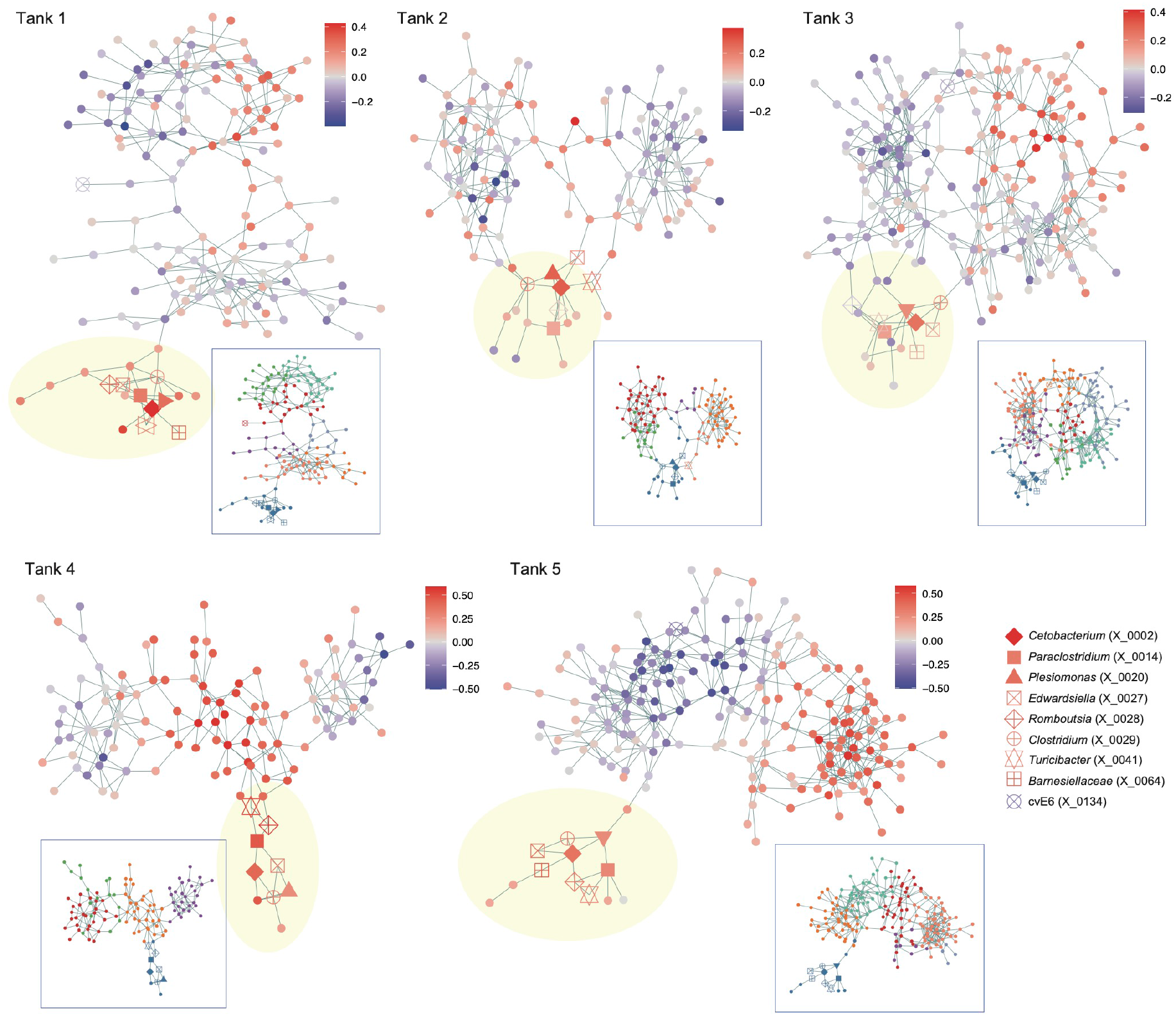
Microbe-to-microbe coexistence networks. For each aquaculture tank, patterns of coexistence were analyzed based on the sparse inverse covariance estimation for ecological associations with the Meinshausen-Bühlmann (MB) model. Only the ASVs that appeared in 30 or more samples were targeted in the analysis of each tank. Within the networks, pairs of microbial ASVs that may interact with each other in facilitative ways and/or those potentially sharing environmental preference are linked with each other. Network modules, which represent groups of densely linked ASVs, are shown for each network. The color of nodes indicates partial correlation between ASV abundance and eels’ activity level (controlled variable = pH). The inferred network modules are shown by colors for each tank in a box. The ASVs that consistently displayed positive or negative correlation with eels’ activity level (Extended Data Fig. 5) are highlighted with the defined symbols. See Extended Data Figures 6–8 for additional information of the nodes (ASVs) and modules within the network. ASVs included in minor sub-networks (number of nodes < 5) are not shown.

To infer the presence/absence of direct interactions between these bacteria with positive relationship with eels’ activity level, we conducted an additional network analysis based on the sparse and low-rank (SLR) decomposition method, which allowed us to remove latent effects of environmental conditions. In the networks reconstructed with the SLR method (Fig. 5), potential effects of niche sharing were controlled and hence the links between bacterial ASVs were expected to represent potential positive interactions. The estimated interaction coefficients were highly correlated between the MB and SLR methods (Extended Data Fig. 10). Meanwhile, in the SLR-based network, removing the effects of potential niche sharing (sharing of environmental preference) resulted in the simplification of network structure, in which estimated direct interactions between microbes were focused (Fig. 5). Despite the considerable difference between MB- and SLR-based network topology, the *Cetobacterium* ASV with the strongest associations with eels’ activity level was, again, linked with the *Plesiomonas, Turicibacter, Paraclostridium, Romboutsia, Edwardsiella*, and *Clostridium* ASVs within the SLR network (Fig. 5), suggesting positive interactions with these bacteria.

**Fig. 5.**
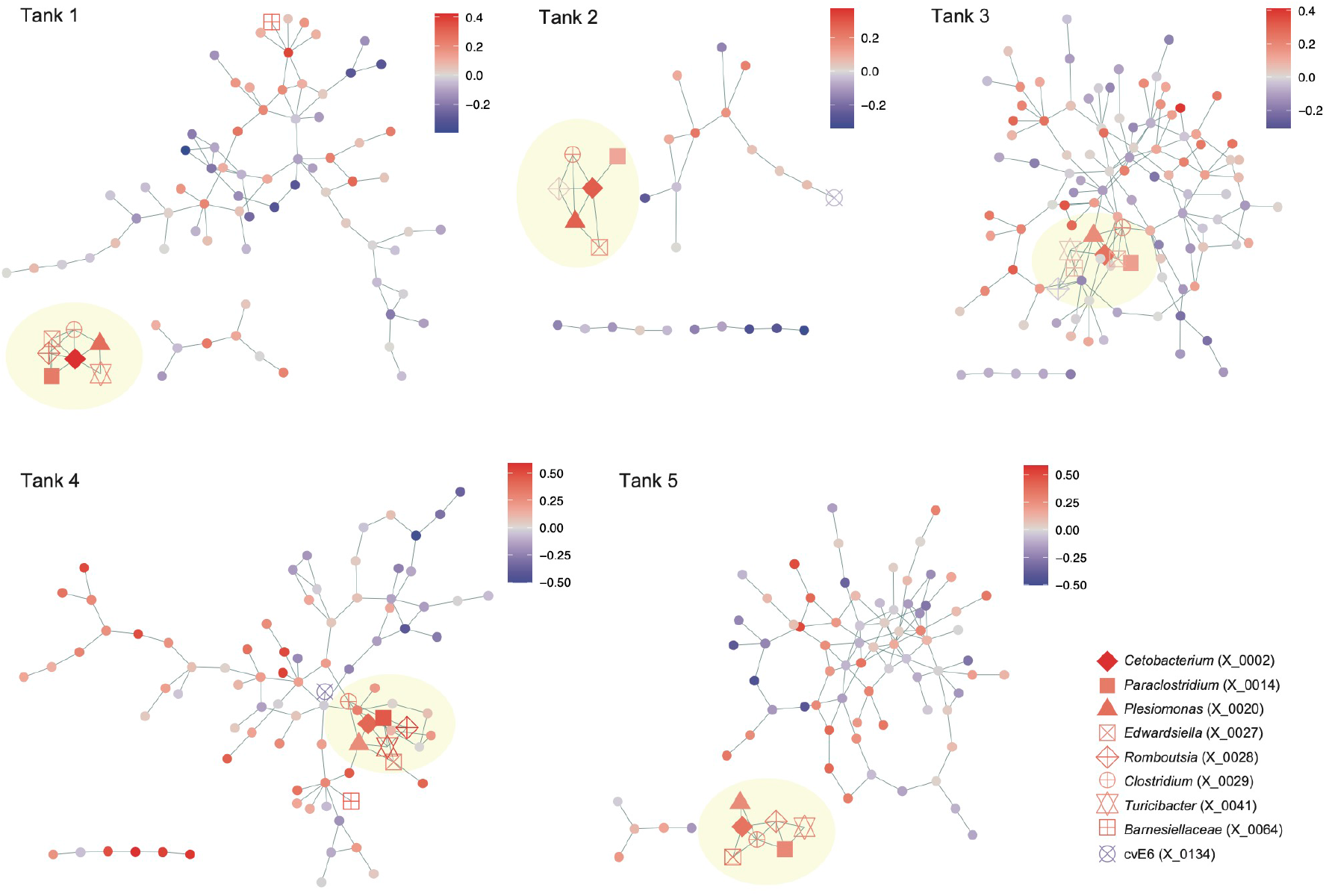
Inferred direct interactions between microbes. Based on the “sparse and low-rank” (SLR) model, direct interactions between microbial ASVs were inferred by controlling the effects of shared environmental preference. Only the ASVs that appeared in 30 or more samples were targeted in the analysis of each tank. The links between nodes represent potentially positive interactions between ASVs. The color of nodes indicates partial correlation between ASV abundance and eels’ activity level (controlled variable = pH). ASVs included in minor sub-networks (number of nodes < 5) are not shown.

#### Potential metabolic interactions

To estimate functional interactions between microbes, we focused on genomic compositions of respective microbes within the aquaculture microbiomes. After retrieving the information of genomic compositions from reference databases, we analyzed the inferred gene repertoires (KEGG metabolic pathway/process profiles) of the microbial ASVs based on multivariate analysis. Along the principal component axes, *Cetobacterium*, which showed consistent correlations with eels’ activity (Fig. 3g; Extended Data Fig. 5d), was located distantly from *Edwardsiella, Plesiomonas*, and *Turicibacter* (Fig. 6a). In contrast, *Romboutsia, Paraclostridium*, and *Clostridium* displayed similar metabolic gene repertories with *Cetobacterium* (Fig. 6a).

**Fig. 6.**
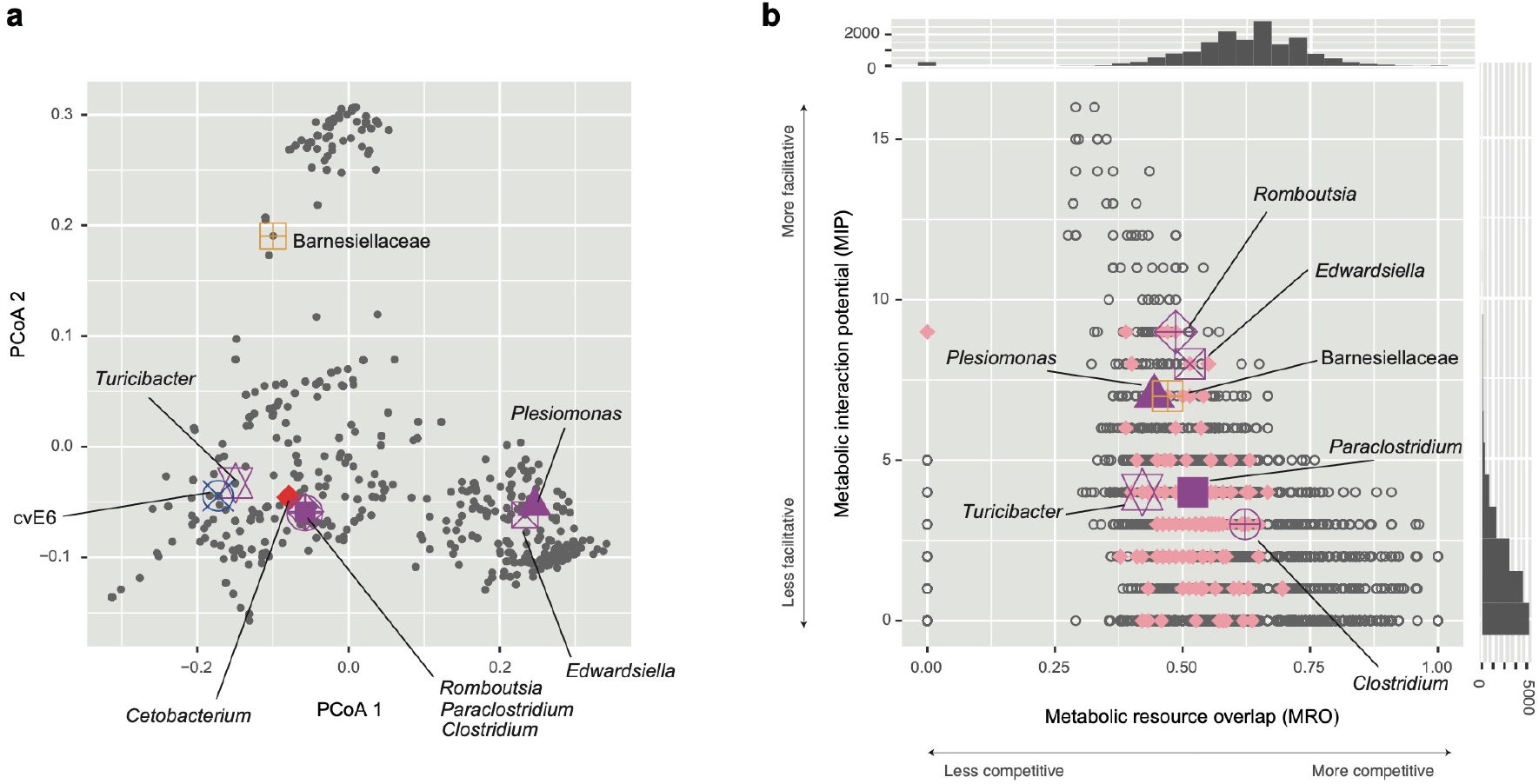
Metabolic interactions between microbes. **a**, Metagenomic niche space. Microbial ASVs are plotted on a two-dimensional surface of PCoA based on their KEGG metabolic pathway/process profiles inferred with a phylogenetic prediction of genomes. Microbial ASVs plotted closely within the surface are expected to have similar gene repertoires. The ASVs highlighted in Figures 4 and 5 are shown with large symbols. **b**, Potential competitive and facilitative interactions. Based on the NCBI RefSeq genome information, potential metabolic interactions between each pair of ASVs were inferred in terms of metabolic resource overlap (MRO) and metabolic interaction potential (MIP). Histograms of MRO and MIP are shown on the horizontal and vertical axes, respectively. ASV pairs including the *Cetobacterium* ASV, whose abundance were positively associated with eels’ activity level in all the five water tanks (Fig. 3g; Extended Data Fig. 5), are shown in pink. Relationships between the *Cetobacterium* ASV and the ASVs highlighted in Figures 4 and 5 are indicated as well.

We next evaluated potential competitive and facilitative interactions between microbes based on a genome-scale metabolic modeling approach. In the analysis, reference genomic information was used to infer competition for available resources and exchanges of metabolites, yielding metabolic resource overlap and metabolic interaction potential scores for each pair of microbial ASVs. We then found that the *Romboutsia, Edwardsiella*, and *Plesiomonas* ASVs had relatively low metabolic resource overlap and relatively high metabolic interaction potential with the *Cetobacterium* ASV among the prokaryotes examined (Fig. 6b).

## Discussion

Through the 128-day monitoring of thousands of microbial species/strains, we here found that aquatic microbiomes associated with fish could show drastic shifts of community structure through time. Such dynamical nature of community processes has been intensively investigated in human-associated microbiomes in light of potential influence on host status^17,18^. In particular, shifts (collapse) of microbial community structure to disease-related states (i.e., dysbiosis) have been considered as essential mechanisms determining human health^33,34^. Given the growing literature on microbiome dynamics in medical science, knowledge of shifts between alternative states of fish-related microbiomes^14^ is expected to shed new lights on physiological and ecological processes of vertebrates.

The aquaculture microbiome dynamics were described as shifts among Fusobacteriaceae-abundant states, Flavobacteriaceae-dominated states, and Chitinophagaceae-dominated states, although intermediate states existed through the time-series (Fig. 1). Among them, Fusobacteriaceae-abundant states, which were characterized by high abundance of *Cetobacterium*, were designated as microbiome compositions positively associated with eels’ activity level (Fig. 2; Extended Data Fig. 4). In fact, among the 9,908 microbial ASVs examined, the ASV representing *Cetobacterium somerae* showed the strongest associations with eel’s activity level through the time-series even after controlling effects of environmental preference (Fig. 3; Extended Data Fig. 5). This *Cetobacterium* species has been reported from a broad taxonomic range of freshwater fish^35–37^, especially from intestines of species that do not require dietary vitamin B_1224_. Although vitamin B_12_ (cobalamin) plays essential roles in animal physiology (e.g., normal functioning of nervous systems and the maturation of red blood cells), they can be synthesized only by specific clades of bacteria and archaea^38,39^. Genomic studies have shown that *C. somerae* has a series of genes for anaerobic vitamin B_12_ biosynthesis^40^. Indeed, the bacterium produces highest concentrations of vitamin B_12_ compared to other culturable bacteria within freshwater fish-associated microbiomes^24,41^. Given the prevalence of *Cetobacterium* in freshwater fish species^35–37^, our results suggest that maintaining microbiomes at *Cetobacterium*-abundant states is the key to build general platforms for stably keeping freshwater aquaculture/aquarium systems.

Further analyses based on network theory and metabolic modeling indicated the possibility that the *Cetobacterium* species form facilitative interactions with some other microbial species/ASVs (Figs. 4-6). Among the bacteria for which interactions with *Cetobacterium* were inferred from multiple analyses, *Edwardsiella tarda* has been known to include notorious pathogens of broad taxonomic groups of fish including eels^28,29,42^. However, we found that the *E. tarda* population of the investigated aquaculture system was dominated by non-pathogenic strains^29,30^ of the species (Extended Data Fig. 6). Thus, the presence of microbial species/strains belonging to broadly-known taxa of pathogens do not necessarily result in negative impacts on fish. Rather, our analyses suggested that “seemingly pathogenic” microbes could be involved in core microbiome components (network modules) constituted by microbes contributing to the maintenance of fish health. Further studies are awaited to explore potential mechanisms such as competitive exclusion of pathogenic strains by non-pathogenic strains^43,44^ or indirect negative impacts on pathogenic strains through the activation of fish immune systems^6,45^ by non-pathogenic strains. In contrast to *E. tarda, Romboutsia* and *Plesiomonas*, which were inferred as microbes with facilitative interactions with *C. somerae*, too (Figs. 4-6), have been poorly investigated in terms of their functions. Their potential roles in competitive exclusion of pathogens or activation of host immune systems deserve further investigations.

While the time-series dataset allowed us to highlight core species and interactions within microbial communities, more sophisticated statistical platforms beyond simple correlational approaches are necessary for confirming causative relationships between microbiome dynamics and vertebrate health/performance. In this respect, methods based on nonlinear mechanics, such as transfer entropy and empirical dynamic modeling^46,47^, are expected to help us infer causative interactions among microbial population dynamics, environmental factors, and vertebrate performance. Albeit promising, these methods require substantial computational resources when we try to analyze microbiomes consisting of thousands of ASVs. Further methodological advances will deepen our understanding of the mechanisms by which microbiome dynamics and vertebrate performance are linked with each other.

As the analyses of microbiome dynamics extend from medical science to researches targeting other vertebrates, we will be more and more aware of overlooked roles of microbes in both terrestrial and aquatic ecosystems. Feedback between intestine and environmental microbiomes, for example, deserves future intensive research in terms of potential great impacts on ecosystem processes. In particular, given that aquatic vertebrates are continuously exposed to excrements of other individuals or species, their gut microbiome dynamics (and related health conditions) may be more likely to be synchronized at the population or community levels than those of terrestrial vertebrates. Therefore, simultaneous monitoring of intestine and background environmental microbiomes will provide platforms for uncovering such feedback and synchronization processes. Further insights into fish-associated microbiome dynamics will reorganize our basic understanding of aquatic ecosystem dynamics, advancing technologies for sustainable food production through stable aquaculture systems^48–50^.

## Methods

### Sampling

Monitoring of microbiome dynamics was conducted targeting the five water tanks of the eel aquaculture system of A-Zero Inc. (Nishiawakura, Okayama Prefecture, Japan). In each water tank (diameter = 2.5 m; height = 1 m; volume = 20 m^3^), 1,400–4,300 eel individuals (average weight = 80–130 g) had been kept. About 10 % of tank water was replaced with warmed fresh well water every day, and the water temperature in the tanks was kept at around 30 ºC. The drainage from the five tanks were mixed and processed in a series of filtration equipment. The filtered drainage was returned to each tank after being processed in another filtration equipment adjacent to each tank. The eels were fed with mixture of commercial artificial diets. The pH, dissolved oxygen (DO), and eels’ activity level were recorded for each tank every day. The eels’ activity level was evaluated based on the sum of the scores of the following eight criteria: initial responses to feeders, the proportion of eels responding to feeders, sharpness of movement, the proportion of eels eating the artificial diet, the level of splashes, the amount of scattered diet, the time to consume the diet, and the proportion of foraging eels at the end of feeding. For each of the criteria, scoring was done on a five-point scale (maximum point = 5) by an expert of eel aquaculture maintenance: thus, 40 (5 × 8 criteria) is the maximum point of the eels’ activity score. Albeit subjective, the criteria evaluated continuously by a professional provide inferences of eel’s health conditions throughout the time series. The water in the tanks were continuously mixed by the movement of eels.

From each aquaculture tank (Tank 1–5), ca. 1.5 mL of water was sampled in the morning every day during the 128 days from March 25 to July 30, 2020, except for 9 days (Days 102, 103, 120, 121, 122, 123, 124, 125, and 126): i.e., the samples of 119 days were available. In Tank 4, samples were unavailable on additional three days (Day 67–69) due to the cleaning and the entire replacement of water. Consequently, the number of collected samples were 592 (119 days ×5 tanks – 3 days in Tank 4). Water sample was collected in a 2.0 mL microtube and they were immediately stored at -20 ºC in a freezer until DNA extraction.

#### Quantitative 16S rRNA sequencing

To extract DNA from each sample, 250 μL of the collected water was mixed with mixed with 400 μL lysis buffer (0.0025 % SDS, 20 mM Tris (pH 8.0), 2.5 mM EDTA, and 0.4 M NaCl) and 250 μL 0.5 mm zirconium beads in a 2.0 mL microtube. The microtubes were then shaken at 25 Hz for 5 min using TissueLyser II (Qiagen, Venlo). After centrifugation, the aliquot was mized with proteinase K solution (×1/100 of the total volume), being incubated at 40 ºC for 60 min followed by 95 ºC for 5 min.

We then performed PCR by applying a quantitative amplicon sequencing method^22,51^. Although most existing microbiome studies were designed to infer “relative” abundance of microbial amplicon sequence variants (ASVs) or operational taxonomic units (OTUs), information of “absolute” abundance provide additional insights into microbiome dynamics: i.e., insights into increase/decrease of the population size of each prokaryote ASV/OTU within a microbiome throughout a time-series^22^. The quantitative amplicon sequencing approach is based on the addition of artificial (standard) DNA sequences with defined concentrations into PCR master solutions. Therefore, even if compositions or concentrations of PCR inhibitor molecules in DNA extracts vary among time-series samples, potential bias caused by such inhibitors can be corrected based on the use of the internal standards (i.e., standard DNAs within PCR master solutions).

Prokaryote 16S rRNA region was PCR-amplified with the forward primer 515f^52^ fused with 3–6-mer Ns for improved Illumina sequencing quality and the forward Illumina sequencing primer (5’-TCG TCG GCA GCG TCA GAT GTG TAT AAG AGA CAG-[3–6-mer Ns] – [515f] -3’) and the reverse primer 806rB^53^ fused with 3–6-mer Ns for improved Illumina sequencing quality^54^ and the reverse sequencing primer (5’-GTC TCG TGG GCT CGG AGA TGT GTA TAA GAG ACA G [3–6-mer Ns] - [806rB] -3’) (0.2 μM each). To apply the quantitative amplicon sequencing, five standard DNA sequence variants with different concentrations of artificial 16S rRNA sequences (0.1, 0.05, 0.02, 0.01, and 0.005 nM) were added to PCR master mix solutions^22^. The buffer and polymerase system of KOD One (Toyobo) was used with the temperature profile of 35 cycles at 98 ºC for 10 s, 55 ºC for 30 s, 68 ºC for 30 s. To prevent generation of chimeric sequences, the ramp rate through the thermal cycles was set to 1 ºC/sec^55^. Illumina sequencing adaptors were then added to respective samples in the supplemental PCR using the forward fusion primers consisting of the P5 Illumina adaptor, 8-mer indexes for sample identification^56^ and a partial sequence of the sequencing primer (5’-AAT GAT ACG GCG ACC ACC GAG ATC TAC AC - [8-mer index] - TCG TCG GCA GCG TC -3’) and the reverse fusion primers consisting of the P7 adaptor, 8-mer indexes, and a partial sequence of the sequencing primer (5’-CAA GCA GAA GAC GGC ATA CGA GAT - [8-mer index] - GTC TCG TGG GCT CGG -3’). KOD One was used with a temperature profile: followed by 8 cycles at 98 ºC for 10 s, 55 ºC for 30 s, 68 ºC for 30 s (ramp rate = 1 ºC/s). The PCR amplicons of the samples were then pooled after a purification/equalization process with the AMPureXP Kit (Beckman Coulter). Primer dimers, which were shorter than 200 bp, were removed from the pooled library by supplemental purification with AMpureXP: the ratio of AMPureXP reagent to the pooled library was set to 0.6 (v/v) in this process. Because the quality of forward sequences is generally higher than that of reverse sequences in Illumina sequencing, we optimized the MiSeq run setting in order to use only forward sequences. Specifically, the run length was set 271 forward (R1) and 31 reverse (R4) cycles to enhance forward sequencing data: the reverse sequences were used only for screening 16S rRNA sequences in the following bioinformatic pipeline.

#### Bioinformatics

In total, 16,298,203 sequencing reads were obtained in the Illumina sequencing. The raw sequencing data were converted into FASTQ files using the program bcl2fastq 1.8.4 distributed by Illumina. The raw sequencing data were converted into FASTQ files using the program bcl2fastq 1.8.4 distributed by Illumina. The output FASTQ files were demultiplexed using Claident v0.2. 2018.05.29^57^. The removal of low-quality sequences and ASV inferences were done using DADA2^58^ v.1.22.0 of R 4.1.2^59^(R Core Team, 2020). The taxonomy of the output ASVs was inferred based on the naive Bayesian classifier method^60^ using the SILVA v.138 database^61^. Based on the calibration with the concentration gradients of the five standard DNAs, concentrations of respective ASVs were obtained for each sample (16S rRNA copy numbers per unit volume of tank water samples; copies/μL). As the number of 16S rRNA copies per genome generally varies among prokaryotic taxa^62^, 16S rRNA copy concentration is not directly the optimal proxy of cell or biomass concentration. Meanwhile, in this study, estimates of 16S rRNA copy concentrations were used to observe increase/decrease of abundance (i.e., population dynamics) *within* the time-series of respective microbial ASVs. Thus, variation in the number 16S rRNA copy numbers among microbial taxa had no qualitative effects on the subsequent population- and community-ecological analyses. The samples in which Pearson’s coefficients of correlations between sequencing read numbers and standard DNA copy numbers (i.e., correlation coefficients representing calibration curves) were less than 0.8 were removed as those with unreliable estimates. Samples with less than 1,000 reads were discarded as well. In total, microbiome data were successfully obtained from 577 out of 592 samples. For each aquaculture tank, we then obtained a sample ×ASV matrix, in which a cell entry depicted the concentration of 16S rRNA copies of an ASV in a sample.

#### Community structure

For each aquaculture tank, Bray-Curtis *β*-diversity was calculated for all pairs of time points based on the matrix describing the relative abundance of prokaryote families using the vegan 2.6.2 package^63^ of R. Based on the -diversity estimates, the community structure of all the samples across the five water tanks were visualized on the surface of non-metric multidimensional scaling (NMDS). The vectors representing the environmental variables (pH and DO) and eels’ activity level were calculated with the “envfit” function of R and they were shown on the NMDS surface. The analysis was conducted as well based on the matrix describing the relative abundance of genera.

#### Environmental preference of ASVs

To evaluate environmental preference of each microbial ASV, Spearman’s correlation between absolute abundance (in the metric of DNA copy numbers of 16S rRNA) and pH was calculated. For each tank, the ASVs that appeared in 30 or more samples were subjected to the analysis. For each ASV in each water tank, the statistical significance of the obtained correlation coefficient was examined with a randomization analysis obtained based on a twin-surrogate method for time-series data^27^ (100,000 permutations). Correlation coefficients less than -0.3 and those larger than 0.3 tended to show statistically significant negative and positive correlations with pH, respectively, after Benjamini-Hochberg adjustment of *P* values in multiple testing [i.e., false discovery rate (FDR)]. Likewise, Pearson’s correlation coefficients between respective ASVs’ absolute abundance and DO concentrations were calculated.

#### ASV abundance and eel’s activity

We explored microbial ASVs that potentially have profound impacts on eels’ health. For each water tank, Spearman’s correlation between absolute abundance and eels’ activity score was calculated for the ASVs that appeared in 30 or more samples. However, because ASV abundance could be affected by pH or dissolved oxygen concentration, the use of such simple correlation coefficients might be misleading. Therefore, we controlled potential effects by environmental factors/conditions based on a partial correlation approach as follows:

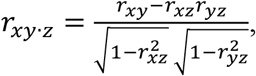

where *r*_*xy*_, *r*_*xz*_, and *r*_*yz*_ were correlation between ASV abundance and eels’ activity level, that between ASV abundance and an environmental factor (pH or dissolved oxygen concentration), and that between eels’ activity level and an environmental factor, respectively. For each ASV, a randomization analysis was performed with the twin-surrogate method (100,000 permutations).

#### Time-lag analysis

We extended the analysis of partial correlation between microbial abundance and eels’ activity level by introducing time lags between the two variables. Specifically, partial correlation between an ASV’s abundance on Day *x* and eels’ activity score on Day *x + l* was calculated. The time lag *l* ranged from -5 to 5 in the analysis (*l* = 0 means no delay introduced to eels’ activity level).

#### Pathogenic and non-pathogenic *Edwardsiella*

We performed an additional analysis to infer the proportion of pathogenic and non-pathogenic clades^29,30^ of *Edwardsiella* bacteria in the aquaculture system. In a previous phylogenetic study based on an internal fragment of iron-cofactored superoxide dismutase gene (*sodB*), *Edwardsiella* species and strains have been classified into two major clades, which differ in the presence of pathogenicity to fish (hereafter, “pathogenic” and “non-pathogenic” clades). Therefore, we characterized *Edwardsiella* bacteria in the aquaculture tanks based on the illumina sequencing of the *Edwardsiella sodB* gene sequences. The fragment of the *sodB* region was PCR-amplified with the forward primer E1F^29^ fused with 3–6-mer Ns for improved Illumina sequencing quality and the forward Illumina sequencing primer (5’-TCG TCG GCA GCG TCA GAT GTG TAT AAG AGA CAG-[3–6-mer Ns] – [E1F] -3’) and the reverse primer 497R^29^ fused with 3–6-mer Ns for improved Illumina sequencing quality^54^ and the reverse sequencing primer (5’-GTC TCG TGG GCT CGG AGA TGT GTA TAA GAG ACA G [3–6-mer Ns] - [497R] -3’) (0.2 μM each). The buffer and polymerase system of KOD One (Toyobo) was used with the temperature profile of 35 cycles at 98 ºC for 10 s, 55 ºC for 5 s, 68 ºC for 30 s (ramp rate = 1 ºC/sec). The sequencing adaptors and sample identifier indexed were added to the amplicons, and the purification of the library and sequencing was performed as detailed above.

The output sequencing data were demultiplexed and processed with DADA2. The ASVs that were not aligned to the *sodB* sequences of *Edwardsiella*^29^ were discarded. The neighbor-joining tree of the remaining ASVs and previously reported *Edwardsiella* sequences was reconstructed based on the maximum composite likelihood method with a bootstrap test (1,000 permutations). The ASVs belonging to the pathogenic clade and those belonging to the non-pathogenic clade of *Edwardsiella* were distinguished within the phylogeny.

#### Microbe–microbe interactions

Potential positive/negative interactions between microbial ASVs were inferred based on the framework of sparse inverse covariance estimation for ecological associations (SPIEC-EASI^64^). For each water tank, patterns in the coexistence (co-occurrence) were examined with the Meinshausen-Bühlmann (MB) method as implemented in the SpiecEasi package^64^ of R. The network inference based on coexistence patterns allowed us to detect pairs of microbial ASVs that potentially interact with each other in facilitative ways and/or those potentially sharing environmental preference. Because estimation of coexistence patterns was not feasible for rare nodes, the microbial ASVs that appeared in less than 30 samples were excluded from the input matrices of the network analysis. Network modules, within which closely associated ASVs were interlinked with each other, were identified with the algorithm based on edge betweenness using the igraph package^65^ of R. For each network module in each water tank, mean partial correlation with eels’ activity level was calculated across ASVs constituting the module.

In addition to the networks representing whole coexistence patterns, we reconstructed networks depicting direct interactions between microbial ASVs. To separate effects of direct microbe–microbe interactions from those of shared environmental preferences between microbes (i.e., shared niches), 10 latent components (latent variables) were included in the analysis based on the “sparse and low-rank” (SLR) model^66^.

#### KEGG pathway/process profiles

To infer metabolic interactions between microbial ASVs, we performed a series of analysis based on reference genome information. We performed phylogenetic prediction of gene repertoires using PICRUSt2 v2.3.0-b^67^ in order to gain the overview of the niche space defined with metagenomic information^68,69^. ASVs that appeared in 30 or more sample across the five tanks were subjected to the analysis. Based on the inferred KEGG metabolic pathway/process profiles^70^, microbial ASVs were plotted on a two-dimensional surface of a principal coordinate analysis (PCoA) based on Bray-Curtis *β*- diversity of KEGG metabolic pathway/process profiles.

#### Metabolic modeling

To infer potential metabolic interactions between microbes, we performed the species metabolic interaction analysis^71^. For the ASVs that appeared in 30 or more samples (day) in at least one aquaculture tank, we explored NCBI RefSeq genome sequences whose 16S rRNA sequences matched those of query ASVs with ≥ 99 % identity. In the database exploration, reference genome information was available for 181 out of 417 ASVs examined. The reference genome information was subjected to genome-scale metabolic modeling as implemented in CarveMe^72^ 1.5.0. Metabolic resource overlap (MRO) and metabolic interaction potential (MIP) were then estimated for each pair of microbial ASVs as implemented in SMETANA^71^ 1.0.0.

## Data availability

The 16S rRNA sequencing data are available from the DNA Data Bank of Japan (DDBJ) with the accession number PRJDB14313. The microbial community data are deposited at the GitHub repository (https://github.com/hiro-toju/EelMicrobiome128).

## Code availability

All the R scripts used to analyze the data are available at the GitHub repository (https://github.com/hiro-toju/EelMicrobiome128).

## Acknowledgements

We thank Yuta Nogi and A Zero inc. for support in sampling. JSPS Grant-in-Aid for Scientific Research (20K20586), and JST FOREST (JPMJFR2048) to H.T., JSPS Grant-in-Aid for Scientific Research (20K06820 and 20H03010) to K.S., and JSPS Fellowship to H.F..

## Author Contributions

H.T. designed the work with D.Y.. D.Y., I.H., G.S., and H.F. performed experiments. D.Y., H.F., K.S., and H.T. analyzed the data. H.T. and D.Y. wrote the paper with all the authors.

## Competing Interests

The authors declare no competing interests.

**Correspondence and requests for materials** should be addressed to.

## Extended Data Figures

**Extended Data Fig. 1.**
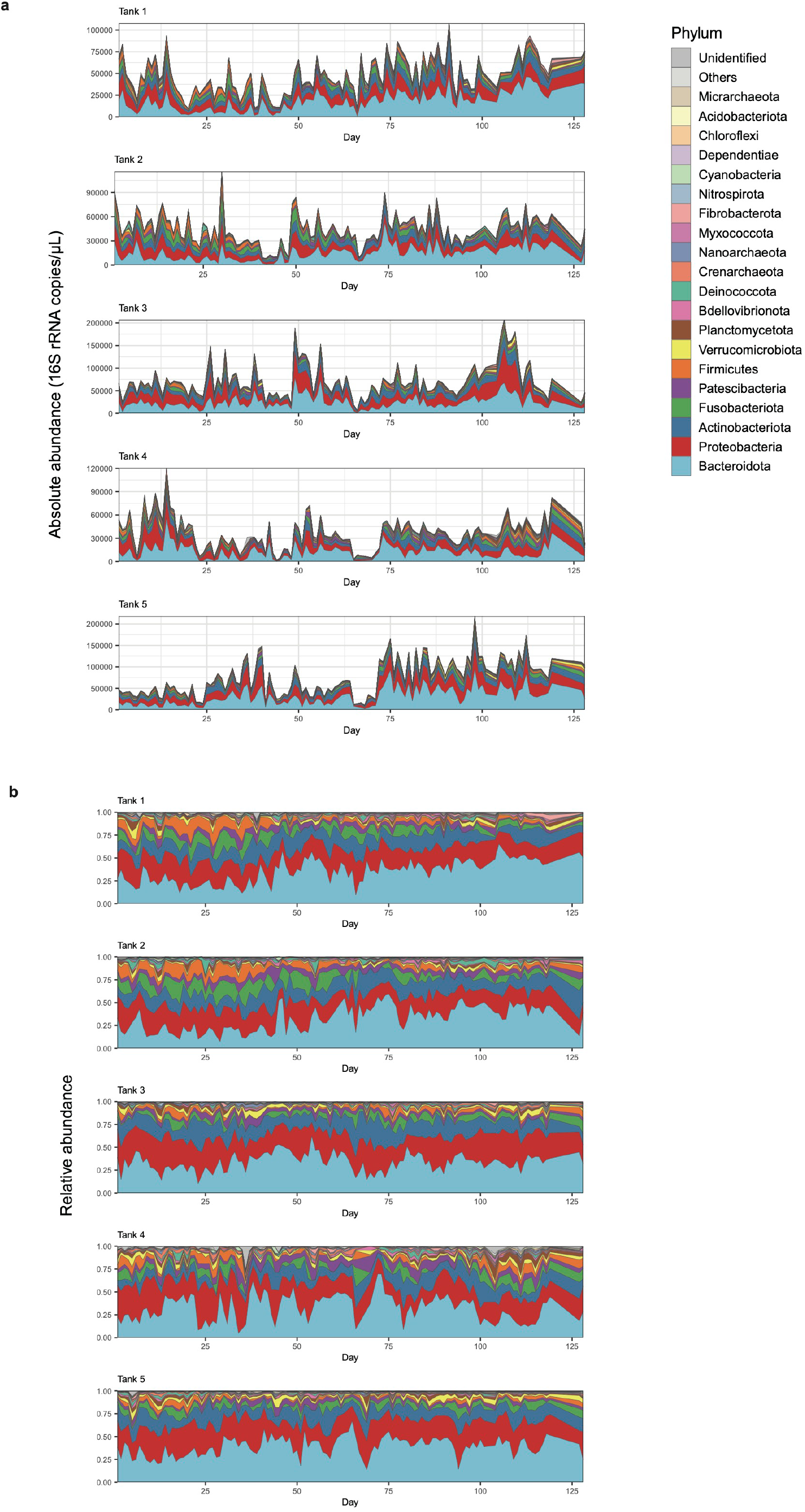
Phylum-level community structure. **a**, Dynamics of absolute abundance. For each water sample of each aquaculture tank, absolute abundance of prokaryotes was inferred as 16S rRNA gene copy concentration based on the quantitative amplicon sequencing approach with standard DNA gradients. **b**, Dynamics of relative abundance. The time-series of the phylum-level taxonomic compositions are shown for each aquaculture tank.

**Extended Data Fig. 2.**
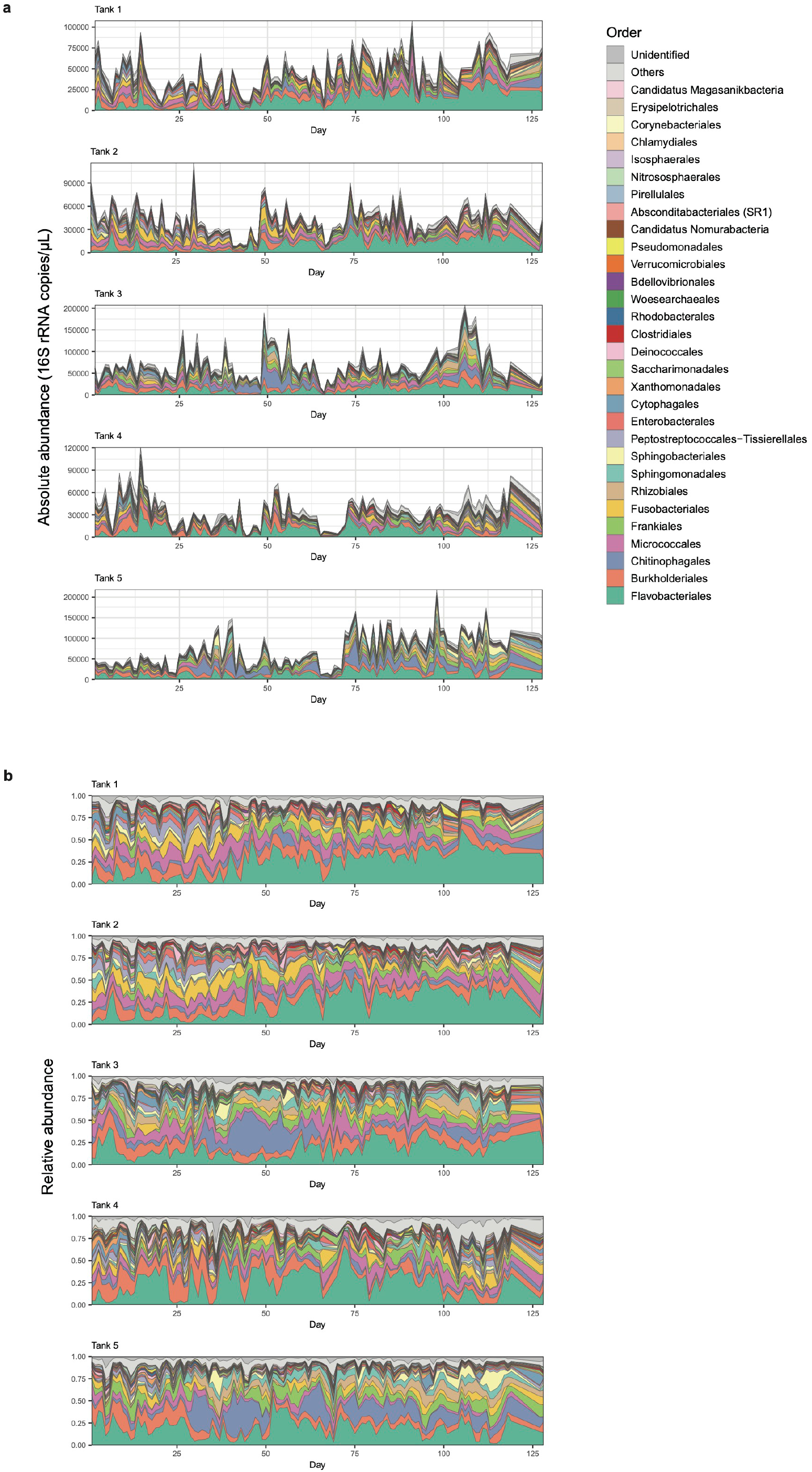
Order-level community structure. **a**, Dynamics of absolute abundance. For each water sample of each aquaculture tank, absolute abundance of prokaryotes was inferred as 16S rRNA gene copy concentration based on the quantitative amplicon sequencing approach with standard DNA gradients. **b**, Dynamics of relative abundance. The time-series of the order-level taxonomic compositions are shown for each aquaculture tank.

**Extended Data Fig. 3.**
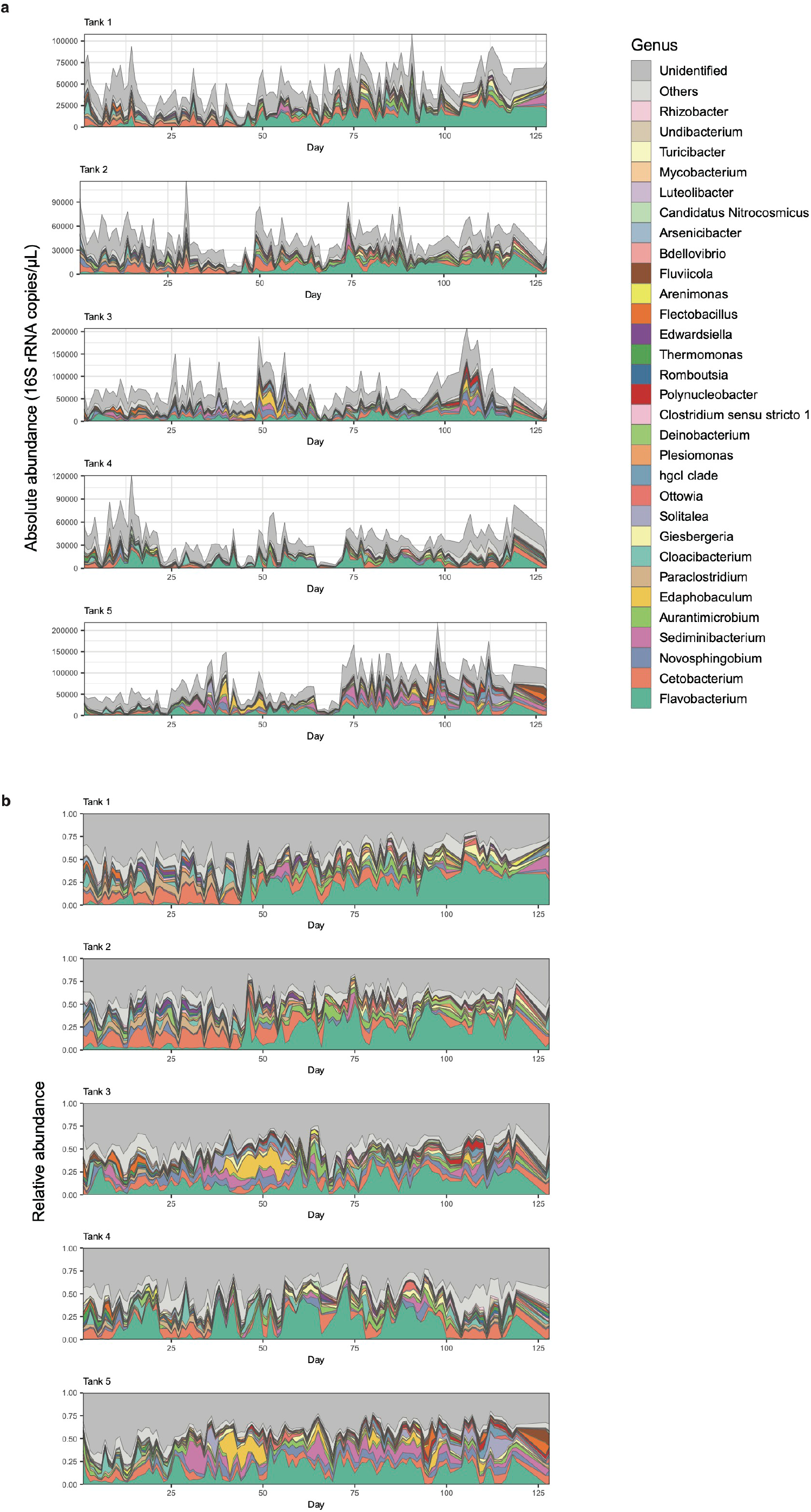
Genus-level community structure. **a**, Dynamics of absolute abundance. For each water sample of each aquaculture tank, absolute abundance of prokaryotes was inferred as 16S rRNA gene copy concentration based on the quantitative amplicon sequencing approach with standard DNA gradients. **b**, Dynamics of relative abundance. The time-series of the genus-level taxonomic compositions are shown for each aquaculture tank.

**Extended Data Fig. 4.**
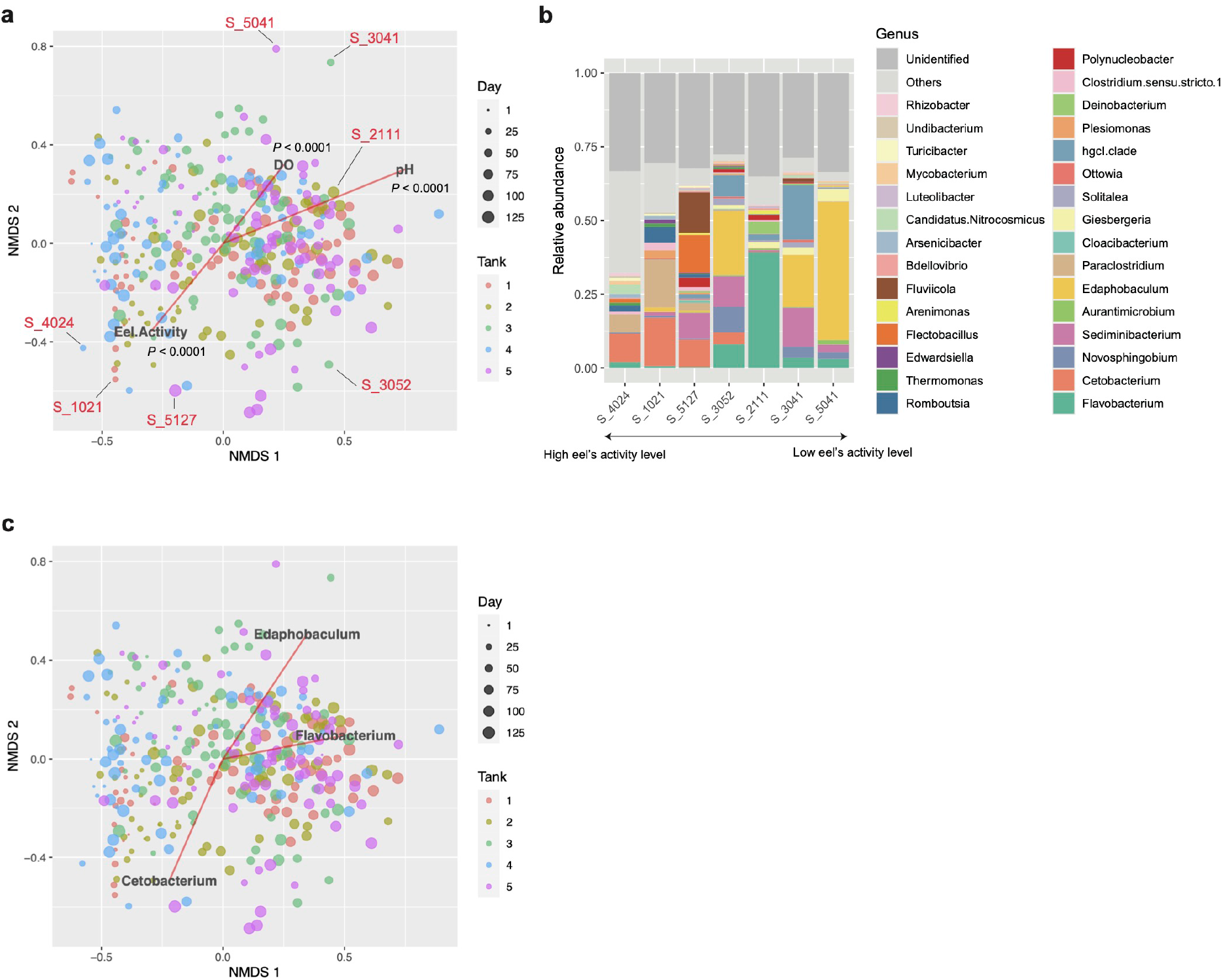
Multivariate analysis of community structure (genus level). **a**, Community state space. Community compositions of the samples are plotted on the two-dimensional surface defined with non-metric multidimensional scaling (NMDS). The NMDS was performed based on the Bray-Curtis -diversity of genus-level taxonomic compositions. The projections of the data points onto the vectors have maximum correlation with the variables examined (pH, DO, and eels’ activity level). Examples of community structure in the NMDS surface. For several points within the NMDS surface (panel **a)**, genus-level taxonomic compositions are shown. The example points are ordered along the vector representing high eels’ activity level. **c**, Indicator genera. The vectors representing the relative abundance of *Cetobacterium, Flavobacterium*, and *Edaphobaculum*, which were highlighted in the main text, are shown.

**Extended Data Fig. 5.**
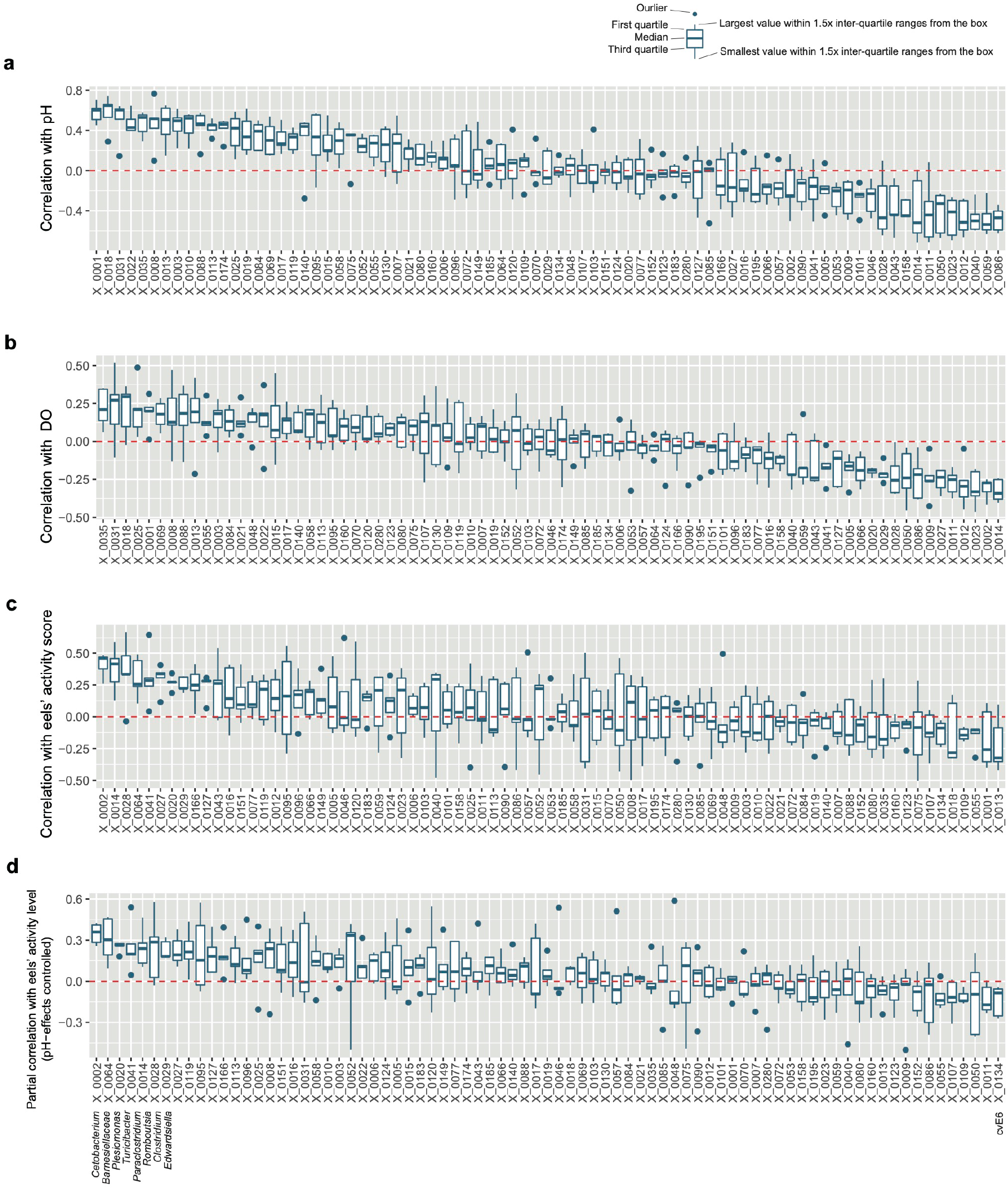
ASV-level comparison of correlation with environmental variables and eels’ activity level. **a**, Correlation with pH. Correlation with eels’ activity level is shown for the ASVs that appeared in all the aquaculture tanks (shown in the decreasing order of mean values). The boxes and bars represent variation across tanks. **b**, Correlation with DO. **c**, Correlation with eels’ activity level. **d**, Partial correlation with eels’ activity level (controlled variable = pH). Taxonomic information is shown for the ASVs discussed in the main text.

**Extended Data Fig. 6.**
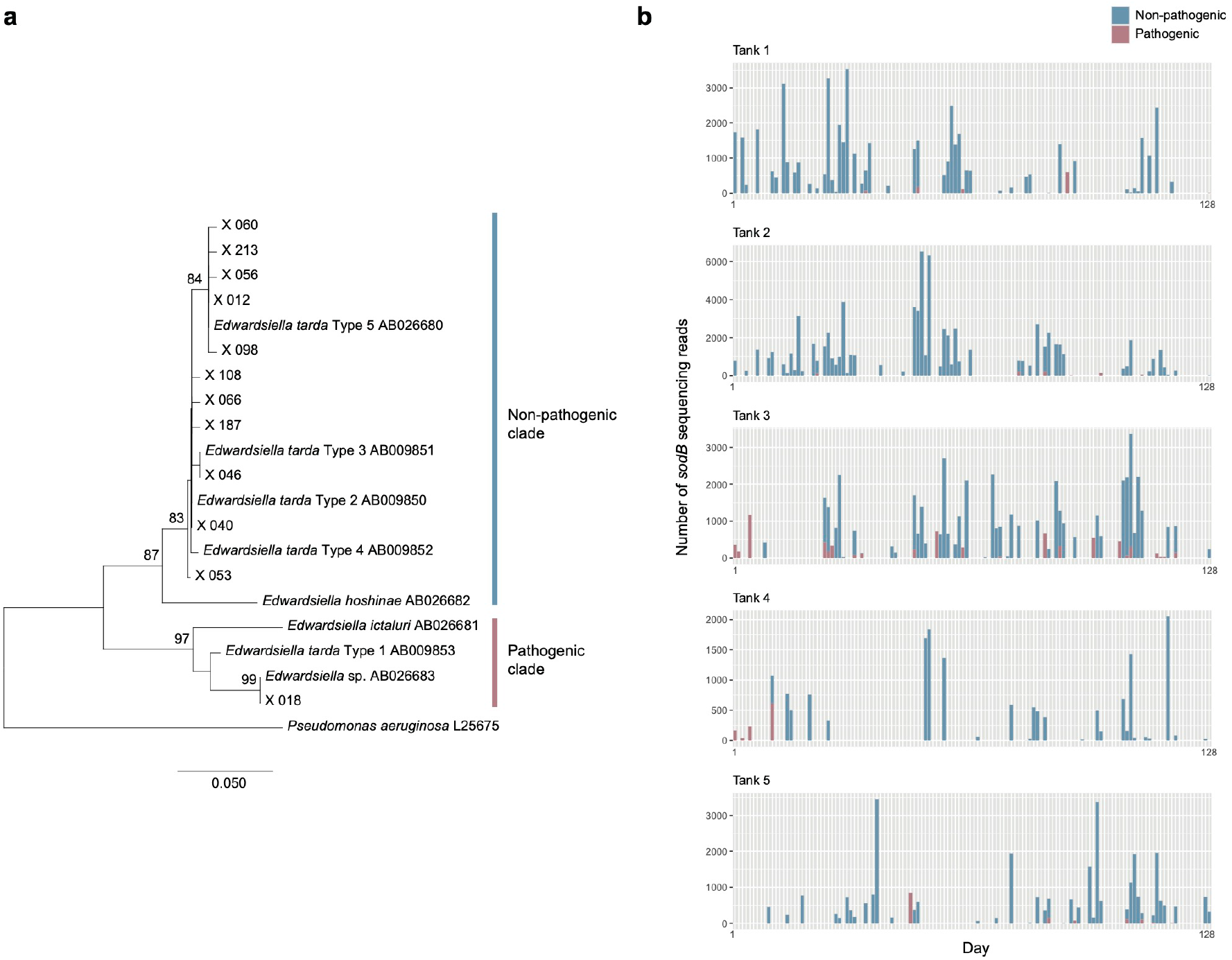
Phylogenetic analysis of *Edwardsiella*. **a**, Phylogeny of *Edwardsiella*. In an additional amplicon sequencing of the *sodB* gene, the neighbor-joining tree of the *Edwardsiella* bacteria was reconstructed with the maximum composite likelihood method. Bootstrap values larger than 70 % are shown on the nodes (1,000 permutations). The pathogenic and non-pathogenic clades identified in a previous study^29^ are indicated. **b**, Time-series of pathogenic and non-pathogenic *Edwardsiella*. The number of detected sequencing reads of the *sodB* fragment is across the time-series of each aquaculture tank.

**Extended Data Fig. 7.**
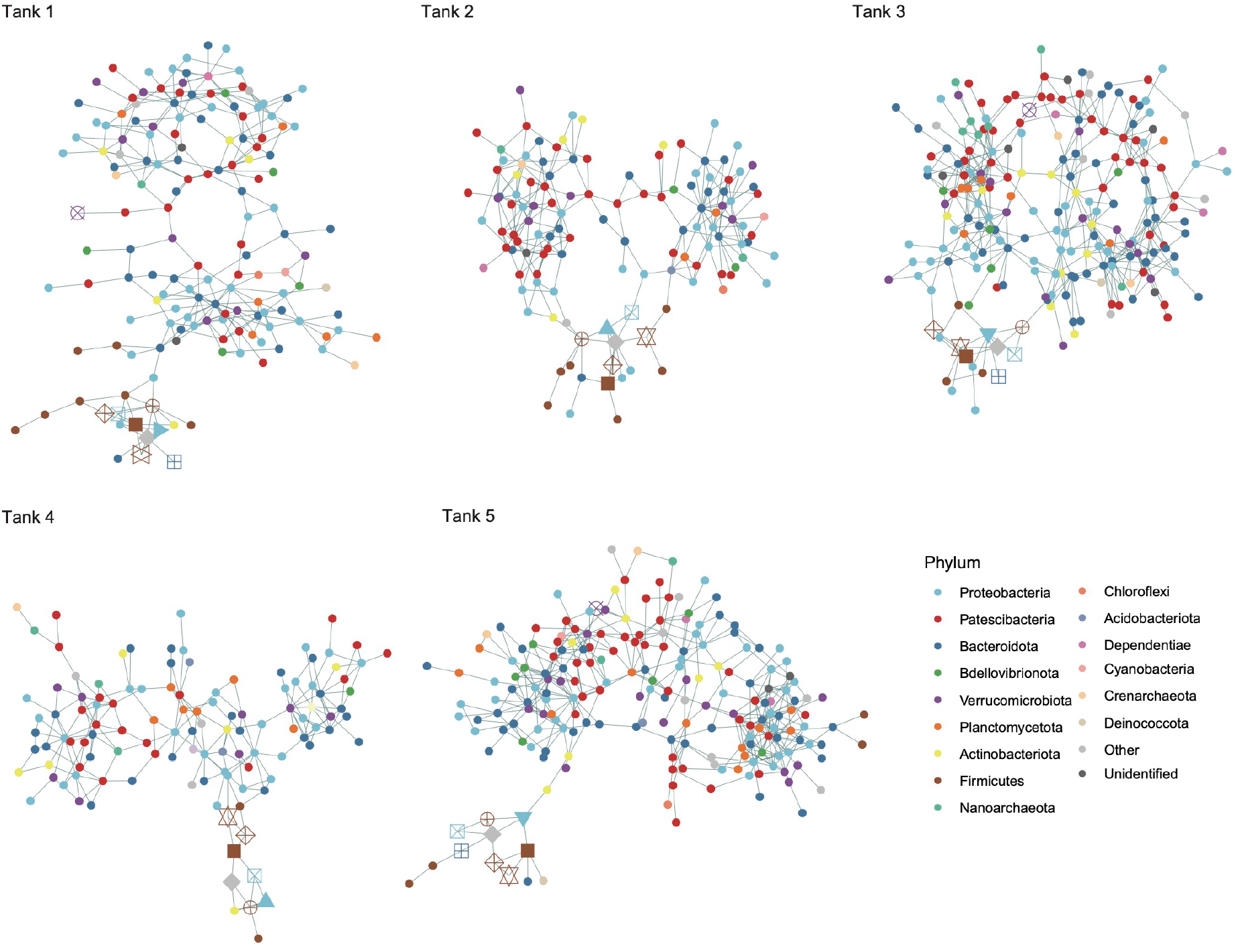
Taxonomy of the nodes within the coexistence networks. Within the coexistence networks shown in Figure 4, phylum-level taxonomy of the ASVs is shown. ASVs included in minor sub-networks (number of nodes < 5) are not shown. Only the ASVs that appeared in 30 or more samples were targeted in the analysis of each tank.

**Extended Data Fig. 8.**
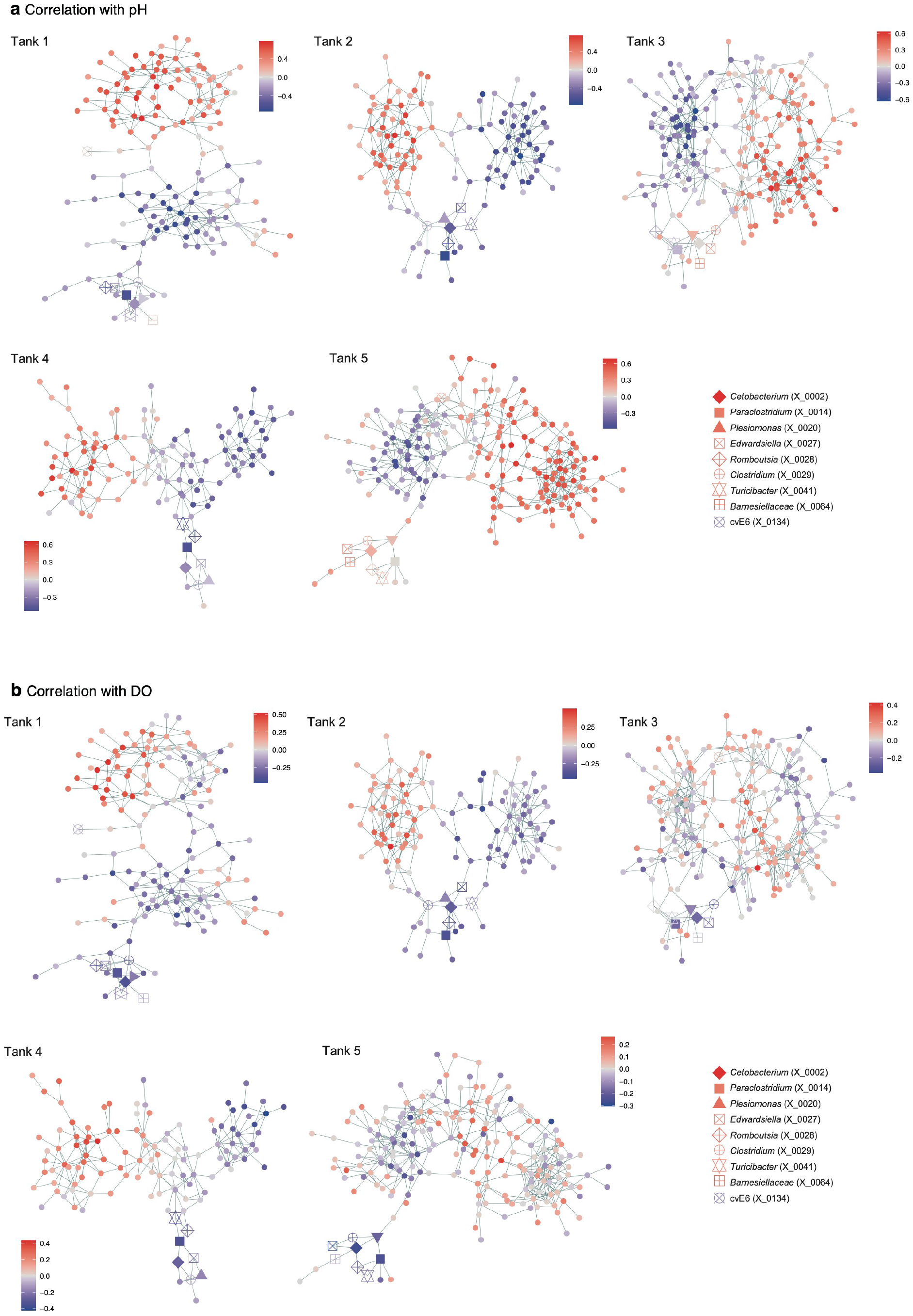
Correlations with environmental variables. **a**, Correlation with pH. For each microbial ASV included within the coexistence network of each aquaculture tank (Fig. 4), correlation between absolute abundance and pH is shown. ASVs included in minor sub-networks (number of nodes < 5) are not shown. Only the ASVs that appeared in 30 or more samples were targeted in the analysis of each tank. **b**, Correlation with dissolved oxygen level. For each microbial ASV included within the coexistence network of each aquaculture tank (Fig. 4), correlation between absolute abundance and dissolved oxygen (DO) level is shown.

**Extended Data Fig. 9.**
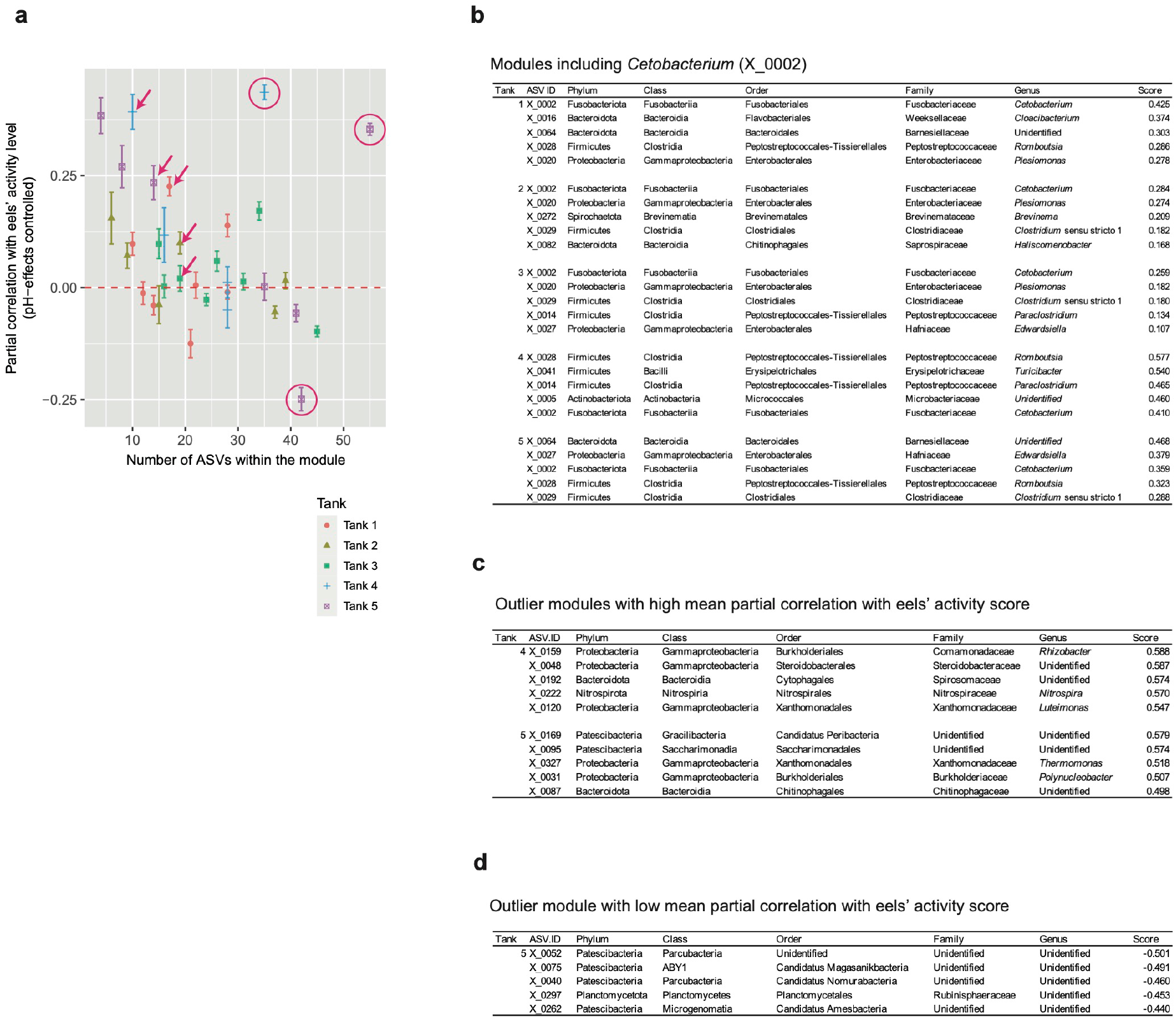
Properties of network modules. **a**, Module size and mean partial correlation with eels’ activity level. For each module within the coexistence network of each aquaculture tank (Fig. 4), the number of ASVs and mean partial correlation with eels’ activity level are shown. The modules including the *Cetobacterium* ASV (X_0002) is indicated by arrows. The outlier modules with large numbers of constituent ASVs and low/high mean partial correlation with eels’ activity level are highlighted by circles. **b**, Modules including the *Cetobacterium* ASV (X_0002). The top-five ASVs with the highest partial correlation with eels’ activity level are shown for each module. **c**, Outlier modules with high mean partial correlation with eels’ activity level. **d**, Outlier module with low mean partial correlation with eels’ activity level.

**Extended Data Fig. 10.**
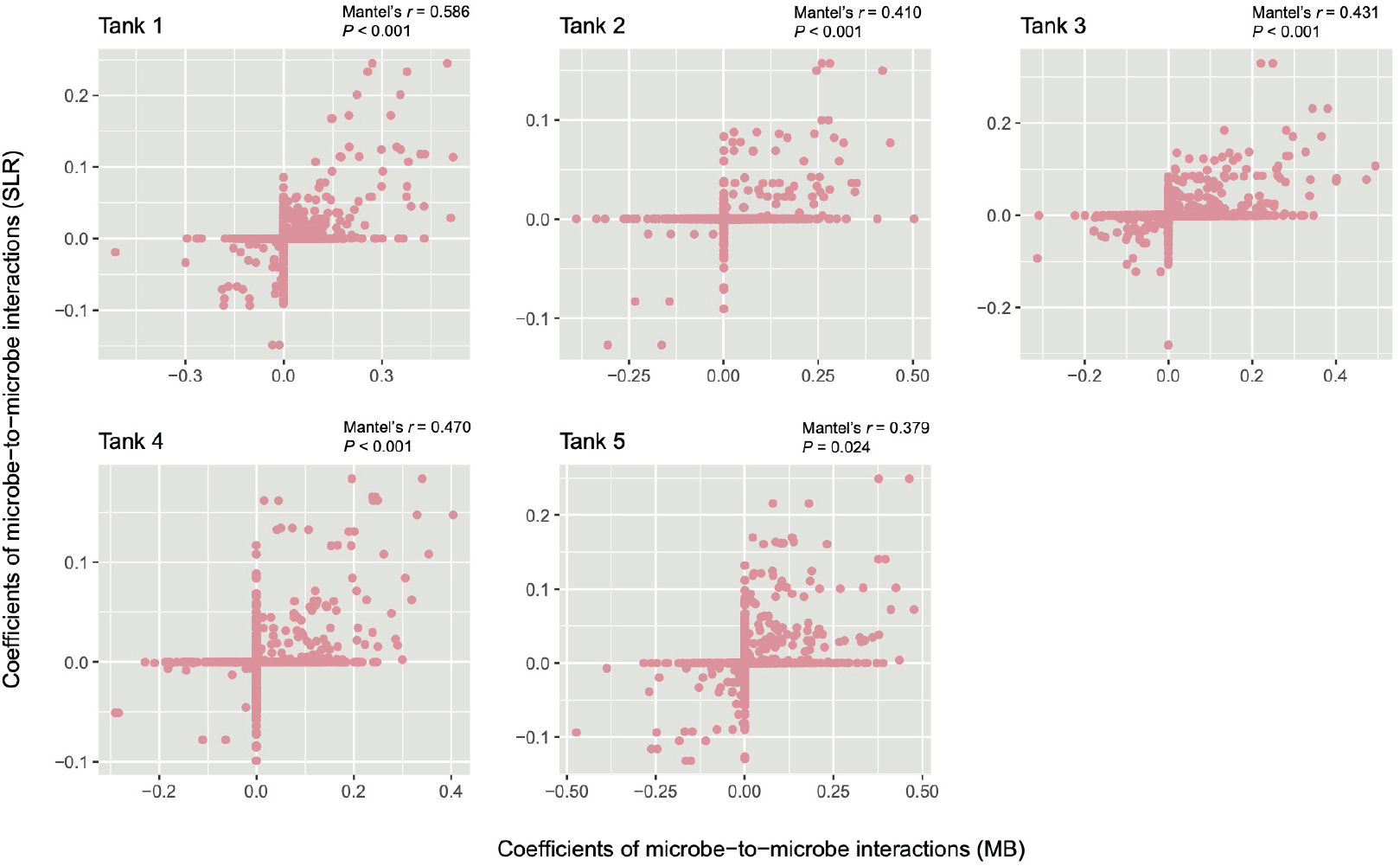
Comparison of network reconstruction methods. For each aquaculture tank, the network links inferred with the MB method was compared with those inferred with the SLR methods. The former is expected to represent interspecific interactions as well as potential sharing of environmental preference (i.e., niches) between nodes (ASVs). Meanwhile, the latter is expected to represent direct interactions between nodes. A positive/negative value indicates a potentially positive/negative interaction between a pair of microbial ASVs. The positive values (> 0) were used to draw networks of potential positive interactions between microbes as shown in Figure 5.

