## Supplementary Table for "Core species and interactions prominent in fish-associated microbiome dynamics"

**Supplementary Table 1 | Properties of the ASVs within the networks**. For the microbial ASVs that appeared in 30 or more samples in each aquaculture tank, coexistence patterns were inferred (Figure 4). For each ASV in each tank, the module it belonged to, taxonomy, partial correlation with eels' activity level (pH-effects controlled), and simple correlation with pH, DO, or eels' activity level are shown.

| Tank | Module | ASV.ID | Kingdom | Phylum | Class | Order | Family | Genus | Partial.Correlation.Eel | Correlation.pH | Correlation.DO | Correlation.Eel |
|---|---|---|---|---|---|---|---|---|---|---|---|---|
| 1 | 1 | X_0142 | Bacteria | Patescibacteria | ABY1 | Candidatus Magasanikbacteria | Unidentified | Unidentified | 0.293 | 0.190 | -0.120 | 0.164 |
| 1 | 1 | X_0106 | Bacteria | Bacteroidota | Bacteroidia | Flavobacteriales | Crocinitomicaceae | Fluviicola | 0.247 | -0.003 | -0.221 | 0.219 |
| 1 | 1 | X_0003 | Bacteria | Actinobacteriota | Actinobacteria | Frankiales | Sporichthyaceae | Unidentified | 0.191 | 0.613 | 0.183 | -0.156 |
| 1 | 1 | X_0185 | Bacteria | Bdellovibrionota | Bdellovibrionia | Bdellovibrionales | Bdellovibrionaceae | Bdellovibrio | 0.113 | 0.285 | 0.036 | -0.039 |
| 1 | 1 | X_0113 | Bacteria | Patescibacteria | Gracilibacteria | Candidatus Peribacteria | Unidentified | Unidentified | 0.109 | 0.404 | -0.009 | -0.102 |
| 1 | 1 | X_0015 | Bacteria | Bacteroidota | Bacteroidia | Chitinophagales | Chitinophagaceae | Unidentified | 0.101 | 0.549 | 0.449 | -0.185 |
| 1 | 1 | X_0010 | Bacteria | Actinobacteriota | Actinobacteria | Micrococcales | Microbacteriaceae | Aurantimicrobium | 0.099 | 0.545 | -0.007 | -0.184 |
| 1 | 1 | X_0084 | Archaea | Crenarchaeota | Nitrososphaeria | Nitrosopumilales | Nitrosopumilaceae | Candidatus Nitrosotenuis | 0.047 | 0.392 | 0.241 | -0.147 |
| 1 | 1 | X_0001 | Bacteria | Bacteroidota | Bacteroidia | Flavobacteriales | Flavobacteriaceae | Flavobacterium | 0.035 | 0.601 | 0.013 | -0.259 |
| 1 | 1 | X_0146 | Bacteria | Unidentified | Unidentified | Unidentified | Unidentified | Unidentified | 0.034 | 0.676 | 0.215 | -0.296 |
| 1 | 1 | X_0160 | Bacteria | Patescibacteria | Gracilibacteria | Candidatus Peregrinibacteria | Unidentified | Unidentified | -0.034 | 0.084 | -0.020 | -0.069 |
| 1 | 1 | X_0019 | Bacteria | Bacteroidota | Bacteroidia | Chitinophagales | Unidentified | Unidentified | -0.034 | 0.616 | 0.212 | -0.314 |
| 1 | 1 | X_0134 | Bacteria | Verrucomicrobiota | Chlamydiae | Chlamydiales | cvE6 | Unidentified | -0.043 | 0.032 | -0.095 | -0.053 |
| 1 | 1 | X_0144 | Bacteria | Bacteroidota | Bacteroidia | Unidentified | Unidentified | Unidentified | -0.047 | 0.326 | 0.103 | -0.193 |

| | | | | | | | | | | | | |
|---|---|---|---|---|---|---|---|---|---|---|---|---|
| 1 | 1 | X_0013 | Bacteria | Bacteroidota | Bacteroidia | Chitinophagales | Chitinophagaceae | Sediminibacterium | -0.069 | 0.649 | 0.419 | -0.352 |
| 1 | 1 | X_0386 | Bacteria | Patescibacteria | Saccharimonadia | Saccharimonadales | LWQ8 | Unidentified | -0.076 | 0.316 | 0.049 | -0.213 |
| 1 | 1 | X_0130 | Bacteria | Patescibacteria | Gracilibacteria | Unidentified | Unidentified | Unidentified | -0.080 | 0.259 | -0.170 | -0.190 |
| 1 | 1 | X_0033 | Bacteria | Proteobacteria | Gammaproteobacteria | Xanthomonadales | Xanthomonadaceae | Arenimonas | -0.083 | 0.755 | 0.287 | -0.404 |
| 1 | 1 | X_0055 | Bacteria | Patescibacteria | Microgenomatia | Candidatus Woesebacteria | Unidentified | Unidentified | -0.090 | 0.044 | 0.035 | -0.100 |
| 1 | 1 | X_0069 | Archaea | Nanoarchaeota | Nanoarchaeia | Woesearchaeales | GW2011_GWC1_47_15 | Unidentified | -0.162 | 0.438 | 0.284 | -0.335 |
| 1 | 1 | X_0017 | Bacteria | Bacteroidota | Bacteroidia | Chitinophagales | Chitinophagaceae | Edaphobaculum | -0.197 | 0.505 | 0.140 | -0.388 |
| 1 | 1 | X_0008 | Bacteria | Proteobacteria | Alphaproteobacteria | Rhizobiales | Rhizobiales Incertae Sedis | Unidentified | -0.240 | 0.765 | 0.471 | -0.497 |
| 1 | 2 | X_0002 | Bacteria | Fusobacteriota | Fusobacteriia | Fusobacteriales | Fusobacteriaceae | Cetobacterium | 0.425 | -0.265 | -0.417 | 0.486 |
| 1 | 2 | X_0016 | Bacteria | Bacteroidota | Bacteroidia | Flavobacteriales | Weeksellaceae | Cloacibacterium | 0.374 | -0.181 | -0.305 | 0.410 |
| 1 | 2 | X_0064 | Bacteria | Bacteroidota | Bacteroidia | Bacteroidales | Barnesiellaceae | Unidentified | 0.303 | 0.060 | -0.125 | 0.238 |
| 1 | 2 | X_0028 | Bacteria | Firmicutes | Clostridia | Peptostreptococcales-Tissierellales | Peptostreptococcaceae | Romboutsia | 0.286 | -0.581 | -0.346 | 0.479 |
| 1 | 2 | X_0020 | Bacteria | Proteobacteria | Gammaproteobacteria | Enterobacterales | Enterobacteriaceae | Plesiomonas | 0.278 | -0.061 | -0.241 | 0.273 |
| 1 | 2 | X_0005 | Bacteria | Actinobacteriota | Actinobacteria | Micrococcales | Microbacteriaceae | Unidentified | 0.260 | -0.182 | -0.335 | 0.312 |
| 1 | 2 | X_0014 | Bacteria | Firmicutes | Clostridia | Peptostreptococcales-Tissierellales | Peptostreptococcaceae | Paraclostridium | 0.239 | -0.620 | -0.379 | 0.458 |
| 1 | 2 | X_0232 | Bacteria | Firmicutes | Clostridia | Lachnospirales | Lachnospiraceae | Epulopiscium | 0.220 | -0.188 | -0.234 | 0.279 |
| 1 | 2 | X_0027 | Bacteria | Proteobacteria | Gammaproteobacteria | Enterobacterales | Hafniaceae | Edwardsiella | 0.192 | -0.308 | -0.192 | 0.307 |
| 1 | 2 | X_0041 | Bacteria | Firmicutes | Bacilli | Erysipelotrichales | Erysipelotrichaceae | Turicibacter | 0.192 | -0.157 | -0.173 | 0.241 |
| 1 | 2 | X_0127 | Bacteria | Firmicutes | Bacilli | Mycoplasmatales | Mycoplasmataceae | Unidentified | 0.182 | -0.275 | -0.307 | 0.284 |

| | | | | | | | | | | | | |
|---|---|---|---|---|---|---|---|---|---|---|---|---|
| 1 | 2 | X_0029 | Bacteria | Firmicutes | Clostridia | Clostridiales | Clostridiaceae | Clostridium sensu stricto 1 | 0.179 | -0.143 | -0.273 | 0.223 |
| 1 | 2 | X_0183 | Bacteria | Firmicutes | Clostridia | Clostridiales | Clostridiaceae | Unidentified | 0.169 | -0.061 | -0.087 | 0.177 |
| 1 | 2 | X_0166 | Bacteria | Firmicutes | Clostridia | Clostridiales | Clostridiaceae | Clostridium sensu stricto 1 | 0.157 | -0.243 | -0.078 | 0.249 |
| 1 | 2 | X_0203 | Bacteria | Firmicutes | Bacilli | Erysipelotrichales | Erysipelotrichaceae | Turicibacter | 0.152 | -0.204 | -0.127 | 0.227 |
| 1 | 2 | X_0136 | Bacteria | Proteobacteria | Alphaproteobacteria | Sphingomonadales | Sphingomonadaceae | Novosphingobium | 0.131 | -0.274 | -0.325 | 0.240 |
| 1 | 2 | X_0066 | Bacteria | Proteobacteria | Gammaproteobacteria | Enterobacterales | Enterobacteriaceae | Citrobacter | 0.100 | -0.234 | -0.257 | 0.197 |
| 1 | 3 | X_0088 | Bacteria | Patescibacteria | Parcubacteria | Candidatus Campbellbacteria | Unidentified | Unidentified | 0.175 | 0.577 | 0.185 | -0.146 |
| 1 | 3 | X_0018 | Bacteria | Proteobacteria | Gammaproteobacteria | Burkholderiales | Comamonadaceae | Giesbergeria | 0.111 | 0.741 | 0.363 | -0.284 |
| 1 | 3 | X_0007 | Bacteria | Proteobacteria | Alphaproteobacteria | Sphingomonadales | Sphingomonadaceae | Novosphingobium | 0.054 | 0.547 | 0.118 | -0.218 |
| 1 | 3 | X_0228 | Bacteria | Patescibacteria | WWE3 | Unidentified | Unidentified | Unidentified | 0.025 | 0.349 | 0.113 | -0.143 |
| 1 | 3 | X_0022 | Bacteria | Proteobacteria | Gammaproteobacteria | Burkholderiales | Comamonadaceae | Ottowia | 0.020 | 0.645 | 0.173 | -0.290 |
| 1 | 3 | X_0072 | Bacteria | Verrucomicrobiota | Verrucomicrobiae | Unidentified | Unidentified | Unidentified | -0.051 | 0.395 | 0.146 | -0.228 |
| 1 | 3 | X_0035 | Bacteria | Verrucomicrobiota | Verrucomicrobiae | Unidentified | Unidentified | Unidentified | -0.094 | 0.552 | 0.351 | -0.329 |
| 1 | 3 | X_0004 | Bacteria | Bacteroidota | Bacteroidia | Flavobacteriales | Flavobacteriaceae | Flavobacterium | -0.105 | 0.706 | 0.414 | -0.398 |
| 1 | 3 | X_0122 | Bacteria | Patescibacteria | Parcubacteria | Candidatus Nomurabacteria | Unidentified | Unidentified | -0.106 | 0.551 | 0.191 | -0.338 |
| 1 | 3 | X_0156 | Bacteria | Nitrospirota | Nitrospiria | Nitrospirales | Nitrospiraceae | Nitrospira | -0.124 | 0.466 | 0.245 | -0.316 |
| 1 | 3 | X_0051 | Bacteria | Bacteroidota | Bacteroidia | Flavobacteriales | Flavobacteriaceae | Flavobacterium | -0.131 | 0.568 | 0.291 | -0.363 |
| 1 | 3 | X_0031 | Bacteria | Proteobacteria | Gammaproteobacteria | Burkholderiales | Burkholderiaceae | Polynucleobacter | -0.156 | 0.641 | 0.517 | -0.407 |
| 1 | 3 | X_0111 | Bacteria | Verrucomicrobiota | Verrucomicrobiae | Verrucomicrobiales | Rubritaleaceae | Luteolibacter | -0.165 | 0.551 | 0.345 | -0.381 |
| 1 | 3 | X_0037 | Bacteria | Proteobacteria | Alphaproteobacteria | Sphingomonadales | Sphingomonadaceae | Novosphingobium | -0.193 | 0.401 | 0.398 | -0.345 |
| 1 | 3 | X_0025 | Bacteria | Actinobacteriota | Actinobacteria | Frankiales | Sporichthyaceae | hgcI clade | -0.206 | 0.516 | 0.487 | -0.398 |

| | | | | | | | | | | | | |
|---|---|---|---|---|---|---|---|---|---|---|---|---|
| 1 | 3 | X_0293 | Bacteria | Proteobacteria | Gammaproteobacteria | Diplorickettsiales | Diplorickettsiaceae | Aquicella | -0.210 | 0.470 | 0.316 | -0.385 |
| 1 | 3 | X_0094 | Bacteria | Bacteroidota | Bacteroidia | Flavobacteriales | Weeksellaceae | Cloacibacterium | -0.213 | 0.255 | 0.246 | -0.302 |
| 1 | 3 | X_0169 | Bacteria | Patescibacteria | Gracilibacteria | Candidatus Peribacteria | Unidentified | Unidentified | -0.240 | 0.527 | 0.324 | -0.428 |
| 1 | 3 | X_0056 | Bacteria | Verrucomicrobiota | Verrucomicrobiae | Unidentified | Unidentified | Unidentified | -0.308 | 0.637 | 0.482 | -0.510 |
| 1 | 3 | X_0145 | Bacteria | Planctomycetota | Planctomycetes | Isosphaerales | Isosphaeraceae | Isosphaera | -0.317 | 0.631 | 0.507 | -0.515 |
| 1 | 3 | X_0045 | Bacteria | Fibrobacterota | Fibrobacteria | Fibrobacterales | Fibrobacteraceae | Unidentified | -0.394 | 0.589 | 0.453 | -0.559 |
| 1 | 4 | X_0006 | Bacteria | Bacteroidota | Bacteroidia | Flavobacteriales | Weeksellaceae | Unidentified | 0.210 | 0.002 | -0.065 | 0.185 |
| 1 | 5 | X_0009 | Bacteria | Proteobacteria | Gammaproteobacteria | Burkholderiales | Comamonadaceae | Unidentified | 0.088 | -0.096 | -0.261 | 0.122 |
| 1 | 6 | X_0073 | Bacteria | Proteobacteria | Gammaproteobacteria | Burkholderiales | Comamonadaceae | Curvibacter | 0.171 | -0.448 | -0.344 | 0.346 |
| 1 | 6 | X_0050 | Bacteria | Patescibacteria | Gracilibacteria | Absconditabacteriales (SR1) | Unidentified | Unidentified | 0.063 | -0.621 | -0.240 | 0.337 |
| 1 | 6 | X_0195 | Bacteria | Bdellovibrionota | Bdellovibrionia | Bdellovibrionales | Bdellovibrionaceae | Bdellovibrio | 0.027 | -0.234 | -0.016 | 0.133 |
| 1 | 6 | X_0135 | Bacteria | Bacteroidota | Bacteroidia | Chitinophagales | 37-13 | Unidentified | 0.018 | -0.127 | -0.100 | 0.076 |
| 1 | 6 | X_0023 | Bacteria | Bacteroidota | Bacteroidia | Flavobacteriales | Flavobacteriaceae | Flavobacterium | 0.001 | -0.642 | -0.464 | 0.304 |
| 1 | 6 | X_0093 | Bacteria | Proteobacteria | Gammaproteobacteria | Xanthomonadales | Xanthomonadaceae | Thermomonas | -0.003 | -0.179 | -0.183 | 0.082 |
| 1 | 6 | X_0116 | Bacteria | Bacteroidota | Bacteroidia | Chitinophagales | Chitinophagaceae | Edaphobaculum | -0.018 | -0.634 | -0.235 | 0.287 |
| 1 | 6 | X_0121 | Bacteria | Bacteroidota | Bacteroidia | Chitinophagales | Chitinophagaceae | Parasediminibacterium | -0.027 | -0.451 | -0.278 | 0.191 |
| 1 | 6 | X_0012 | Bacteria | Proteobacteria | Gammaproteobacteria | Burkholderiales | Neisseriaceae | Unidentified | -0.032 | -0.662 | -0.296 | 0.291 |
| 1 | 6 | X_0132 | Bacteria | Patescibacteria | Parcubacteria | Candidatus Campbellbacteria | Unidentified | Unidentified | -0.041 | -0.331 | -0.277 | 0.122 |
| 1 | 6 | X_0123 | Bacteria | Verrucomicrobiota | Chlamydiae | Chlamydiales | cvE6 | Unidentified | -0.137 | -0.064 | 0.117 | -0.091 |
| 1 | 6 | X_0011 | Bacteria | Actinobacteriota | Actinobacteria | Micrococcales | Microbacteriaceae | Unidentified | -0.171 | -0.659 | -0.331 | 0.197 |
| 1 | 7 | X_0101 | Bacteria | Bacteroidota | Bacteroidia | Cytophagales | Microscillaceae | Unidentified | 0.098 | -0.238 | -0.221 | 0.196 |

| | | | | | | | | | | | | |
|---|---|---|---|---|---|---|---|---|---|---|---|---|
| 1 | 7 | X_0099 | Bacteria | Planctomycetota | Planctomycetes | Pirellulales | Pirellulaceae | Unidentified | 0.049 | 0.360 | 0.164 | -0.130 |
| 1 | 7 | X_0021 | Bacteria | Deinococcota | Deinococci | Deinococcales | Deinococcaceae | Deinobacterium | 0.034 | 0.219 | 0.133 | -0.074 |
| 1 | 7 | X_0240 | Bacteria | Proteobacteria | Alphaproteobacteria | Rhizobiales | Rhizobiales Incertae Sedis | Unidentified | 0.016 | -0.505 | -0.187 | 0.250 |
| 1 | 7 | X_0062 | Bacteria | Planctomycetota | Planctomycetes | Isosphaerales | Isosphaeraceae | Tundrisphaera | 0.003 | -0.329 | -0.069 | 0.157 |
| 1 | 7 | X_0085 | Bacteria | Proteobacteria | Gammaproteobacteria | Pseudomonadales | Pseudomonadaceae | Pseudomonas | -0.007 | 0.077 | 0.120 | -0.043 |
| 1 | 7 | X_0083 | Bacteria | Planctomycetota | Planctomycetes | Isosphaerales | Isosphaeraceae | Unidentified | -0.015 | 0.012 | -0.002 | -0.019 |
| 1 | 7 | X_0034 | Archaea | Crenarchaeota | Nitrososphaeria | Nitrososphaerales | Nitrososphaeraceae | Candidatus Nitrocosmicus | -0.047 | 0.250 | 0.252 | -0.158 |
| 1 | 7 | X_0046 | Bacteria | Proteobacteria | Gammaproteobacteria | Burkholderiales | Comamonadaceae | Rhizobacter | -0.053 | -0.079 | 0.073 | -0.009 |
| 1 | 7 | X_0173 | Bacteria | Proteobacteria | Alphaproteobacteria | Acetobacterales | Acetobacteraceae | Roseomonas | -0.077 | -0.224 | -0.091 | 0.039 |
| 1 | 7 | X_0275 | Bacteria | Chloroflexi | Chloroflexia | Thermomicrobiales | JG30-KF-CM45 | Unidentified | -0.095 | -0.026 | 0.104 | -0.072 |
| 1 | 7 | X_0172 | Bacteria | Proteobacteria | Alphaproteobacteria | Acetobacterales | Acetobacteraceae | Rhodovastum | -0.110 | 0.164 | 0.153 | -0.173 |
| 1 | 7 | X_0077 | Bacteria | Bacteroidota | Bacteroidia | Chitinophagales | Chitinophagaceae | Taibaiella | -0.152 | -0.232 | -0.284 | -0.020 |
| 1 | 7 | X_0048 | Bacteria | Proteobacteria | Gammaproteobacteria | Steroidobacterales | Steroidobacteraceae | Unidentified | -0.196 | 0.161 | 0.219 | -0.246 |
| 1 | 8 | X_0052 | Bacteria | Patescibacteria | Parcubacteria | Unidentified | Unidentified | Unidentified | 0.355 | 0.184 | -0.316 | 0.221 |
| 1 | 8 | X_0388 | Bacteria | Proteobacteria | Gammaproteobacteria | Unidentified | Unidentified | Unidentified | 0.323 | 0.200 | -0.090 | 0.185 |
| 1 | 8 | X_0067 | Bacteria | Proteobacteria | Gammaproteobacteria | Burkholderiales | Comamonadaceae | Unidentified | 0.295 | 0.287 | -0.022 | 0.114 |
| 1 | 8 | X_0140 | Bacteria | Bacteroidota | Bacteroidia | Chitinophagales | Saprospiraceae | Unidentified | 0.269 | 0.468 | 0.190 | -0.011 |
| 1 | 8 | X_0078 | Bacteria | Proteobacteria | Gammaproteobacteria | Alteromonadales | Alteromonadaceae | Rheinheimera | 0.256 | 0.429 | 0.050 | 0.002 |
| 1 | 8 | X_0075 | Bacteria | Patescibacteria | ABY1 | Candidatus Magasanikbacteria | Unidentified | Unidentified | 0.243 | 0.347 | -0.062 | 0.037 |
| 1 | 8 | X_0253 | Bacteria | Proteobacteria | Gammaproteobacteria | Unidentified | Unidentified | Unidentified | 0.238 | 0.245 | -0.127 | 0.088 |
| 1 | 8 | X_0076 | Bacteria | Bdellovibrionota | Bdellovibrionia | Bacteriovoracales | Bacteriovoracaceae | Bacteriovorax | 0.228 | 0.417 | 0.044 | -0.014 |

| | | | | | | | | | | | | |
|---|---|---|---|---|---|---|---|---|---|---|---|---|
| 1 | 8 | X_0082 | Bacteria | Bacteroidota | Bacteroidia | Chitinophagales | Saprospiraceae | Haliscomenobacter | 0.224 | 0.446 | -0.050 | -0.034 |
| 1 | 8 | X_0287 | Bacteria | Proteobacteria | Alphaproteobacteria | Micavibrionales | Unidentified | Unidentified | 0.217 | 0.373 | -0.064 | 0.002 |
| 1 | 8 | X_0174 | Bacteria | Patescibacteria | Saccharimonadia | Saccharimonadales | Unidentified | Unidentified | 0.214 | 0.238 | -0.302 | 0.071 |
| 1 | 8 | X_0026 | Bacteria | Bacteroidota | Bacteroidia | Flavobacteriales | Flavobacteriaceae | Flavobacterium | 0.205 | 0.055 | -0.280 | 0.155 |
| 1 | 8 | X_0196 | Bacteria | Proteobacteria | Gammaproteobacteria | Xanthomonadales | Xanthomonadaceae | Thermomonas | 0.196 | 0.004 | -0.334 | 0.171 |
| 1 | 8 | X_0192 | Bacteria | Bacteroidota | Bacteroidia | Cytophagales | Spirosomaceae | Unidentified | 0.184 | 0.231 | -0.214 | 0.049 |
| 1 | 8 | X_0119 | Bacteria | Patescibacteria | Parcubacteria | Candidatus Zambryskibacteria | Unidentified | Unidentified | 0.163 | 0.334 | -0.208 | -0.022 |
| 1 | 8 | X_0058 | Bacteria | Patescibacteria | Parcubacteria | Candidatus Nomurabacteria | Unidentified | Unidentified | 0.145 | 0.413 | 0.181 | -0.078 |
| 1 | 8 | X_0129 | Bacteria | Bacteroidota | Bacteroidia | Cytophagales | Spirosomaceae | Runella | 0.099 | 0.306 | -0.041 | -0.061 |
| 1 | 8 | X_0133 | Bacteria | Proteobacteria | Alphaproteobacteria | Rhodobacterales | Rhodobacteraceae | Tabrizicola | 0.077 | 0.439 | 0.140 | -0.145 |
| 1 | 8 | X_0153 | Bacteria | Proteobacteria | Gammaproteobacteria | Burkholderiales | Comamonadaceae | Inhella | 0.072 | 0.188 | 0.070 | -0.027 |
| 1 | 8 | X_0061 | Bacteria | Myxococcota | Polyangia | Haliangiales | Haliangiaceae | Haliangium | 0.064 | 0.500 | 0.168 | -0.187 |
| 1 | 8 | X_0171 | Bacteria | Planctomycetota | Planctomycetes | Planctomycetales | Rubinisphaeraceae | Rubinisphaera | 0.059 | 0.418 | 0.141 | -0.150 |
| 1 | 8 | X_0233 | Bacteria | Proteobacteria | Alphaproteobacteria | Sphingomonadales | Sphingomonadaceae | Sphingopyxis | 0.058 | 0.182 | -0.021 | -0.035 |
| 1 | 8 | X_0415 | Bacteria | Proteobacteria | Gammaproteobacteria | Burkholderiales | Comamonadaceae | Unidentified | 0.054 | 0.547 | 0.327 | -0.218 |
| 1 | 8 | X_0089 | Bacteria | Bacteroidota | Kapabacteria | Kapabacteriales | Unidentified | Unidentified | 0.017 | 0.631 | 0.172 | -0.286 |
| 1 | 8 | X_0176 | Bacteria | Verrucomicrobiota | Verrucomicrobiae | Pedosphaerales | Pedosphaeraceae | SH3-11 | -0.028 | 0.485 | 0.115 | -0.251 |
| 1 | 8 | X_0255 | Bacteria | Spirochaetota | Leptospirae | Leptospirales | Leptospiraceae | Turneriella | -0.078 | 0.486 | 0.133 | -0.289 |
| 1 | 8 | X_0496 | Bacteria | Bacteroidota | Bacteroidia | Cytophagales | Cyclobacteriaceae | Unidentified | -0.131 | 0.593 | 0.364 | -0.372 |
| 1 | 8 | X_0081 | Bacteria | Patescibacteria | Saccharimonadia | Saccharimonadales | Unidentified | Unidentified | -0.139 | 0.418 | 0.041 | -0.309 |
| 1 | 9 | X_0237 | Bacteria | Proteobacteria | Gammaproteobacteria | Diplorickettsiales | Diplorickettsiaceae | Unidentified | 0.166 | -0.187 | -0.211 | 0.232 |

| | | | | | | | | | | | | |
|---|---|---|---|---|---|---|---|---|---|---|---|---|
| 1 | 9 | X_0158 | Bacteria | Bdellovibrionota | Oligoflexia | 0319-6G20 | Unidentified | Unidentified | 0.121 | -0.283 | -0.103 | 0.235 |
| 1 | 9 | X_0065 | Bacteria | Patescibacteria | Saccharimonadia | Saccharimonadales | Unidentified | Unidentified | 0.077 | -0.632 | -0.467 | 0.350 |
| 1 | 9 | X_0044 | Bacteria | Unidentified | Unidentified | Unidentified | Unidentified | Unidentified | 0.072 | -0.302 | -0.119 | 0.203 |
| 1 | 9 | X_0348 | Bacteria | Patescibacteria | Parcubacteria | Unidentified | Unidentified | Unidentified | 0.069 | -0.180 | -0.196 | 0.145 |
| 1 | 9 | X_0043 | Bacteria | Proteobacteria | Gammaproteobacteria | Burkholderiales | Oxalobacteraceae | Undibacterium | 0.068 | -0.443 | -0.265 | 0.263 |
| 1 | 9 | X_0333 | Bacteria | Proteobacteria | Gammaproteobacteria | Xanthomonadales | Rhodanobacteraceae | Dokdonella | 0.067 | -0.383 | -0.382 | 0.235 |
| 1 | 9 | X_0155 | Bacteria | Proteobacteria | Alphaproteobacteria | Rhizobiales | Rhizobiaceae | Unidentified | 0.060 | -0.281 | -0.133 | 0.183 |
| 1 | 9 | X_0036 | Bacteria | Bacteroidota | Bacteroidia | Cytophagales | Spirosomaceae | Arsenicibacter | 0.041 | -0.692 | -0.343 | 0.352 |
| 1 | 9 | X_0193 | Bacteria | Proteobacteria | Alphaproteobacteria | Rhodobacterales | Rhodobacteraceae | Defluviimonas | 0.030 | -0.298 | -0.111 | 0.166 |
| 1 | 9 | X_0040 | Bacteria | Patescibacteria | Parcubacteria | Candidatus Nomurabacteria | Unidentified | Unidentified | 0.019 | -0.596 | -0.317 | 0.295 |
| 1 | 9 | X_0184 | Bacteria | Proteobacteria | Gammaproteobacteria | Oceanospirillales | Unidentified | Unidentified | -0.002 | -0.269 | -0.040 | 0.126 |
| 1 | 9 | X_0091 | Bacteria | Bacteroidota | Bacteroidia | Chitinophagales | Chitinophagaceae | Sediminibacterium | -0.010 | -0.734 | -0.268 | 0.340 |
| 1 | 9 | X_0071 | Bacteria | Proteobacteria | Gammaproteobacteria | Xanthomonadales | Xanthomonadaceae | Thermomonas | -0.013 | -0.622 | -0.292 | 0.284 |
| 1 | 9 | X_0346 | Bacteria | Firmicutes | Bacilli | Lactobacillales | Streptococcaceae | Lactococcus | -0.015 | -0.044 | 0.103 | 0.007 |
| 1 | 9 | X_0060 | Bacteria | Verrucomicrobiota | Verrucomicrobiae | Unidentified | Unidentified | Unidentified | -0.020 | -0.703 | -0.184 | 0.319 |
| 1 | 9 | X_0181 | Bacteria | Proteobacteria | Alphaproteobacteria | Sphingomonadales | Sphingomonadaceae | Novosphingobium | -0.029 | -0.458 | -0.356 | 0.193 |
| 1 | 9 | X_0039 | Bacteria | Bacteroidota | Bacteroidia | Sphingobacteriales | env.OPS 17 | Unidentified | -0.036 | -0.725 | -0.241 | 0.320 |
| 1 | 9 | X_0575 | Bacteria | Proteobacteria | Alphaproteobacteria | Sphingomonadales | Sphingomonadaceae | Unidentified | -0.038 | -0.416 | -0.212 | 0.165 |
| 1 | 9 | X_0108 | Bacteria | Proteobacteria | Gammaproteobacteria | Burkholderiales | Comamonadaceae | Ottowia | -0.050 | -0.646 | -0.248 | 0.271 |
| 1 | 9 | X_0343 | Bacteria | Patescibacteria | Saccharimonadia | Saccharimonadales | Unidentified | Unidentified | -0.062 | -0.599 | -0.197 | 0.238 |
| 1 | 9 | X_0149 | Bacteria | Bacteroidota | Kapabacteria | Kapabacteriales | Unidentified | Unidentified | -0.064 | -0.179 | -0.163 | 0.029 |

| | | | | | | | | | | | | |
|---|---|---|---|---|---|---|---|---|---|---|---|---|
| 1 | 9 | X_0226 | Bacteria | Firmicutes | Clostridia | Clostridiales | Clostridiaceae | Clostridium sensu stricto 1 | -0.071 | -0.064 | 0.057 | -0.032 |
| 1 | 9 | X_0059 | Bacteria | Verrucomicrobiota | Verrucomicrobiae | Verrucomicrobiales | Rubritaleaceae | Luteolibacter | -0.100 | -0.590 | -0.235 | 0.207 |
| 1 | 9 | X_0189 | Bacteria | Planctomycetota | Planctomycetes | Pirellulales | Pirellulaceae | Pirellula | -0.117 | -0.243 | -0.173 | 0.015 |
| 1 | 9 | X_0070 | Bacteria | Proteobacteria | Gammaproteobacteria | Burkholderiales | Chromobacteriaceae | Vogesella | -0.120 | -0.012 | 0.173 | -0.100 |
| 1 | 9 | X_0112 | Bacteria | Bacteroidota | Bacteroidia | Chitinophagales | Saprospiraceae | Unidentified | -0.155 | -0.438 | -0.217 | 0.084 |
| 1 | 9 | X_0030 | Bacteria | Bacteroidota | Bacteroidia | Cytophagales | Spirosomaceae | Flectobacillus | -0.177 | -0.380 | -0.087 | 0.034 |
| 1 | 10 | X_0032 | Bacteria | Bacteroidota | Bacteroidia | Sphingobacteriales | env.OPS 17 | Unidentified | 0.243 | -0.222 | -0.089 | 0.313 |
| 1 | 10 | X_0199 | Bacteria | Proteobacteria | Alphaproteobacteria | Rhodobacterales | Rhodobacteraceae | Unidentified | 0.167 | 0.108 | -0.169 | 0.096 |
| 1 | 10 | X_0086 | Bacteria | Patescibacteria | Saccharimonadia | Saccharimonadales | Unidentified | Unidentified | 0.137 | -0.623 | -0.381 | 0.388 |
| 1 | 10 | X_0053 | Bacteria | Bacteroidota | Bacteroidia | Chitinophagales | Chitinophagaceae | Unidentified | 0.117 | -0.443 | -0.325 | 0.301 |
| 1 | 10 | X_0147 | Bacteria | Verrucomicrobiota | Verrucomicrobiae | Unidentified | Unidentified | Unidentified | 0.111 | -0.326 | -0.282 | 0.247 |
| 1 | 10 | X_0507 | Bacteria | Bdellovibrionota | Oligoflexia | 0319-6G20 | Unidentified | Unidentified | 0.103 | -0.100 | 0.134 | 0.138 |
| 1 | 10 | X_0115 | Bacteria | Patescibacteria | Gracilibacteria | Absconditabacteriales (SR1) | Unidentified | Unidentified | 0.097 | -0.220 | -0.328 | 0.187 |
| 1 | 10 | X_0214 | Bacteria | Proteobacteria | Alphaproteobacteria | Sphingomonadales | Sphingomonadaceae | Unidentified | 0.041 | 0.063 | -0.165 | 0.007 |
| 1 | 10 | X_0154 | Bacteria | Bacteroidota | Bacteroidia | Flavobacteriales | Crocinitomicaceae | Fluviicola | -0.008 | -0.081 | -0.078 | 0.031 |
| 1 | 10 | X_0090 | Bacteria | Cyanobacteria | Vampirivibrionia | Obscuribacterales | Obscuribacteraceae | Candidatus Obscuribacter | -0.031 | -0.090 | 0.056 | 0.015 |
| 1 | 11 | X_0057 | Bacteria | Proteobacteria | Alphaproteobacteria | Rhodobacterales | Rhodobacteraceae | Gemmobacter | -0.062 | 0.112 | 0.043 | -0.107 |
| 1 | 12 | X_0068 | Bacteria | Patescibacteria | Saccharimonadia | Saccharimonadales | Saccharimonadaceae | TM7a | 0.166 | 0.139 | -0.094 | 0.080 |
| 1 | 13 | X_0109 | Bacteria | Proteobacteria | Gammaproteobacteria | Pseudomonadales | Moraxellaceae | Acinetobacter | -0.145 | 0.037 | 0.282 | -0.146 |
| 1 | 13 | X_0080 | Bacteria | Proteobacteria | Gammaproteobacteria | Pseudomonadales | Moraxellaceae | Acinetobacter | -0.303 | 0.122 | 0.158 | -0.323 |
| 1 | 13 | X_0107 | Bacteria | Proteobacteria | Gammaproteobacteria | Aeromonadales | Aeromonadaceae | Aeromonas | -0.310 | 0.211 | 0.303 | -0.367 |

| | | | | | | | | | | | | |
|---|---|---|---|---|---|---|---|---|---|---|---|---|
| 1 | 14 | X_0095 | Bacteria | Patescibacteria | Saccharimonadia | Saccharimonadales | Unidentified | Unidentified | -0.074 | 0.154 | 0.273 | -0.137 |
| 1 | 15 | X_0096 | Bacteria | Bacteroidota | Bacteroidia | Cytophagales | Spirosomaceae | Emticicia | 0.015 | 0.313 | -0.001 | -0.135 |
| 1 | 16 | X_0103 | Bacteria | Bacteroidota | Bacteroidia | Flavobacteriales | Weeksellaceae | Bergeyella | -0.158 | 0.057 | 0.082 | -0.166 |
| 1 | 17 | X_0120 | Bacteria | Proteobacteria | Gammaproteobacteria | Xanthomonadales | Xanthomonadaceae | Luteimonas | 0.015 | 0.076 | 0.238 | -0.023 |
| 1 | 18 | X_0124 | Bacteria | Proteobacteria | Gammaproteobacteria | Burkholderiales | Leeiaceae | Leeia | 0.211 | 0.081 | -0.293 | 0.148 |
| 1 | 19 | X_0126 | Bacteria | Nitrospirota | Nitrospiria | Nitrospirales | Nitrospiraceae | Nitrospira | -0.178 | 0.340 | 0.277 | -0.308 |
| 1 | 20 | X_0131 | Bacteria | Proteobacteria | Gammaproteobacteria | Xanthomonadales | Xanthomonadaceae | Thermomonas | 0.058 | -0.091 | -0.215 | 0.094 |
| 1 | 21 | X_0151 | Bacteria | Proteobacteria | Alphaproteobacteria | Rhizobiales | Rhizobiales Incertae Sedis | Phreatobacter | 0.270 | 0.012 | -0.200 | 0.232 |
| 1 | 22 | X_0152 | Bacteria | Proteobacteria | Alphaproteobacteria | Rhizobiales | Beijerinckiaceae | Methylobacterium-Methylorubrum | -0.076 | -0.092 | 0.075 | -0.023 |
| 1 | 23 | X_0186 | Bacteria | Proteobacteria | Gammaproteobacteria | Burkholderiales | Oxalobacteraceae | Undibacterium | 0.111 | -0.067 | -0.010 | 0.129 |
| 1 | 24 | X_0188 | Bacteria | Patescibacteria | Unidentified | Unidentified | Unidentified | Unidentified | -0.103 | 0.022 | 0.066 | -0.101 |
| 1 | 25 | X_0191 | Bacteria | Proteobacteria | Gammaproteobacteria | Pseudomonadales | Pseudomonadaceae | Unidentified | -0.273 | 0.048 | 0.063 | -0.263 |
| 1 | 26 | X_0204 | Bacteria | Bdellovibrionota | Bdellovibrionia | Bdellovibrionales | Bdellovibrionaceae | Bdellovibrio | 0.085 | 0.201 | 0.009 | -0.021 |
| 1 | 27 | X_0229 | Bacteria | Patescibacteria | Parcubacteria | Unidentified | Unidentified | Unidentified | 0.041 | 0.149 | 0.104 | -0.034 |
| 1 | 28 | X_0495 | Bacteria | Verrucomicrobiota | Chlamydiae | Chlamydiales | Chlamydiaceae | Unidentified | -0.037 | 0.172 | 0.180 | -0.114 |
| 1 | 28 | X_0230 | Bacteria | Dependentiae | Babeliae | Babeliales | Unidentified | Unidentified | -0.204 | -0.023 | 0.009 | -0.169 |
| 1 | 29 | X_0269 | Bacteria | Bacteroidota | Bacteroidia | Chitinophagales | Unidentified | Unidentified | -0.118 | 0.329 | 0.200 | -0.254 |
| 1 | 30 | X_0280 | Bacteria | Patescibacteria | Saccharimonadia | Saccharimonadales | Unidentified | Unidentified | 0.062 | -0.115 | -0.019 | 0.108 |
| 1 | 31 | X_0282 | Bacteria | Nitrospirota | Nitrospiria | Nitrospirales | Nitrospiraceae | Nitrospira | 0.001 | 0.287 | 0.070 | -0.134 |
| 1 | 32 | X_0301 | Bacteria | Unidentified | Unidentified | Unidentified | Unidentified | Unidentified | 0.092 | 0.208 | 0.193 | -0.018 |

| | | | | | | | | | | | | |
|---|---|---|---|---|---|---|---|---|---|---|---|---|
| 1 | 33 | X_0394 | Bacteria | Proteobacteria | Alphaproteobacteria | Rhizobiales | Rhizobiales Incertae Sedis | Unidentified | 0.190 | 0.381 | 0.181 | -0.025 |
| 1 | 34 | X_0429 | Bacteria | Patescibacteria | Saccharimonadia | Saccharimonadales | Unidentified | Unidentified | 0.062 | -0.098 | -0.165 | 0.100 |
| 1 | 35 | X_0433 | Bacteria | Proteobacteria | Gammaproteobacteria | Burkholderiales | Nitrosomonadaceae | Nitrosomonas | -0.089 | 0.160 | 0.024 | -0.153 |
| 1 | 36 | X_0452 | Bacteria | Unidentified | Unidentified | Unidentified | Unidentified | Unidentified | -0.111 | 0.187 | 0.076 | -0.184 |
| 2 | 1 | X_0025 | Bacteria | Actinobacteriota | Actinobacteria | Frankiales | Sporichthyaceae | hgcI clade | 0.226 | 0.422 | 0.031 | -0.028 |
| 2 | 1 | X_0019 | Bacteria | Bacteroidota | Bacteroidia | Chitinophagales | Unidentified | Unidentified | 0.224 | 0.492 | 0.090 | -0.071 |
| 2 | 1 | X_0058 | Bacteria | Patescibacteria | Parcubacteria | Candidatus Nomurabacteria | Unidentified | Unidentified | 0.217 | 0.480 | 0.213 | -0.069 |
| 2 | 1 | X_0010 | Bacteria | Actinobacteriota | Actinobacteria | Micrococcales | Microbacteriaceae | Aurantimicrobium | 0.202 | 0.558 | 0.160 | -0.127 |
| 2 | 1 | X_0113 | Bacteria | Patescibacteria | Gracilibacteria | Candidatus Peribacteria | Unidentified | Unidentified | 0.124 | 0.453 | 0.179 | -0.125 |
| 2 | 1 | X_0277 | Bacteria | Patescibacteria | Parcubacteria | Candidatus Staskawiczbacteria | Unidentified | Unidentified | 0.116 | -0.022 | -0.044 | 0.112 |
| 2 | 1 | X_0004 | Bacteria | Bacteroidota | Bacteroidia | Flavobacteriales | Flavobacteriaceae | Flavobacterium | 0.115 | 0.597 | 0.182 | -0.211 |
| 2 | 1 | X_0075 | Bacteria | Patescibacteria | ABY1 | Candidatus Magasanikbacteria | Unidentified | Unidentified | 0.114 | 0.362 | 0.102 | -0.085 |
| 2 | 1 | X_0229 | Bacteria | Patescibacteria | Parcubacteria | Unidentified | Unidentified | Unidentified | 0.106 | 0.153 | 0.196 | 0.017 |
| 2 | 1 | X_0248 | Bacteria | Proteobacteria | Gammaproteobacteria | Burkholderiales | Comamonadaceae | Unidentified | 0.099 | 0.395 | 0.067 | -0.114 |
| 2 | 1 | X_0008 | Bacteria | Proteobacteria | Alphaproteobacteria | Rhizobiales | Rhizobiales Incertae Sedis | Unidentified | 0.098 | 0.523 | 0.108 | -0.183 |
| 2 | 1 | X_0266 | Bacteria | Verrucomicrobiota | Omnitrophia | Omnitrophales | Omnitrophaceae | Candidatus Omnitrophus | 0.078 | 0.357 | 0.111 | -0.111 |
| 2 | 1 | X_0228 | Bacteria | Patescibacteria | WWE3 | Unidentified | Unidentified | Unidentified | 0.065 | 0.227 | 0.110 | -0.056 |
| 2 | 1 | X_0069 | Archaea | Nanoarchaeota | Nanoarchaeia | Woesearchaeales | GW2011_GWC1_47_15 | Unidentified | 0.059 | 0.301 | 0.113 | -0.098 |
| 2 | 1 | X_0160 | Bacteria | Patescibacteria | Gracilibacteria | Candidatus Peregrinibacteria | Unidentified | Unidentified | 0.052 | 0.178 | 0.210 | -0.042 |
| 2 | 1 | X_0123 | Bacteria | Verrucomicrobiota | Chlamydiae | Chlamydiales | cvE6 | Unidentified | 0.049 | 0.166 | 0.087 | -0.039 |
| 2 | 1 | X_0055 | Bacteria | Patescibacteria | Microgenomatia | Candidatus Woesebacteria | Unidentified | Unidentified | 0.035 | 0.272 | 0.129 | -0.104 |

| | | | | | | | | | | | | |
|---|---|---|---|---|---|---|---|---|---|---|---|---|
| 2 | 1 | X_0022 | Bacteria | Proteobacteria | Gammaproteobacteria | Burkholderiales | Comamonadaceae | Ottowia | 0.024 | 0.400 | 0.098 | -0.177 |
| 2 | 1 | X_0401 | Bacteria | Patescibacteria | Microgenomatia | Candidatus Woesebacteria | Unidentified | Unidentified | 0.024 | 0.374 | 0.150 | -0.164 |
| 2 | 1 | X_0148 | Bacteria | Patescibacteria | Saccharimonadia | Saccharimonadales | Unidentified | Unidentified | 0.000 | 0.422 | 0.218 | -0.207 |
| 2 | 1 | X_0018 | Bacteria | Proteobacteria | Gammaproteobacteria | Burkholderiales | Comamonadaceae | Giesbergeria | 0.000 | 0.659 | 0.295 | -0.323 |
| 2 | 1 | X_0042 | Bacteria | Bacteroidota | Bacteroidia | Flavobacteriales | Flavobacteriaceae | Flavobacterium | -0.008 | 0.614 | 0.364 | -0.306 |
| 2 | 1 | X_0001 | Bacteria | Bacteroidota | Bacteroidia | Flavobacteriales | Flavobacteriaceae | Flavobacterium | -0.014 | 0.704 | 0.313 | -0.353 |
| 2 | 1 | X_0095 | Bacteria | Patescibacteria | Saccharimonadia | Saccharimonadales | Unidentified | Unidentified | -0.026 | 0.553 | 0.217 | -0.289 |
| 2 | 1 | X_0013 | Bacteria | Bacteroidota | Bacteroidia | Chitinophagales | Chitinophagaceae | Sediminibacterium | -0.030 | 0.620 | 0.283 | -0.324 |
| 2 | 1 | X_0084 | Archaea | Crenarchaeota | Nitrososphaeria | Nitrosopumilales | Nitrosopumilaceae | Candidatus Nitrosotenuis | -0.034 | 0.456 | 0.208 | -0.249 |
| 2 | 1 | X_0109 | Bacteria | Proteobacteria | Gammaproteobacteria | Pseudomonadales | Moraxellaceae | Acinetobacter | -0.041 | 0.127 | 0.119 | -0.098 |
| 2 | 1 | X_0035 | Bacteria | Verrucomicrobiota | Verrucomicrobiae | Unidentified | Unidentified | Unidentified | -0.043 | 0.290 | 0.210 | -0.178 |
| 2 | 1 | X_0033 | Bacteria | Proteobacteria | Gammaproteobacteria | Xanthomonadales | Xanthomonadaceae | Arenimonas | -0.047 | 0.702 | 0.308 | -0.372 |
| 2 | 1 | X_0111 | Bacteria | Verrucomicrobiota | Verrucomicrobiae | Verrucomicrobiales | Rubritaleaceae | Luteolibacter | -0.048 | 0.747 | 0.375 | -0.393 |
| 2 | 1 | X_0003 | Bacteria | Actinobacteriota | Actinobacteria | Frankiales | Sporichthyaceae | Unidentified | -0.052 | 0.496 | 0.278 | -0.281 |
| 2 | 1 | X_0169 | Bacteria | Patescibacteria | Gracilibacteria | Candidatus Peribacteria | Unidentified | Unidentified | -0.054 | 0.573 | 0.343 | -0.319 |
| 2 | 1 | X_0088 | Bacteria | Patescibacteria | Parcubacteria | Candidatus Campbellbacteria | Unidentified | Unidentified | -0.055 | 0.547 | 0.367 | -0.307 |
| 2 | 1 | X_0031 | Bacteria | Proteobacteria | Gammaproteobacteria | Burkholderiales | Burkholderiaceae | Polynucleobacter | -0.076 | 0.614 | 0.271 | -0.353 |
| 2 | 1 | X_0017 | Bacteria | Bacteroidota | Bacteroidia | Chitinophagales | Chitinophagaceae | Edaphobaculum | -0.096 | 0.165 | 0.052 | -0.163 |
| 2 | 1 | X_0080 | Bacteria | Proteobacteria | Gammaproteobacteria | Pseudomonadales | Moraxellaceae | Acinetobacter | -0.153 | 0.261 | 0.183 | -0.256 |
| 2 | 1 | X_0299 | Bacteria | Patescibacteria | Gracilibacteria | Candidatus Peribacteria | Unidentified | Unidentified | -0.161 | 0.135 | 0.116 | -0.206 |
| 2 | 1 | X_0056 | Bacteria | Verrucomicrobiota | Verrucomicrobiae | Unidentified | Unidentified | Unidentified | -0.236 | 0.630 | 0.360 | -0.468 |

| | | | | | | | | | | | | |
|---|---|---|---|---|---|---|---|---|---|---|---|---|
| 2 | 1 | X_0089 | Bacteria | Bacteroidota | Kapabacteria | Kapabacteriales | Unidentified | Unidentified | -0.255 | 0.507 | 0.157 | -0.439 |
| 2 | 2 | X_0002 | Bacteria | Fusobacteriota | Fusobacteriia | Fusobacteriales | Fusobacteriaceae | Cetobacterium | 0.284 | -0.503 | -0.331 | 0.460 |
| 2 | 2 | X_0020 | Bacteria | Proteobacteria | Gammaproteobacteria | Enterobacterales | Enterobacteriaceae | Plesiomonas | 0.274 | -0.221 | -0.207 | 0.341 |
| 2 | 2 | X_0272 | Bacteria | Spirochaetota | Brevinematia | Brevinematales | Brevinemataceae | Brevinema | 0.209 | -0.080 | -0.143 | 0.221 |
| 2 | 2 | X_0029 | Bacteria | Firmicutes | Clostridia | Clostridiales | Clostridiaceae | Clostridium sensu stricto 1 | 0.182 | -0.135 | -0.211 | 0.223 |
| 2 | 2 | X_0082 | Bacteria | Bacteroidota | Bacteroidia | Chitinophagales | Saprospiraceae | Haliscomenobacter | 0.168 | 0.111 | -0.116 | 0.092 |
| 2 | 2 | X_0027 | Bacteria | Proteobacteria | Gammaproteobacteria | Enterobacterales | Hafniaceae | Edwardsiella | 0.152 | -0.466 | -0.361 | 0.345 |
| 2 | 2 | X_0196 | Bacteria | Proteobacteria | Gammaproteobacteria | Xanthomonadales | Xanthomonadaceae | Thermomonas | 0.144 | -0.405 | -0.254 | 0.313 |
| 2 | 2 | X_0073 | Bacteria | Proteobacteria | Gammaproteobacteria | Burkholderiales | Comamonadaceae | Curvibacter | 0.107 | -0.591 | -0.383 | 0.364 |
| 2 | 2 | X_0014 | Bacteria | Firmicutes | Clostridia | Peptostreptococcales-Tissierellales | Peptostreptococcaceae | Paraclostridium | 0.105 | -0.718 | -0.403 | 0.415 |
| 2 | 2 | X_0183 | Bacteria | Firmicutes | Clostridia | Clostridiales | Clostridiaceae | Unidentified | 0.097 | -0.254 | -0.201 | 0.206 |
| 2 | 2 | X_0344 | Bacteria | Bacteroidota | Bacteroidia | Flavobacteriales | Flavobacteriaceae | Flavobacterium | 0.097 | -0.048 | -0.379 | 0.108 |
| 2 | 2 | X_0127 | Bacteria | Firmicutes | Bacilli | Mycoplasmatales | Mycoplasmataceae | Unidentified | 0.067 | -0.468 | -0.297 | 0.281 |
| 2 | 2 | X_0121 | Bacteria | Bacteroidota | Bacteroidia | Chitinophagales | Chitinophagaceae | Parasediminibacterium | 0.062 | -0.284 | -0.099 | 0.191 |
| 2 | 2 | X_0233 | Bacteria | Proteobacteria | Alphaproteobacteria | Sphingomonadales | Sphingomonadaceae | Sphingopyxis | 0.060 | -0.177 | -0.247 | 0.138 |
| 2 | 2 | X_0066 | Bacteria | Proteobacteria | Gammaproteobacteria | Enterobacterales | Enterobacteriaceae | Citrobacter | 0.058 | -0.338 | -0.315 | 0.213 |
| 2 | 2 | X_0226 | Bacteria | Firmicutes | Clostridia | Clostridiales | Clostridiaceae | Clostridium sensu stricto 1 | 0.047 | -0.119 | -0.112 | 0.099 |
| 2 | 2 | X_0028 | Bacteria | Firmicutes | Clostridia | Peptostreptococcales-Tissierellales | Peptostreptococcaceae | Romboutsia | 0.028 | -0.644 | -0.337 | 0.334 |
| 2 | 2 | X_0346 | Bacteria | Firmicutes | Bacilli | Lactobacillales | Streptococcaceae | Lactococcus | -0.115 | -0.086 | -0.194 | -0.057 |

| | | | | | | | | | | | | |
|---|---|---|---|---|---|---|---|---|---|---|---|---|
| 2 | 2 | X_0155 | Bacteria | Proteobacteria | Alphaproteobacteria | Rhizobiales | Rhizobiaceae | Unidentified | -0.136 | -0.253 | -0.173 | 0.009 |
| 2 | 3 | X_0133 | Bacteria | Proteobacteria | Alphaproteobacteria | Rhodobacterales | Rhodobacteraceae | Tabrizicola | 0.166 | 0.243 | -0.095 | 0.022 |
| 2 | 3 | X_0292 | Bacteria | Dependentiae | Babeliae | Babeliales | UBA12409 | Unidentified | 0.156 | 0.264 | 0.066 | 0.002 |
| 2 | 3 | X_0509 | Bacteria | Patescibacteria | Saccharimonadia | Saccharimonadales | Unidentified | Unidentified | 0.140 | 0.197 | 0.050 | 0.024 |
| 2 | 3 | X_0078 | Bacteria | Proteobacteria | Gammaproteobacteria | Alteromonadales | Alteromonadaceae | Rheinheimera | 0.098 | 0.304 | 0.035 | -0.067 |
| 2 | 3 | X_0124 | Bacteria | Proteobacteria | Gammaproteobacteria | Burkholderiales | Leeiaceae | Leeia | 0.076 | -0.122 | 0.016 | 0.126 |
| 2 | 3 | X_0119 | Bacteria | Patescibacteria | Parcubacteria | Candidatus Zambryskibacteria | Unidentified | Unidentified | 0.013 | 0.164 | 0.274 | -0.069 |
| 2 | 3 | X_0162 | Bacteria | Proteobacteria | Gammaproteobacteria | Burkholderiales | Comamonadaceae | Ideonella | 0.003 | 0.264 | 0.001 | -0.126 |
| 2 | 3 | X_0052 | Bacteria | Patescibacteria | Parcubacteria | Unidentified | Unidentified | Unidentified | -0.001 | 0.367 | 0.318 | -0.180 |
| 2 | 3 | X_0140 | Bacteria | Bacteroidota | Bacteroidia | Chitinophagales | Saprospiraceae | Unidentified | -0.039 | 0.439 | 0.069 | -0.245 |
| 2 | 3 | X_0005 | Bacteria | Actinobacteriota | Actinobacteria | Micrococcales | Microbacteriaceae | Unidentified | -0.040 | -0.228 | -0.084 | 0.078 |
| 2 | 3 | X_0174 | Bacteria | Patescibacteria | Saccharimonadia | Saccharimonadales | Unidentified | Unidentified | -0.089 | 0.495 | 0.236 | -0.309 |
| 2 | 3 | X_0217 | Bacteria | Patescibacteria | Gracilibacteria | Absconditabacteriales (SR1) | Unidentified | Unidentified | -0.157 | 0.319 | 0.141 | -0.286 |
| 2 | 3 | X_0146 | Bacteria | Unidentified | Unidentified | Unidentified | Unidentified | Unidentified | -0.271 | 0.546 | 0.499 | -0.465 |
| 2 | 3 | X_0068 | Bacteria | Patescibacteria | Saccharimonadia | Saccharimonadales | Saccharimonadaceae | TM7a | -0.296 | 0.275 | 0.222 | -0.383 |
| 2 | 3 | X_0051 | Bacteria | Bacteroidota | Bacteroidia | Flavobacteriales | Flavobacteriaceae | Flavobacterium | -0.334 | 0.457 | 0.414 | -0.483 |
| 2 | 4 | X_0006 | Bacteria | Bacteroidota | Bacteroidia | Flavobacteriales | Weeksellaceae | Unidentified | 0.152 | 0.134 | -0.062 | 0.066 |
| 2 | 5 | X_0007 | Bacteria | Proteobacteria | Alphaproteobacteria | Sphingomonadales | Sphingomonadaceae | Novosphingobium | -0.222 | -0.131 | 0.100 | -0.128 |
| 2 | 6 | X_0106 | Bacteria | Bacteroidota | Bacteroidia | Flavobacteriales | Crocinitomicaceae | Fluviicola | 0.125 | -0.251 | -0.039 | 0.228 |
| 2 | 6 | X_0009 | Bacteria | Proteobacteria | Gammaproteobacteria | Burkholderiales | Comamonadaceae | Unidentified | 0.023 | -0.113 | -0.048 | 0.075 |
| 2 | 7 | X_0015 | Bacteria | Bacteroidota | Bacteroidia | Chitinophagales | Chitinophagaceae | Unidentified | 0.371 | 0.198 | -0.154 | 0.221 |

| | | | | | | | | | | | | |
|---|---|---|---|---|---|---|---|---|---|---|---|---|
| 2 | 7 | X_0050 | Bacteria | Patescibacteria | Gracilibacteria | Absconditabacteriales (SR1) | Unidentified | Unidentified | 0.206 | -0.672 | -0.454 | 0.461 |
| 2 | 7 | X_0221 | Bacteria | Patescibacteria | Gracilibacteria | Absconditabacteriales (SR1) | Unidentified | Unidentified | 0.181 | -0.088 | -0.245 | 0.200 |
| 2 | 7 | X_0367 | Bacteria | Patescibacteria | Parcubacteria | Candidatus Nomurabacteria | Unidentified | Unidentified | 0.162 | -0.303 | -0.343 | 0.283 |
| 2 | 7 | X_0366 | Bacteria | Patescibacteria | Parcubacteria | Unidentified | Unidentified | Unidentified | 0.054 | -0.306 | -0.333 | 0.194 |
| 2 | 7 | X_0011 | Bacteria | Actinobacteriota | Actinobacteria | Micrococcales | Microbacteriaceae | Unidentified | -0.044 | -0.712 | -0.374 | 0.321 |
| 2 | 8 | X_0012 | Bacteria | Proteobacteria | Gammaproteobacteria | Burkholderiales | Neisseriaceae | Unidentified | 0.120 | -0.591 | -0.448 | 0.374 |
| 2 | 9 | X_0016 | Bacteria | Bacteroidota | Bacteroidia | Flavobacteriales | Weeksellaceae | Cloacibacterium | 0.030 | -0.091 | -0.112 | 0.070 |
| 2 | 10 | X_0021 | Bacteria | Deinococcota | Deinococci | Deinococcales | Deinococcaceae | Deinobacterium | 0.021 | 0.114 | 0.290 | -0.038 |
| 2 | 11 | X_0040 | Bacteria | Patescibacteria | Parcubacteria | Candidatus Nomurabacteria | Unidentified | Unidentified | 0.087 | -0.583 | -0.219 | 0.347 |
| 2 | 11 | X_0090 | Bacteria | Cyanobacteria | Vampirivibrionia | Obscuribacterales | Obscuribacteraceae | Candidatus Obscuribacter | 0.080 | -0.395 | -0.289 | 0.257 |
| 2 | 11 | X_0239 | Bacteria | Proteobacteria | Gammaproteobacteria | Oceanospirillales | Halomonadaceae | Halomonas | 0.073 | -0.280 | -0.138 | 0.198 |
| 2 | 11 | X_0023 | Bacteria | Bacteroidota | Bacteroidia | Flavobacteriales | Flavobacteriaceae | Flavobacterium | 0.071 | -0.693 | -0.358 | 0.383 |
| 2 | 11 | X_0263 | Archaea | Nanoarchaeota | Nanoarchaeia | Woesearchaeales | SCGC AAA011-D5 | Unidentified | 0.065 | -0.376 | -0.095 | 0.236 |
| 2 | 11 | X_0158 | Bacteria | Bdellovibrionota | Oligoflexia | 0319-6G20 | Unidentified | Unidentified | 0.048 | -0.457 | -0.086 | 0.261 |
| 2 | 11 | X_0086 | Bacteria | Patescibacteria | Saccharimonadia | Saccharimonadales | Unidentified | Unidentified | 0.023 | -0.592 | -0.217 | 0.305 |
| 2 | 11 | X_0101 | Bacteria | Bacteroidota | Bacteroidia | Cytophagales | Microscillaceae | Unidentified | 0.021 | -0.493 | -0.182 | 0.257 |
| 2 | 11 | X_0065 | Bacteria | Patescibacteria | Saccharimonadia | Saccharimonadales | Unidentified | Unidentified | 0.010 | -0.653 | -0.248 | 0.326 |
| 2 | 11 | X_0193 | Bacteria | Proteobacteria | Alphaproteobacteria | Rhodobacterales | Rhodobacteraceae | Defluviimonas | 0.007 | -0.418 | -0.119 | 0.210 |
| 2 | 11 | X_0059 | Bacteria | Verrucomicrobiota | Verrucomicrobiae | Verrucomicrobiales | Rubritaleaceae | Luteolibacter | 0.001 | -0.632 | -0.284 | 0.310 |
| 2 | 11 | X_0147 | Bacteria | Verrucomicrobiota | Verrucomicrobiae | Unidentified | Unidentified | Unidentified | -0.003 | -0.421 | -0.099 | 0.204 |
| 2 | 11 | X_0039 | Bacteria | Bacteroidota | Bacteroidia | Sphingobacteriales | env.OPS 17 | Unidentified | -0.008 | -0.762 | -0.334 | 0.368 |

| | | | | | | | | | | | | |
|---|---|---|---|---|---|---|---|---|---|---|---|---|
| 2 | 11 | X_0060 | Bacteria | Verrucomicrobiota | Verrucomicrobiae | Unidentified | Unidentified | Unidentified | -0.009 | -0.633 | -0.246 | 0.304 |
| 2 | 11 | X_0237 | Bacteria | Proteobacteria | Gammaproteobacteria | Diplorickettsiales | Diplorickettsiaceae | Unidentified | -0.016 | -0.353 | -0.156 | 0.160 |
| 2 | 11 | X_0071 | Bacteria | Proteobacteria | Gammaproteobacteria | Xanthomonadales | Xanthomonadaceae | Thermomonas | -0.042 | -0.505 | -0.133 | 0.216 |
| 2 | 11 | X_0046 | Bacteria | Proteobacteria | Gammaproteobacteria | Burkholderiales | Comamonadaceae | Rhizobacter | -0.045 | -0.475 | -0.107 | 0.197 |
| 2 | 11 | X_0079 | Bacteria | Bacteroidota | Bacteroidia | Sphingobacteriales | env.OPS 17 | Unidentified | -0.050 | -0.592 | -0.214 | 0.254 |
| 2 | 11 | X_0245 | Bacteria | Proteobacteria | Gammaproteobacteria | Burkholderiales | SC-I-84 | Unidentified | -0.062 | -0.053 | 0.036 | -0.028 |
| 2 | 11 | X_0268 | Bacteria | Proteobacteria | Alphaproteobacteria | Micropepsales | Micropepsaceae | Unidentified | -0.081 | -0.465 | -0.248 | 0.165 |
| 2 | 11 | X_0047 | Bacteria | Proteobacteria | Gammaproteobacteria | Xanthomonadales | Xanthomonadaceae | Thermomonas | -0.084 | -0.690 | -0.238 | 0.284 |
| 2 | 11 | X_0091 | Bacteria | Bacteroidota | Bacteroidia | Chitinophagales | Chitinophagaceae | Sediminibacterium | -0.090 | -0.635 | -0.240 | 0.250 |
| 2 | 11 | X_0333 | Bacteria | Proteobacteria | Gammaproteobacteria | Xanthomonadales | Rhodanobacteraceae | Dokdonella | -0.094 | -0.317 | -0.093 | 0.078 |
| 2 | 11 | X_0410 | Bacteria | Patescibacteria | Gracilibacteria | JGI 0000069-P22 | Unidentified | Unidentified | -0.098 | -0.584 | -0.176 | 0.217 |
| 2 | 11 | X_0062 | Bacteria | Planctomycetota | Planctomycetes | Isosphaerales | Isosphaeraceae | Tundrisphaera | -0.098 | -0.456 | -0.086 | 0.147 |
| 2 | 11 | X_0184 | Bacteria | Proteobacteria | Gammaproteobacteria | Oceanospirillales | Unidentified | Unidentified | -0.105 | -0.284 | -0.048 | 0.051 |
| 2 | 11 | X_0112 | Bacteria | Bacteroidota | Bacteroidia | Chitinophagales | Saprospiraceae | Unidentified | -0.111 | -0.475 | -0.112 | 0.147 |
| 2 | 11 | X_0240 | Bacteria | Proteobacteria | Alphaproteobacteria | Rhizobiales | Rhizobiales Incertae Sedis | Unidentified | -0.112 | -0.578 | -0.213 | 0.203 |
| 2 | 11 | X_0275 | Bacteria | Chloroflexi | Chloroflexia | Thermomicrobiales | JG30-KF-CM45 | Unidentified | -0.117 | -0.105 | -0.112 | -0.050 |
| 2 | 11 | X_0195 | Bacteria | Bdellovibrionota | Bdellovibrionia | Bdellovibrionales | Bdellovibrionaceae | Bdellovibrio | -0.120 | -0.308 | -0.239 | 0.051 |
| 2 | 11 | X_0466 | Bacteria | Patescibacteria | Parcubacteria | Unidentified | Unidentified | Unidentified | -0.126 | -0.213 | 0.014 | -0.004 |
| 2 | 11 | X_0125 | Bacteria | Bacteroidota | Bacteroidia | Chitinophagales | Chitinophagaceae | Ferruginibacter | -0.141 | -0.611 | -0.161 | 0.202 |
| 2 | 11 | X_0209 | Bacteria | Bacteroidota | Bacteroidia | Chitinophagales | Chitinophagaceae | Parafilimonas | -0.149 | -0.384 | -0.084 | 0.068 |
| 2 | 11 | X_0108 | Bacteria | Proteobacteria | Gammaproteobacteria | Burkholderiales | Comamonadaceae | Ottowia | -0.158 | -0.631 | -0.194 | 0.201 |

| | | | | | | | | | | | | | |
|---|---|---|---|---|---|---|---|---|---|---|---|---|---|
| 2 | 11 | X_0036 | Bacteria | Bacteroidota | Bacteroidia | Cytophagales | Spirosomaceae | Arsenicibacter | | -0.174 | -0.650 | -0.261 | 0.202 |
| 2 | 11 | X_0188 | Bacteria | Patescibacteria | Unidentified | Unidentified | Unidentified | Unidentified | | -0.199 | -0.267 | -0.079 | -0.037 |
| 2 | 11 | X_0120 | Bacteria | Proteobacteria | Gammaproteobacteria | Xanthomonadales | Xanthomonadaceae | Luteimonas | | -0.227 | -0.078 | 0.018 | -0.159 |
| 2 | 12 | X_0026 | Bacteria | Bacteroidota | Bacteroidia | Flavobacteriales | Flavobacteriaceae | Flavobacterium | | 0.114 | -0.248 | 0.041 | 0.218 |
| 2 | 13 | X_0181 | Bacteria | Proteobacteria | Alphaproteobacteria | Sphingomonadales | Sphingomonadaceae | Novosphingobium | | -0.033 | -0.285 | -0.017 | 0.112 |
| 2 | 13 | X_0132 | Bacteria | Patescibacteria | Parcubacteria | Candidatus Campbellbacteria | Unidentified | Unidentified | | -0.062 | -0.273 | -0.070 | 0.081 |
| 2 | 13 | X_0030 | Bacteria | Bacteroidota | Bacteroidia | Cytophagales | Spirosomaceae | Flectobacillus | | -0.154 | -0.358 | -0.026 | 0.050 |
| 2 | 14 | X_0041 | Bacteria | Firmicutes | Bacilli | Erysipelotrichales | Erysipelotrichaceae | Turicibacter | | 0.198 | -0.286 | -0.182 | 0.305 |
| 2 | 14 | X_0166 | Bacteria | Firmicutes | Clostridia | Clostridiales | Clostridiaceae | Clostridium sensu stricto 1 | | 0.181 | -0.384 | -0.194 | 0.333 |
| 2 | 14 | X_0374 | Bacteria | Acidobacteriota | Holophagae | Holophagales | Holophagaceae | Holophaga | | 0.100 | -0.535 | -0.285 | 0.335 |
| 2 | 14 | X_0177 | Bacteria | Proteobacteria | Alphaproteobacteria | Rhodobacterales | Rhodobacteraceae | Pseudorhodobacter | | 0.095 | -0.559 | -0.269 | 0.342 |
| 2 | 14 | X_0232 | Bacteria | Firmicutes | Clostridia | Lachnospirales | Lachnospiraceae | Epulopiscium | | 0.090 | -0.423 | -0.244 | 0.278 |
| 2 | 14 | X_0220 | Bacteria | Planctomycetota | Planctomycetes | Gemmatales | Gemmataceae | Tuwongella | | 0.047 | -0.536 | -0.309 | 0.297 |
| 2 | 14 | X_0596 | Bacteria | Patescibacteria | Saccharimonadia | Saccharimonadales | Unidentified | Unidentified | | -0.006 | -0.473 | -0.290 | 0.227 |
| 2 | 14 | X_0116 | Bacteria | Bacteroidota | Bacteroidia | Chitinophagales | Chitinophagaceae | Edaphobaculum | | -0.021 | -0.634 | -0.344 | 0.295 |
| 2 | 14 | X_0354 | Bacteria | Patescibacteria | Parcubacteria | Candidatus Moranbacteria | Unidentified | Unidentified | | -0.042 | -0.309 | -0.241 | 0.116 |
| 2 | 15 | X_0043 | Bacteria | Proteobacteria | Gammaproteobacteria | Burkholderiales | Oxalobacteraceae | Undibacterium | | -0.063 | -0.669 | -0.239 | 0.286 |
| 2 | 16 | X_0048 | Bacteria | Proteobacteria | Gammaproteobacteria | Steroidobacterales | Steroidobacteraceae | Unidentified | | -0.160 | -0.040 | 0.127 | -0.120 |
| 2 | 16 | X_0083 | Bacteria | Planctomycetota | Planctomycetes | Isosphaerales | Isosphaeraceae | Unidentified | | -0.179 | -0.044 | 0.148 | -0.135 |
| 2 | 17 | X_0053 | Bacteria | Bacteroidota | Bacteroidia | Chitinophagales | Chitinophagaceae | Unidentified | | -0.090 | -0.118 | 0.064 | -0.020 |
| 2 | 18 | X_0057 | Bacteria | Proteobacteria | Alphaproteobacteria | Rhodobacterales | Rhodobacteraceae | Gemmobacter | | -0.174 | -0.271 | -0.077 | -0.013 |

| | | | | | | | | | | | | |
|---|---|---|---|---|---|---|---|---|---|---|---|---|
| 2 | 19 | X_0064 | Bacteria | Bacteroidota | Bacteroidia | Bacteroidales | Barnesiellaceae | Unidentified | 0.192 | -0.188 | -0.029 | 0.256 |
| 2 | 20 | X_0070 | Bacteria | Proteobacteria | Gammaproteobacteria | Burkholderiales | Chromobacteriaceae | Vogesella | -0.096 | -0.135 | -0.058 | -0.017 |
| 2 | 21 | X_0315 | Bacteria | Patescibacteria | Gracilibacteria | Candidatus Peribacteria | Unidentified | Unidentified | 0.095 | -0.036 | 0.127 | 0.101 |
| 2 | 21 | X_0072 | Bacteria | Verrucomicrobiota | Verrucomicrobiae | Unidentified | Unidentified | Unidentified | -0.056 | -0.006 | 0.079 | -0.046 |
| 2 | 22 | X_0077 | Bacteria | Bacteroidota | Bacteroidia | Chitinophagales | Chitinophagaceae | Taibaiella | -0.055 | -0.289 | -0.144 | 0.096 |
| 2 | 23 | X_0085 | Bacteria | Proteobacteria | Gammaproteobacteria | Pseudomonadales | Pseudomonadaceae | Pseudomonas | 0.114 | -0.009 | 0.069 | 0.104 |
| 2 | 24 | X_0093 | Bacteria | Proteobacteria | Gammaproteobacteria | Xanthomonadales | Xanthomonadaceae | Thermomonas | 0.159 | -0.178 | -0.247 | 0.224 |
| 2 | 25 | X_0096 | Bacteria | Bacteroidota | Bacteroidia | Cytophagales | Spirosomaceae | Emticicia | 0.079 | -0.289 | -0.198 | 0.207 |
| 2 | 26 | X_0159 | Bacteria | Proteobacteria | Gammaproteobacteria | Burkholderiales | Comamonadaceae | Rhizobacter | -0.060 | 0.048 | 0.103 | -0.075 |
| 2 | 26 | X_0099 | Bacteria | Planctomycetota | Planctomycetes | Pirellulales | Pirellulaceae | Unidentified | -0.082 | 0.039 | 0.280 | -0.091 |
| 2 | 27 | X_0103 | Bacteria | Bacteroidota | Bacteroidia | Flavobacteriales | Weeksellaceae | Bergeyella | 0.015 | -0.121 | -0.057 | 0.072 |
| 2 | 28 | X_0107 | Bacteria | Proteobacteria | Gammaproteobacteria | Aeromonadales | Aeromonadaceae | Aeromonas | -0.117 | 0.051 | 0.130 | -0.127 |
| 2 | 29 | X_0115 | Bacteria | Patescibacteria | Gracilibacteria | Absconditabacteriales (SR1) | Unidentified | Unidentified | 0.193 | -0.125 | -0.215 | 0.228 |
| 2 | 30 | X_0126 | Bacteria | Nitrospirota | Nitrospiria | Nitrospirales | Nitrospiraceae | Nitrospira | -0.034 | 0.025 | 0.025 | -0.042 |
| 2 | 31 | X_0129 | Bacteria | Bacteroidota | Bacteroidia | Cytophagales | Spirosomaceae | Runella | 0.074 | -0.047 | -0.069 | 0.087 |
| 2 | 32 | X_0130 | Bacteria | Patescibacteria | Gracilibacteria | Unidentified | Unidentified | Unidentified | -0.032 | -0.064 | 0.111 | 0.004 |
| 2 | 33 | X_0134 | Bacteria | Verrucomicrobiota | Chlamydiae | Chlamydiales | cvE6 | Unidentified | -0.056 | -0.043 | 0.043 | -0.028 |
| 2 | 34 | X_0135 | Bacteria | Bacteroidota | Bacteroidia | Chitinophagales | 37-13 | Unidentified | 0.116 | -0.546 | -0.243 | 0.352 |
| 2 | 35 | X_0219 | Bacteria | Bacteroidota | Bacteroidia | Flavobacteriales | Flavobacteriaceae | Flavobacterium | 0.085 | -0.139 | -0.191 | 0.142 |
| 2 | 35 | X_0136 | Bacteria | Proteobacteria | Alphaproteobacteria | Sphingomonadales | Sphingomonadaceae | Novosphingobium | 0.040 | -0.142 | -0.074 | 0.104 |
| 2 | 36 | X_0149 | Bacteria | Bacteroidota | Kapabacteria | Kapabacteriales | Unidentified | Unidentified | -0.038 | -0.037 | 0.079 | -0.015 |

| | | | | | | | | | | | | |
|---|---|---|---|---|---|---|---|---|---|---|---|---|
| 2 | 37 | X_0151 | Bacteria | Proteobacteria | Alphaproteobacteria | Rhizobiales | Rhizobiales Incertae Sedis | Phreatobacter | -0.084 | -0.048 | -0.037 | -0.050 |
| 2 | 38 | X_0152 | Bacteria | Proteobacteria | Alphaproteobacteria | Rhizobiales | Beijerinckiaceae | Methylobacterium-Methylorubrum | 0.074 | -0.037 | -0.009 | 0.083 |
| 2 | 39 | X_0165 | Bacteria | Bacteroidota | Bacteroidia | Flavobacteriales | Weeksellaceae | Chryseobacterium | -0.193 | 0.045 | 0.085 | -0.190 |
| 2 | 40 | X_0172 | Bacteria | Proteobacteria | Alphaproteobacteria | Acetobacterales | Acetobacteraceae | Rhodovastum | -0.116 | -0.156 | 0.010 | -0.024 |
| 2 | 41 | X_0173 | Bacteria | Proteobacteria | Alphaproteobacteria | Acetobacterales | Acetobacteraceae | Roseomonas | -0.012 | -0.221 | -0.038 | 0.098 |
| 2 | 42 | X_0185 | Bacteria | Bdellovibrionota | Bdellovibrionia | Bdellovibrionales | Bdellovibrionaceae | Bdellovibrio | 0.056 | 0.021 | 0.033 | 0.039 |
| 2 | 43 | X_0191 | Bacteria | Proteobacteria | Gammaproteobacteria | Pseudomonadales | Pseudomonadaceae | Unidentified | -0.035 | -0.150 | -0.058 | 0.043 |
| 2 | 44 | X_0192 | Bacteria | Bacteroidota | Bacteroidia | Cytophagales | Spirosomaceae | Unidentified | 0.020 | 0.061 | 0.043 | -0.012 |
| 2 | 45 | X_0203 | Bacteria | Firmicutes | Bacilli | Erysipelotrichales | Erysipelotrichaceae | Turicibacter | -0.149 | -0.315 | -0.005 | 0.031 |
| 2 | 46 | X_0204 | Bacteria | Bdellovibrionota | Bdellovibrionia | Bdellovibrionales | Bdellovibrionaceae | Bdellovibrio | 0.031 | -0.107 | -0.088 | 0.080 |
| 2 | 47 | X_0218 | Archaea | Nanoarchaeota | Nanoarchaeia | Woesearchaeales | Unidentified | Unidentified | -0.054 | -0.346 | -0.141 | 0.125 |
| 2 | 48 | X_0223 | Bacteria | Bacteroidota | Bacteroidia | Chitinophagales | Chitinophagaceae | Niabella | -0.091 | 0.140 | 0.233 | -0.147 |
| 2 | 49 | X_0230 | Bacteria | Dependentiae | Babeliae | Babeliales | Unidentified | Unidentified | 0.190 | -0.192 | -0.047 | 0.256 |
| 2 | 50 | X_0244 | Bacteria | Proteobacteria | Gammaproteobacteria | Gammaproteobacteria Incertae Sedis | Unknown Family | Unidentified | -0.087 | -0.190 | 0.080 | 0.019 |
| 2 | 51 | X_0262 | Bacteria | Patescibacteria | Microgenomatia | Candidatus Amesbacteria | Unidentified | Unidentified | 0.068 | -0.158 | -0.042 | 0.135 |
| 2 | 52 | X_0280 | Bacteria | Patescibacteria | Saccharimonadia | Saccharimonadales | Unidentified | Unidentified | -0.023 | -0.177 | 0.032 | 0.067 |
| 2 | 53 | X_0318 | Bacteria | Dependentiae | Babeliae | Babeliales | Vermiphilaceae | Unidentified | 0.036 | -0.031 | -0.025 | 0.047 |
| 2 | 54 | X_0326 | Bacteria | Proteobacteria | Alphaproteobacteria | Reyranellales | Reyranellaceae | Reyranella | -0.164 | 0.048 | 0.203 | -0.166 |

| | | | | | | | | | | | | |
|---|---|---|---|---|---|---|---|---|---|---|---|---|
| 2 | 55 | X_0336 | Bacteria | Firmicutes | Clostridia | Peptostreptococcales-Tissierellales | Peptostreptococcaceae | Terrisporobacter | -0.003 | -0.360 | -0.159 | 0.174 |
| 2 | 56 | X_0375 | Bacteria | Verrucomicrobiota | Chlamydiae | Chlamydiales | Simkaniaceae | Unidentified | 0.080 | -0.285 | -0.026 | 0.206 |
| 2 | 57 | X_0419 | Bacteria | Patescibacteria | ABY1 | Candidatus Magasanikbacteria | Unidentified | Unidentified | -0.060 | 0.146 | 0.177 | -0.123 |
| 2 | 58 | X_0431 | Bacteria | Patescibacteria | Gracilibacteria | Candidatus Peribacteria | Unidentified | Unidentified | 0.128 | -0.178 | -0.064 | 0.197 |
| 2 | 59 | X_0433 | Bacteria | Proteobacteria | Gammaproteobacteria | Burkholderiales | Nitrosomonadaceae | Nitrosomonas | -0.022 | -0.149 | 0.087 | 0.054 |
| 2 | 60 | X_0484 | Bacteria | Patescibacteria | Saccharimonadia | Saccharimonadales | Unidentified | Unidentified | -0.151 | 0.039 | 0.189 | -0.151 |
| 2 | 61 | X_0492 | Bacteria | Proteobacteria | Alphaproteobacteria | Rickettsiales | Candidatus Jidaibacter | Unidentified | 0.174 | -0.114 | -0.054 | 0.206 |
| 2 | 62 | X_0541 | Bacteria | Patescibacteria | Saccharimonadia | Saccharimonadales | LWQ8 | Unidentified | -0.066 | 0.067 | -0.034 | -0.090 |
| 2 | 63 | X_0819 | Bacteria | Patescibacteria | WWE3 | Unidentified | Unidentified | Unidentified | -0.145 | 0.003 | 0.057 | -0.128 |
| 3 | 1 | X_0075 | Bacteria | Patescibacteria | ABY1 | Candidatus Magasanikbacteria | Unidentified | Unidentified | 0.284 | 0.354 | 0.042 | 0.284 |
| 3 | 1 | X_0175 | Archaea | Micrarchaeota | Micrarchaeia | Micrarchaeales | Unidentified | Unidentified | 0.239 | 0.148 | 0.082 | 0.244 |
| 3 | 1 | X_0008 | Bacteria | Proteobacteria | Alphaproteobacteria | Rhizobiales | Rhizobiales Incertae Sedis | Unidentified | 0.238 | 0.097 | 0.126 | 0.241 |
| 3 | 1 | X_0025 | Bacteria | Actinobacteriota | Actinobacteria | Frankiales | Sporichthyaceae | hgcI clade | 0.204 | 0.247 | 0.210 | 0.210 |
| 3 | 1 | X_0138 | Bacteria | Actinobacteriota | Actinobacteria | Corynebacteriales | Mycobacteriaceae | Mycobacterium | 0.194 | -0.046 | 0.110 | 0.191 |
| 3 | 1 | X_0094 | Bacteria | Bacteroidota | Bacteroidia | Flavobacteriales | Weeksellaceae | Cloacibacterium | 0.173 | 0.217 | 0.062 | 0.180 |
| 3 | 1 | X_0022 | Bacteria | Proteobacteria | Gammaproteobacteria | Burkholderiales | Comamonadaceae | Ottowia | 0.107 | 0.429 | 0.186 | 0.119 |
| 3 | 1 | X_0007 | Bacteria | Proteobacteria | Alphaproteobacteria | Sphingomonadales | Sphingomonadaceae | Novosphingobium | 0.106 | -0.015 | -0.063 | 0.105 |
| 3 | 1 | X_0018 | Bacteria | Proteobacteria | Gammaproteobacteria | Burkholderiales | Comamonadaceae | Giesbergeria | 0.095 | 0.289 | 0.097 | 0.106 |
| 3 | 1 | X_0037 | Bacteria | Proteobacteria | Alphaproteobacteria | Sphingomonadales | Sphingomonadaceae | Novosphingobium | 0.063 | 0.329 | 0.188 | 0.077 |
| 3 | 1 | X_0001 | Bacteria | Bacteroidota | Bacteroidia | Flavobacteriales | Flavobacteriaceae | Flavobacterium | 0.014 | 0.508 | 0.200 | 0.038 |

| | | | | | | | | | | | | |
|---|---|---|---|---|---|---|---|---|---|---|---|---|
| 3 | 1 | X_0024 | Bacteria | Bacteroidota | Bacteroidia | Sphingobacteriales | Sphingobacteriaceae | Solitalea | 0.001 | 0.245 | 0.292 | 0.014 |
| 3 | 1 | X_0035 | Bacteria | Verrucomicrobiota | Verrucomicrobiae | Unidentified | Unidentified | Unidentified | -0.017 | 0.528 | 0.344 | 0.013 |
| 3 | 1 | X_0236 | Bacteria | Verrucomicrobiota | Verrucomicrobiae | Verrucomicrobiales | Rubritaleaceae | Luteolibacter | -0.053 | 0.435 | 0.098 | -0.025 |
| 3 | 1 | X_0042 | Bacteria | Bacteroidota | Bacteroidia | Flavobacteriales | Flavobacteriaceae | Flavobacterium | -0.180 | 0.330 | 0.426 | -0.152 |
| 3 | 2 | X_0002 | Bacteria | Fusobacteriota | Fusobacteriia | Fusobacteriales | Fusobacteriaceae | Cetobacterium | 0.259 | 0.009 | -0.250 | 0.259 |
| 3 | 2 | X_0020 | Bacteria | Proteobacteria | Gammaproteobacteria | Enterobacterales | Enterobacteriaceae | Plesiomonas | 0.182 | 0.136 | -0.180 | 0.187 |
| 3 | 2 | X_0029 | Bacteria | Firmicutes | Clostridia | Clostridiales | Clostridiaceae | Clostridium sensu stricto 1 | 0.180 | 0.231 | -0.235 | 0.187 |
| 3 | 2 | X_0014 | Bacteria | Firmicutes | Clostridia | Peptostreptococcales-Tissierellales | Peptostreptococcaceae | Paraclostridium | 0.134 | -0.102 | -0.251 | 0.128 |
| 3 | 2 | X_0027 | Bacteria | Proteobacteria | Gammaproteobacteria | Enterobacterales | Hafniaceae | Edwardsiella | 0.107 | 0.175 | -0.189 | 0.115 |
| 3 | 2 | X_0064 | Bacteria | Bacteroidota | Bacteroidia | Bacteroidales | Barnesiellaceae | Unidentified | 0.094 | 0.259 | -0.030 | 0.104 |
| 3 | 2 | X_0412 | Bacteria | Firmicutes | Clostridia | Lachnospirales | Lachnospiraceae | Epulopiscium | 0.077 | 0.168 | 0.184 | 0.084 |
| 3 | 2 | X_0151 | Bacteria | Proteobacteria | Alphaproteobacteria | Rhizobiales | Rhizobiales Incertae Sedis | Phreatobacter | 0.062 | -0.009 | -0.068 | 0.061 |
| 3 | 2 | X_0041 | Bacteria | Firmicutes | Bacilli | Erysipelotrichales | Erysipelotrichaceae | Turicibacter | 0.046 | -0.070 | -0.025 | 0.042 |
| 3 | 2 | X_0375 | Bacteria | Verrucomicrobiota | Chlamydiae | Chlamydiales | Simkaniaceae | Unidentified | 0.033 | 0.086 | -0.050 | 0.037 |
| 3 | 2 | X_0166 | Bacteria | Firmicutes | Clostridia | Clostridiales | Clostridiaceae | Clostridium sensu stricto 1 | 0.014 | 0.110 | 0.040 | 0.020 |
| 3 | 2 | X_0066 | Bacteria | Proteobacteria | Gammaproteobacteria | Enterobacterales | Enterobacteriaceae | Citrobacter | -0.015 | -0.134 | -0.190 | -0.022 |
| 3 | 2 | X_0028 | Bacteria | Firmicutes | Clostridia | Peptostreptococcales-Tissierellales | Peptostreptococcaceae | Romboutsia | -0.029 | -0.153 | -0.003 | -0.037 |
| 3 | 2 | X_0109 | Bacteria | Proteobacteria | Gammaproteobacteria | Pseudomonadales | Moraxellaceae | Acinetobacter | -0.091 | 0.176 | 0.004 | -0.080 |
| 3 | 2 | X_0183 | Bacteria | Firmicutes | Clostridia | Clostridiales | Clostridiaceae | Unidentified | -0.092 | -0.016 | 0.028 | -0.093 |

| | | | | | | | | | | | | |
|---|---|---|---|---|---|---|---|---|---|---|---|---|
| 3 | 2 | X_0012 | Bacteria | Proteobacteria | Gammaproteobacteria | Burkholderiales | Neisseriaceae | Unidentified | -0.113 | -0.300 | -0.257 | -0.123 |
| 3 | 2 | X_0155 | Bacteria | Proteobacteria | Alphaproteobacteria | Rhizobiales | Rhizobiaceae | Unidentified | -0.139 | -0.186 | -0.166 | -0.146 |
| 3 | 2 | X_0057 | Bacteria | Proteobacteria | Alphaproteobacteria | Rhodobacterales | Rhodobacteraceae | Gemmobacter | -0.156 | -0.179 | -0.023 | -0.163 |
| 3 | 2 | X_0080 | Bacteria | Proteobacteria | Gammaproteobacteria | Pseudomonadales | Moraxellaceae | Acinetobacter | -0.162 | 0.058 | 0.124 | -0.158 |
| 3 | 3 | X_0003 | Bacteria | Actinobacteriota | Actinobacteria | Frankiales | Sporichthyaceae | Unidentified | 0.241 | 0.223 | -0.072 | 0.247 |
| 3 | 3 | X_0118 | Bacteria | Proteobacteria | Gammaproteobacteria | Alteromonadales | Alteromonadaceae | Rheinheimera | 0.134 | 0.240 | -0.066 | 0.142 |
| 3 | 3 | X_0010 | Bacteria | Actinobacteriota | Actinobacteria | Micrococcales | Microbacteriaceae | Aurantimicrobium | 0.086 | 0.373 | -0.056 | 0.099 |
| 3 | 3 | X_0077 | Bacteria | Bacteroidota | Bacteroidia | Chitinophagales | Chitinophagaceae | Taibaiella | 0.069 | 0.111 | -0.036 | 0.074 |
| 3 | 3 | X_0096 | Bacteria | Bacteroidota | Bacteroidia | Cytophagales | Spirosomaceae | Emticicia | 0.056 | 0.362 | -0.131 | 0.071 |
| 3 | 3 | X_0019 | Bacteria | Bacteroidota | Bacteroidia | Chitinophagales | Unidentified | Unidentified | 0.015 | 0.336 | -0.077 | 0.032 |
| 3 | 3 | X_0004 | Bacteria | Bacteroidota | Bacteroidia | Flavobacteriales | Flavobacteriaceae | Flavobacterium | 0.007 | 0.550 | 0.082 | 0.035 |
| 3 | 3 | X_0006 | Bacteria | Bacteroidota | Bacteroidia | Flavobacteriales | Weeksellaceae | Unidentified | 0.003 | 0.297 | -0.014 | 0.019 |
| 3 | 3 | X_0085 | Bacteria | Proteobacteria | Gammaproteobacteria | Pseudomonadales | Pseudomonadaceae | Pseudomonas | 0.002 | 0.023 | 0.050 | 0.003 |
| 3 | 3 | X_0283 | Bacteria | Verrucomicrobiota | Chlamydiae | Chlamydiales | Simkaniaceae | Unidentified | -0.006 | -0.082 | -0.003 | -0.010 |
| 3 | 3 | X_0009 | Bacteria | Proteobacteria | Gammaproteobacteria | Burkholderiales | Comamonadaceae | Unidentified | -0.023 | -0.192 | -0.261 | -0.032 |
| 3 | 3 | X_0026 | Bacteria | Bacteroidota | Bacteroidia | Flavobacteriales | Flavobacteriaceae | Flavobacterium | -0.071 | -0.086 | -0.368 | -0.076 |
| 3 | 3 | X_0005 | Bacteria | Actinobacteriota | Actinobacteria | Micrococcales | Microbacteriaceae | Unidentified | -0.094 | 0.074 | -0.191 | -0.089 |
| 3 | 3 | X_0013 | Bacteria | Bacteroidota | Bacteroidia | Chitinophagales | Chitinophagaceae | Sediminibacterium | -0.096 | 0.149 | 0.124 | -0.087 |
| 3 | 3 | X_0243 | Bacteria | Verrucomicrobiota | Chlamydiae | Chlamydiales | Unidentified | Unidentified | -0.125 | 0.152 | 0.001 | -0.116 |
| 3 | 3 | X_0184 | Bacteria | Proteobacteria | Gammaproteobacteria | Oceanospirillales | Unidentified | Unidentified | -0.157 | -0.117 | 0.271 | -0.162 |
| 3 | 4 | X_0032 | Bacteria | Bacteroidota | Bacteroidia | Sphingobacteriales | env.OPS 17 | Unidentified | 0.165 | -0.310 | -0.234 | 0.141 |

| | | | | | | | | | | | | |
|---|---|---|---|---|---|---|---|---|---|---|---|---|
| 3 | 4 | X_0135 | Bacteria | Bacteroidota | Bacteroidia | Chitinophagales | 37-13 | Unidentified | 0.031 | -0.134 | -0.126 | 0.024 |
| 3 | 4 | X_0202 | Bacteria | Verrucomicrobiota | Verrucomicrobiae | Pedosphaerales | Pedosphaeraceae | Unidentified | 0.031 | -0.227 | 0.020 | 0.018 |
| 3 | 4 | X_0163 | Bacteria | Actinobacteriota | Actinobacteria | Corynebacteriales | Mycobacteriaceae | Mycobacterium | 0.029 | -0.293 | 0.026 | 0.012 |
| 3 | 4 | X_0036 | Bacteria | Bacteroidota | Bacteroidia | Cytophagales | Spirosomaceae | Arsenicibacter | 0.019 | -0.224 | -0.127 | 0.007 |
| 3 | 4 | X_0173 | Bacteria | Proteobacteria | Alphaproteobacteria | Acetobacterales | Acetobacteraceae | Roseomonas | 0.012 | -0.146 | 0.026 | 0.004 |
| 3 | 4 | X_0158 | Bacteria | Bdellovibrionota | Oligoflexia | 0319-6G20 | Unidentified | Unidentified | 0.007 | -0.292 | -0.145 | -0.009 |
| 3 | 4 | X_0011 | Bacteria | Actinobacteriota | Actinobacteria | Micrococcales | Microbacteriaceae | Unidentified | 0.007 | -0.442 | -0.251 | -0.017 |
| 3 | 4 | X_0121 | Bacteria | Bacteroidota | Bacteroidia | Chitinophagales | Chitinophagaceae | Parasediminibacterium | -0.005 | 0.067 | 0.042 | -0.002 |
| 3 | 4 | X_0428 | Bacteria | Planctomycetota | Planctomycetes | Isosphaerales | Isosphaeraceae | Unidentified | -0.011 | -0.240 | -0.071 | -0.023 |
| 3 | 4 | X_0115 | Bacteria | Patescibacteria | Gracilibacteria | Absconditabacteriales (SR1) | Unidentified | Unidentified | -0.016 | -0.076 | -0.129 | -0.020 |
| 3 | 4 | X_0114 | Bacteria | Verrucomicrobiota | Verrucomicrobiae | Unidentified | Unidentified | Unidentified | -0.025 | -0.371 | -0.064 | -0.042 |
| 3 | 4 | X_0062 | Bacteria | Planctomycetota | Planctomycetes | Isosphaerales | Isosphaeraceae | Tundrisphaera | -0.025 | -0.145 | -0.101 | -0.032 |
| 3 | 4 | X_0326 | Bacteria | Proteobacteria | Alphaproteobacteria | Reyranellales | Reyranellaceae | Reyranella | -0.033 | -0.091 | 0.099 | -0.037 |
| 3 | 4 | X_0131 | Bacteria | Proteobacteria | Gammaproteobacteria | Xanthomonadales | Xanthomonadaceae | Thermomonas | -0.049 | 0.133 | -0.083 | -0.041 |
| 3 | 4 | X_0405 | Bacteria | Planctomycetota | Planctomycetes | Isosphaerales | Isosphaeraceae | Candidatus Nostocoida | -0.053 | 0.037 | 0.061 | -0.051 |
| 3 | 4 | X_0046 | Bacteria | Proteobacteria | Gammaproteobacteria | Burkholderiales | Comamonadaceae | Rhizobacter | -0.057 | -0.182 | 0.160 | -0.066 |
| 3 | 4 | X_0071 | Bacteria | Proteobacteria | Gammaproteobacteria | Xanthomonadales | Xanthomonadaceae | Thermomonas | -0.062 | -0.295 | -0.020 | -0.074 |
| 3 | 4 | X_0268 | Bacteria | Proteobacteria | Alphaproteobacteria | Micropepsales | Micropepsaceae | Unidentified | -0.064 | -0.309 | 0.004 | -0.077 |
| 3 | 4 | X_0204 | Bacteria | Bdellovibrionota | Bdellovibrionia | Bdellovibrionales | Bdellovibrionaceae | Bdellovibrio | -0.068 | -0.058 | -0.086 | -0.071 |
| 3 | 4 | X_0048 | Bacteria | Proteobacteria | Gammaproteobacteria | Steroidobacterales | Steroidobacteraceae | Unidentified | -0.072 | 0.118 | 0.182 | -0.065 |
| 3 | 4 | X_0116 | Bacteria | Bacteroidota | Bacteroidia | Chitinophagales | Chitinophagaceae | Edaphobaculum | -0.111 | 0.024 | -0.026 | -0.110 |

| | | | | | | | | | | | | |
|---|---|---|---|---|---|---|---|---|---|---|---|---|
| 3 | 4 | X_0023 | Bacteria | Bacteroidota | Bacteroidia | Flavobacteriales | Flavobacteriaceae | Flavobacterium | -0.119 | -0.262 | -0.199 | -0.128 |
| 3 | 4 | X_0195 | Bacteria | Bdellovibrionota | Bdellovibrionia | Bdellovibrionales | Bdellovibrionaceae | Bdellovibrio | -0.178 | 0.150 | -0.004 | -0.168 |
| 3 | 5 | X_0017 | Bacteria | Bacteroidota | Bacteroidia | Chitinophagales | Chitinophagaceae | Edaphobaculum | 0.314 | 0.251 | 0.154 | 0.316 |
| 3 | 5 | X_0306 | Bacteria | Bacteroidota | Bacteroidia | Flavobacteriales | Crocinitomicaceae | Fluviicola | 0.242 | -0.037 | 0.032 | 0.239 |
| 3 | 5 | X_0206 | Bacteria | Patescibacteria | Unidentified | Unidentified | Unidentified | Unidentified | 0.209 | 0.113 | -0.026 | 0.213 |
| 3 | 5 | X_0221 | Bacteria | Patescibacteria | Gracilibacteria | Absconditabacteriales (SR1) | Unidentified | Unidentified | 0.207 | 0.300 | 0.045 | 0.213 |
| 3 | 5 | X_0167 | Bacteria | Bacteroidota | Bacteroidia | Sphingobacteriales | env.OPS 17 | Unidentified | 0.146 | -0.062 | 0.175 | 0.142 |
| 3 | 5 | X_0546 | Bacteria | Patescibacteria | Parcubacteria | Unidentified | Unidentified | Unidentified | 0.144 | -0.087 | 0.030 | 0.138 |
| 3 | 5 | X_0264 | Archaea | Aenigmarchaeota | Aenigmarchaeia | Aenigmarchaeales | Unidentified | Unidentified | 0.122 | -0.069 | 0.105 | 0.118 |
| 3 | 5 | X_0149 | Bacteria | Bacteroidota | Kapabacteria | Kapabacteriales | Unidentified | Unidentified | 0.121 | 0.207 | -0.010 | 0.129 |
| 3 | 5 | X_0128 | Bacteria | Verrucomicrobiota | Verrucomicrobiae | Chthoniobacterales | Terrimicrobiaceae | FukuN18 freshwater group | 0.113 | -0.151 | 0.078 | 0.104 |
| 3 | 5 | X_0261 | Archaea | Micrarchaeota | Micrarchaeia | Micrarchaeales | Unidentified | Unidentified | 0.112 | 0.063 | 0.029 | 0.115 |
| 3 | 5 | X_0315 | Bacteria | Patescibacteria | Gracilibacteria | Candidatus Peribacteria | Unidentified | Unidentified | 0.111 | 0.026 | 0.103 | 0.112 |
| 3 | 5 | X_0431 | Bacteria | Patescibacteria | Gracilibacteria | Candidatus Peribacteria | Unidentified | Unidentified | 0.082 | -0.429 | -0.038 | 0.051 |
| 3 | 5 | X_0318 | Bacteria | Dependentiae | Babeliae | Babeliales | Vermiphilaceae | Unidentified | 0.065 | -0.206 | -0.001 | 0.053 |
| 3 | 5 | X_0368 | Bacteria | Unidentified | Unidentified | Unidentified | Unidentified | Unidentified | 0.064 | -0.164 | 0.153 | 0.055 |
| 3 | 5 | X_0015 | Bacteria | Bacteroidota | Bacteroidia | Chitinophagales | Chitinophagaceae | Unidentified | 0.043 | 0.090 | 0.236 | 0.047 |
| 3 | 5 | X_0160 | Bacteria | Patescibacteria | Gracilibacteria | Candidatus Peregrinibacteria | Unidentified | Unidentified | 0.018 | 0.028 | 0.101 | 0.019 |
| 3 | 5 | X_0269 | Bacteria | Bacteroidota | Bacteroidia | Chitinophagales | Unidentified | Unidentified | -0.005 | 0.118 | 0.105 | 0.001 |
| 3 | 5 | X_0277 | Bacteria | Patescibacteria | Parcubacteria | Candidatus Staskawiczbacteria | Unidentified | Unidentified | -0.013 | -0.318 | 0.083 | -0.029 |
| 3 | 5 | X_0316 | Bacteria | Patescibacteria | Saccharimonadia | Saccharimonadales | Saccharimonadaceae | TM7a | -0.014 | -0.326 | 0.121 | -0.031 |

| | | | | | | | | | | | | |
|---|---|---|---|---|---|---|---|---|---|---|---|---|
| 3 | 5 | X_0084 | Archaea | Crenarchaeota | Nitrososphaeria | Nitrosopumilales | Nitrosopumilaceae | Candidatus Nitrosotenuis | -0.034 | 0.200 | 0.027 | -0.023 |
| 3 | 5 | X_0069 | Archaea | Nanoarchaeota | Nanoarchaeia | Woesearchaeales | GW2011_GWC1_47_15 | Unidentified | -0.046 | 0.161 | 0.179 | -0.037 |
| 3 | 5 | X_0509 | Bacteria | Patescibacteria | Saccharimonadia | Saccharimonadales | Unidentified | Unidentified | -0.058 | -0.082 | 0.069 | -0.062 |
| 3 | 5 | X_0597 | Archaea | Nanoarchaeota | Nanoarchaeia | Woesearchaeales | GW2011_GWC1_47_15 | Unidentified | -0.083 | -0.084 | 0.208 | -0.087 |
| 3 | 5 | X_0134 | Bacteria | Verrucomicrobiota | Chlamydiae | Chlamydiales | cvE6 | Unidentified | -0.085 | -0.070 | 0.054 | -0.088 |
| 3 | 5 | X_0299 | Bacteria | Patescibacteria | Gracilibacteria | Candidatus Peribacteria | Unidentified | Unidentified | -0.095 | -0.058 | -0.023 | -0.097 |
| 3 | 5 | X_0055 | Bacteria | Patescibacteria | Microgenomatia | Candidatus Woesebacteria | Unidentified | Unidentified | -0.140 | 0.134 | 0.122 | -0.131 |
| 3 | 6 | X_0016 | Bacteria | Bacteroidota | Bacteroidia | Flavobacteriales | Weeksellaceae | Cloacibacterium | 0.136 | 0.183 | -0.178 | 0.143 |
| 3 | 7 | X_0592 | Bacteria | Bacteroidota | Bacteroidia | Chitinophagales | Saprospiraceae | Unidentified | 0.210 | 0.203 | -0.053 | 0.216 |
| 3 | 7 | X_0065 | Bacteria | Patescibacteria | Saccharimonadia | Saccharimonadales | Unidentified | Unidentified | 0.203 | 0.409 | -0.006 | 0.207 |
| 3 | 7 | X_0095 | Bacteria | Patescibacteria | Saccharimonadia | Saccharimonadales | Unidentified | Unidentified | 0.154 | 0.335 | 0.034 | 0.162 |
| 3 | 7 | X_0231 | Bacteria | Bacteroidota | Bacteroidia | Cytophagales | Spirosomaceae | Runella | 0.133 | 0.455 | -0.084 | 0.142 |
| 3 | 7 | X_0097 | Bacteria | Actinobacteriota | Actinobacteria | Corynebacteriales | Mycobacteriaceae | Mycobacterium | 0.130 | 0.383 | 0.146 | 0.140 |
| 3 | 7 | X_0176 | Bacteria | Verrucomicrobiota | Verrucomicrobiae | Pedosphaerales | Pedosphaeraceae | SH3-11 | 0.116 | 0.473 | -0.008 | 0.127 |
| 3 | 7 | X_0089 | Bacteria | Bacteroidota | Kapabacteria | Kapabacteriales | Unidentified | Unidentified | 0.111 | 0.546 | 0.087 | 0.122 |
| 3 | 7 | X_0068 | Bacteria | Patescibacteria | Saccharimonadia | Saccharimonadales | Saccharimonadaceae | TM7a | 0.101 | 0.434 | -0.137 | 0.114 |
| 3 | 7 | X_0072 | Bacteria | Verrucomicrobiota | Verrucomicrobiae | Unidentified | Unidentified | Unidentified | 0.074 | 0.460 | 0.029 | 0.090 |
| 3 | 7 | X_0391 | Bacteria | Bacteroidota | Bacteroidia | Chitinophagales | Chitinophagaceae | Unidentified | 0.072 | 0.365 | 0.076 | 0.086 |
| 3 | 7 | X_0021 | Bacteria | Deinococcota | Deinococci | Deinococcales | Deinococcaceae | Deinobacterium | 0.046 | 0.313 | 0.116 | 0.060 |
| 3 | 7 | X_0129 | Bacteria | Bacteroidota | Bacteroidia | Cytophagales | Spirosomaceae | Runella | 0.035 | 0.364 | -0.007 | 0.051 |
| 3 | 7 | X_0307 | Bacteria | Bacteroidota | Bacteroidia | Chitinophagales | Chitinophagaceae | Rurimicrobium | 0.027 | 0.213 | -0.026 | 0.038 |

| | | | | | | | | | | | | |
|---|---|---|---|---|---|---|---|---|---|---|---|---|
| 3 | 7 | X_0144 | Bacteria | Bacteroidota | Bacteroidia | Unidentified | Unidentified | Unidentified | 0.012 | 0.386 | -0.059 | 0.031 |
| 3 | 7 | X_0884 | Bacteria | Actinobacteriota | Thermoleophilia | Solirubrobacterales | 67-14 | Unidentified | 0.010 | 0.323 | 0.037 | 0.026 |
| 3 | 7 | X_0126 | Bacteria | Nitrospirota | Nitrospiria | Nitrospirales | Nitrospiraceae | Nitrospira | 0.005 | 0.419 | 0.131 | 0.027 |
| 3 | 7 | X_0034 | Archaea | Crenarchaeota | Nitrososphaeria | Nitrososphaerales | Nitrososphaeraceae | Candidatus Nitrocosmicus | 0.000 | 0.486 | 0.158 | 0.025 |
| 3 | 7 | X_0192 | Bacteria | Bacteroidota | Bacteroidia | Cytophagales | Spirosomaceae | Unidentified | -0.005 | 0.504 | 0.111 | 0.022 |
| 3 | 7 | X_0031 | Bacteria | Proteobacteria | Gammaproteobacteria | Burkholderiales | Burkholderiaceae | Polynucleobacter | -0.007 | 0.511 | 0.114 | 0.021 |
| 3 | 7 | X_0466 | Bacteria | Patescibacteria | Parcubacteria | Unidentified | Unidentified | Unidentified | -0.007 | 0.270 | 0.163 | 0.008 |
| 3 | 7 | X_0172 | Bacteria | Proteobacteria | Alphaproteobacteria | Acetobacterales | Acetobacteraceae | Rhodovastum | -0.021 | 0.398 | 0.096 | 0.002 |
| 3 | 7 | X_0363 | Bacteria | Proteobacteria | Gammaproteobacteria | Xanthomonadales | Rhodanobacteraceae | Dokdonella | -0.030 | 0.320 | 0.139 | -0.011 |
| 3 | 7 | X_0083 | Bacteria | Planctomycetota | Planctomycetes | Isosphaerales | Isosphaeraceae | Unidentified | -0.047 | 0.483 | 0.115 | -0.016 |
| 3 | 7 | X_0452 | Bacteria | Unidentified | Unidentified | Unidentified | Unidentified | Unidentified | -0.061 | 0.351 | 0.003 | -0.039 |
| 3 | 7 | X_0293 | Bacteria | Proteobacteria | Gammaproteobacteria | Diplorickettsiales | Diplorickettsiaceae | Aquicella | -0.072 | 0.323 | 0.095 | -0.051 |
| 3 | 7 | X_0190 | Bacteria | Verrucomicrobiota | Verrucomicrobiae | Verrucomicrobiales | Rubritaleaceae | Luteolibacter | -0.079 | 0.558 | 0.065 | -0.036 |
| 3 | 7 | X_0033 | Bacteria | Proteobacteria | Gammaproteobacteria | Xanthomonadales | Xanthomonadaceae | Arenimonas | -0.090 | 0.598 | 0.067 | -0.040 |
| 3 | 7 | X_0323 | Archaea | Nanoarchaeota | Nanoarchaeia | Woesearchaeales | Unidentified | Unidentified | -0.101 | 0.282 | 0.010 | -0.082 |
| 3 | 7 | X_0087 | Bacteria | Bacteroidota | Bacteroidia | Chitinophagales | Chitinophagaceae | Unidentified | -0.123 | 0.637 | 0.164 | -0.061 |
| 3 | 7 | X_0049 | Bacteria | Bacteroidota | Bacteroidia | Flavobacteriales | Crocinitomicaceae | Fluviicola | -0.171 | 0.582 | 0.069 | -0.108 |
| 3 | 7 | X_0132 | Bacteria | Patescibacteria | Parcubacteria | Candidatus Campbellbacteria | Unidentified | Unidentified | -0.214 | 0.316 | 0.214 | -0.187 |
| 3 | 8 | X_0521 | Bacteria | Proteobacteria | Gammaproteobacteria | Legionellales | Legionellaceae | Unidentified | 0.054 | -0.237 | 0.116 | 0.040 |
| 3 | 8 | X_0218 | Archaea | Nanoarchaeota | Nanoarchaeia | Woesearchaeales | Unidentified | Unidentified | 0.043 | -0.386 | -0.006 | 0.020 |
| 3 | 8 | X_0449 | Bacteria | Bdellovibrionota | Oligoflexia | 0319-6G20 | Unidentified | Unidentified | 0.033 | -0.195 | -0.011 | 0.022 |

| | | | | | | | | | | | | | |
|---|---|---|---|---|---|---|---|---|---|---|---|---|---|
| 3 | 8 | X_0043 | Bacteria | Proteobacteria | Gammaproteobacteria | Burkholderiales | Oxalobacteraceae | Undibacterium | | 0.000 | -0.075 | 0.005 | -0.004 |
| 3 | 8 | X_0053 | Bacteria | Bacteroidota | Bacteroidia | Chitinophagales | Chitinophagaceae | Unidentified | | -0.001 | -0.358 | -0.055 | -0.020 |
| 3 | 8 | X_0354 | Bacteria | Patescibacteria | Parcubacteria | Candidatus Moranbacteria | Unidentified | Unidentified | | -0.015 | -0.293 | 0.033 | -0.029 |
| 3 | 8 | X_0103 | Bacteria | Bacteroidota | Bacteroidia | Flavobacteriales | Weeksellaceae | Bergeyella | | -0.016 | -0.118 | -0.090 | -0.022 |
| 3 | 8 | X_0086 | Bacteria | Patescibacteria | Saccharimonadia | Saccharimonadales | Unidentified | Unidentified | | -0.025 | -0.403 | -0.040 | -0.044 |
| 3 | 8 | X_0282 | Bacteria | Nitrospirota | Nitrospiria | Nitrospirales | Nitrospiraceae | Nitrospira | | -0.039 | -0.046 | 0.029 | -0.042 |
| 3 | 8 | X_0123 | Bacteria | Verrucomicrobiota | Chlamydiae | Chlamydiales | cvE6 | Unidentified | | -0.044 | -0.267 | 0.125 | -0.057 |
| 3 | 8 | X_0101 | Bacteria | Bacteroidota | Bacteroidia | Cytophagales | Microscillaceae | Unidentified | | -0.045 | -0.264 | -0.060 | -0.057 |
| 3 | 8 | X_0586 | Bacteria | Proteobacteria | Gammaproteobacteria | Diplorickettsiales | Diplorickettsiaceae | Aquicella | | -0.048 | -0.131 | 0.090 | -0.054 |
| 3 | 8 | X_0263 | Archaea | Nanoarchaeota | Nanoarchaeia | Woesearchaeales | SCGC AAA011-D5 | Unidentified | | -0.049 | -0.175 | 0.120 | -0.057 |
| 3 | 8 | X_0063 | Bacteria | Proteobacteria | Alphaproteobacteria | Rhizobiales | Devosiaceae | Devosia | | -0.049 | -0.635 | 0.025 | -0.071 |
| 3 | 8 | X_0182 | Bacteria | Bacteroidota | Bacteroidia | Chitinophagales | Chitinophagaceae | Unidentified | | -0.053 | 0.291 | 0.086 | -0.036 |
| 3 | 8 | X_0187 | Bacteria | Bacteroidota | Bacteroidia | Sphingobacteriales | AKYH767 | Unidentified | | -0.059 | -0.376 | 0.154 | -0.074 |
| 3 | 8 | X_0059 | Bacteria | Verrucomicrobiota | Verrucomicrobiae | Verrucomicrobiales | Rubritaleaceae | Luteolibacter | | -0.061 | -0.220 | 0.180 | -0.071 |
| 3 | 8 | X_0091 | Bacteria | Bacteroidota | Bacteroidia | Chitinophagales | Chitinophagaceae | Sediminibacterium | | -0.063 | -0.461 | 0.056 | -0.080 |
| 3 | 8 | X_0641 | Bacteria | Unidentified | Unidentified | Unidentified | Unidentified | Unidentified | | -0.067 | -0.248 | 0.087 | -0.078 |
| 3 | 8 | X_0387 | Bacteria | Proteobacteria | Gammaproteobacteria | Xanthomonadales | Rhodanobacteraceae | Rudaea | | -0.068 | 0.054 | -0.153 | -0.065 |
| 3 | 8 | X_0108 | Bacteria | Proteobacteria | Gammaproteobacteria | Burkholderiales | Comamonadaceae | Ottowia | | -0.075 | -0.089 | 0.137 | -0.080 |
| 3 | 8 | X_0787 | Bacteria | Patescibacteria | Unidentified | Unidentified | Unidentified | Unidentified | | -0.085 | -0.270 | 0.170 | -0.096 |
| 3 | 8 | X_0047 | Bacteria | Proteobacteria | Gammaproteobacteria | Xanthomonadales | Xanthomonadaceae | Thermomonas | | -0.085 | -0.544 | -0.027 | -0.100 |
| 3 | 8 | X_0215 | Bacteria | Patescibacteria | Saccharimonadia | Saccharimonadales | Unidentified | Unidentified | | -0.088 | -0.438 | 0.025 | -0.102 |

| | | | | | | | | | | | | |
|---|---|---|---|---|---|---|---|---|---|---|---|---|
| 3 | 8 | X_0120 | Bacteria | Proteobacteria | Gammaproteobacteria | Xanthomonadales | Xanthomonadaceae | Luteimonas | -0.093 | 0.107 | 0.162 | -0.087 |
| 3 | 8 | X_0050 | Bacteria | Patescibacteria | Gracilibacteria | Absconditabacteriales (SR1) | Unidentified | Unidentified | -0.095 | -0.328 | -0.080 | -0.107 |
| 3 | 8 | X_0040 | Bacteria | Patescibacteria | Parcubacteria | Candidatus Nomurabacteria | Unidentified | Unidentified | -0.099 | -0.499 | 0.044 | -0.111 |
| 3 | 8 | X_0030 | Bacteria | Bacteroidota | Bacteroidia | Cytophagales | Spirosomaceae | Flectobacillus | -0.111 | -0.265 | -0.200 | -0.121 |
| 3 | 8 | X_0098 | Bacteria | Proteobacteria | Alphaproteobacteria | Rhizobiales | Beijerinckiaceae | alphaI cluster | -0.114 | -0.573 | -0.111 | -0.123 |
| 3 | 8 | X_0360 | Bacteria | Patescibacteria | Gracilibacteria | Candidatus Peregrinibacteria | Unidentified | Unidentified | -0.127 | -0.288 | 0.038 | -0.137 |
| 3 | 8 | X_0107 | Bacteria | Proteobacteria | Gammaproteobacteria | Aeromonadales | Aeromonadaceae | Aeromonas | -0.154 | -0.127 | 0.185 | -0.160 |
| 3 | 8 | X_0284 | Bacteria | Proteobacteria | Alphaproteobacteria | Rhodospirillales | Unidentified | Unidentified | -0.155 | -0.558 | -0.032 | -0.157 |
| 3 | 8 | X_0541 | Bacteria | Patescibacteria | Saccharimonadia | Saccharimonadales | LWQ8 | Unidentified | -0.164 | -0.205 | 0.085 | -0.171 |
| 3 | 8 | X_0235 | Bacteria | Planctomycetota | Planctomycetes | Pirellulales | Pirellulaceae | Unidentified | -0.167 | -0.478 | -0.025 | -0.171 |
| 3 | 8 | X_0254 | Archaea | Nanoarchaeota | Nanoarchaeia | Woesearchaeales | GW2011_GWC1_47_15 | Unidentified | -0.167 | -0.471 | -0.038 | -0.172 |
| 3 | 8 | X_0189 | Bacteria | Planctomycetota | Planctomycetes | Pirellulales | Pirellulaceae | Pirellula | -0.169 | -0.271 | 0.000 | -0.177 |
| 3 | 8 | X_0328 | Bacteria | Patescibacteria | ABY1 | Candidatus Magasanikbacteria | Unidentified | Unidentified | -0.169 | -0.547 | -0.044 | -0.170 |
| 3 | 8 | X_0044 | Bacteria | Unidentified | Unidentified | Unidentified | Unidentified | Unidentified | -0.174 | -0.181 | 0.166 | -0.180 |
| 3 | 8 | X_0112 | Bacteria | Bacteroidota | Bacteroidia | Chitinophagales | Saprospiraceae | Unidentified | -0.177 | -0.454 | 0.106 | -0.182 |
| 3 | 8 | X_0060 | Bacteria | Verrucomicrobiota | Verrucomicrobiae | Unidentified | Unidentified | Unidentified | -0.183 | -0.516 | 0.101 | -0.184 |
| 3 | 8 | X_0143 | Archaea | Nanoarchaeota | Nanoarchaeia | Woesearchaeales | Unidentified | Unidentified | -0.186 | -0.294 | 0.194 | -0.193 |
| 3 | 8 | X_0212 | Bacteria | Actinobacteriota | Actinobacteria | Corynebacteriales | Mycobacteriaceae | Mycobacterium | -0.203 | -0.578 | 0.038 | -0.195 |
| 3 | 8 | X_0193 | Bacteria | Proteobacteria | Alphaproteobacteria | Rhodobacterales | Rhodobacteraceae | Defluviimonas | -0.208 | 0.010 | 0.087 | -0.207 |
| 3 | 8 | X_0624 | Bacteria | Patescibacteria | Saccharimonadia | Saccharimonadales | LWQ8 | Unidentified | -0.277 | -0.530 | 0.013 | -0.263 |
| 3 | 8 | X_0188 | Bacteria | Patescibacteria | Unidentified | Unidentified | Unidentified | Unidentified | -0.304 | -0.447 | 0.186 | -0.295 |

| | | | | | | | | | | | | | |
|---|---|---|---|---|---|---|---|---|---|---|---|---|---|
| 3 | 9 | X_0148 | Bacteria | Patescibacteria | Saccharimonadia | Saccharimonadales | Unidentified | Unidentified | | 0.412 | 0.468 | -0.095 | 0.388 |
| 3 | 9 | X_0225 | Bacteria | Bdellovibrionota | Bdellovibrionia | Bdellovibrionales | Bdellovibrionaceae | Bdellovibrio | | 0.399 | 0.097 | -0.133 | 0.402 |
| 3 | 9 | X_0052 | Bacteria | Patescibacteria | Parcubacteria | Unidentified | Unidentified | Unidentified | | 0.336 | 0.320 | -0.184 | 0.334 |
| 3 | 9 | X_0113 | Bacteria | Patescibacteria | Gracilibacteria | Candidatus Peribacteria | Unidentified | Unidentified | | 0.332 | 0.457 | -0.055 | 0.319 |
| 3 | 9 | X_0142 | Bacteria | Patescibacteria | ABY1 | Candidatus Magasanikbacteria | Unidentified | Unidentified | | 0.318 | 0.303 | -0.136 | 0.319 |
| 3 | 9 | X_0122 | Bacteria | Patescibacteria | Parcubacteria | Candidatus Nomurabacteria | Unidentified | Unidentified | | 0.287 | 0.302 | -0.218 | 0.290 |
| 3 | 9 | X_0153 | Bacteria | Proteobacteria | Gammaproteobacteria | Burkholderiales | Comamonadaceae | Inhella | | 0.267 | 0.294 | -0.088 | 0.271 |
| 3 | 9 | X_0093 | Bacteria | Proteobacteria | Gammaproteobacteria | Xanthomonadales | Xanthomonadaceae | Thermomonas | | 0.261 | -0.058 | -0.243 | 0.257 |
| 3 | 9 | X_0171 | Bacteria | Planctomycetota | Planctomycetes | Planctomycetales | Rubinisphaeraceae | Rubinisphaera | | 0.256 | 0.382 | 0.076 | 0.256 |
| 3 | 9 | X_0295 | Bacteria | Bacteroidota | Bacteroidia | Chitinophagales | Chitinophagaceae | Terrimonas | | 0.249 | 0.287 | -0.127 | 0.253 |
| 3 | 9 | X_0139 | Bacteria | Proteobacteria | Gammaproteobacteria | Unidentified | Unidentified | Unidentified | | 0.220 | 0.205 | -0.069 | 0.226 |
| 3 | 9 | X_0119 | Bacteria | Patescibacteria | Parcubacteria | Candidatus Zambryskibacteria | Unidentified | Unidentified | | 0.214 | 0.427 | -0.069 | 0.216 |
| 3 | 9 | X_0593 | Bacteria | Unidentified | Unidentified | Unidentified | Unidentified | Unidentified | | 0.209 | 0.193 | -0.085 | 0.215 |
| 3 | 9 | X_0272 | Bacteria | Spirochaetota | Brevinematia | Brevinematales | Brevinemataceae | Brevinema | | 0.203 | 0.071 | -0.342 | 0.205 |
| 3 | 9 | X_0292 | Bacteria | Dependentiae | Babeliae | Babeliales | UBA12409 | Unidentified | | 0.193 | 0.214 | -0.020 | 0.199 |
| 3 | 9 | X_0230 | Bacteria | Dependentiae | Babeliae | Babeliales | Unidentified | Unidentified | | 0.187 | 0.131 | -0.123 | 0.192 |
| 3 | 9 | X_0174 | Bacteria | Patescibacteria | Saccharimonadia | Saccharimonadales | Unidentified | Unidentified | | 0.169 | 0.483 | -0.120 | 0.173 |
| 3 | 9 | X_0223 | Bacteria | Bacteroidota | Bacteroidia | Chitinophagales | Chitinophagaceae | Niabella | | 0.165 | 0.315 | -0.032 | 0.173 |
| 3 | 9 | X_0038 | Bacteria | Bacteroidota | Bacteroidia | Chitinophagales | Chitinophagaceae | Sediminibacterium | | 0.139 | 0.179 | -0.002 | 0.146 |
| 3 | 9 | X_0088 | Bacteria | Patescibacteria | Parcubacteria | Candidatus Campbellbacteria | Unidentified | Unidentified | | 0.134 | 0.451 | 0.105 | 0.143 |
| 3 | 9 | X_0106 | Bacteria | Bacteroidota | Bacteroidia | Flavobacteriales | Crocinitomicaceae | Fluviicola | | 0.123 | -0.034 | -0.117 | 0.121 |

| | | | | | | | | | | | | |
|---|---|---|---|---|---|---|---|---|---|---|---|---|
| 3 | 9 | X_0214 | Bacteria | Proteobacteria | Alphaproteobacteria | Sphingomonadales | Sphingomonadaceae | Unidentified | 0.120 | 0.110 | -0.029 | 0.125 |
| 3 | 9 | X_0712 | Bacteria | Proteobacteria | Gammaproteobacteria | Burkholderiales | Chitinimonadaceae | Chitinimonas | 0.119 | 0.290 | -0.197 | 0.129 |
| 3 | 9 | X_0058 | Bacteria | Patescibacteria | Parcubacteria | Candidatus Nomurabacteria | Unidentified | Unidentified | 0.114 | 0.299 | -0.053 | 0.124 |
| 3 | 9 | X_0253 | Bacteria | Proteobacteria | Gammaproteobacteria | Unidentified | Unidentified | Unidentified | 0.111 | 0.089 | -0.022 | 0.115 |
| 3 | 9 | X_0537 | Bacteria | Margulisbacteria | Unidentified | Unidentified | Unidentified | Unidentified | 0.095 | 0.255 | 0.084 | 0.105 |
| 3 | 9 | X_0411 | Bacteria | Patescibacteria | Saccharimonadia | Saccharimonadales | Unidentified | Unidentified | 0.088 | 0.357 | 0.012 | 0.101 |
| 3 | 9 | X_0246 | Bacteria | Bacteroidota | Bacteroidia | Chitinophagales | Saprospiraceae | Unidentified | 0.078 | 0.339 | -0.152 | 0.091 |
| 3 | 9 | X_0130 | Bacteria | Patescibacteria | Gracilibacteria | Unidentified | Unidentified | Unidentified | 0.059 | 0.411 | 0.105 | 0.076 |
| 3 | 9 | X_0082 | Bacteria | Bacteroidota | Bacteroidia | Chitinophagales | Saprospiraceae | Haliscomenobacter | 0.052 | 0.408 | -0.193 | 0.069 |
| 3 | 9 | X_0549 | Bacteria | Patescibacteria | Saccharimonadia | Saccharimonadales | Unidentified | Unidentified | 0.014 | 0.040 | -0.215 | 0.017 |
| 3 | 9 | X_0140 | Bacteria | Bacteroidota | Bacteroidia | Chitinophagales | Saprospiraceae | Unidentified | 0.013 | 0.478 | 0.053 | 0.036 |
| 3 | 9 | X_0164 | Bacteria | Proteobacteria | Gammaproteobacteria | Xanthomonadales | Xanthomonadaceae | Thermomonas | 0.004 | 0.529 | 0.188 | 0.031 |
| 3 | 9 | X_0285 | Bacteria | Bacteroidota | Bacteroidia | Chitinophagales | Chitinophagaceae | Unidentified | -0.118 | 0.560 | 0.099 | -0.068 |
| 3 | 10 | X_0070 | Bacteria | Proteobacteria | Gammaproteobacteria | Burkholderiales | Chromobacteriaceae | Vogesella | -0.095 | -0.014 | 0.178 | -0.096 |
| 3 | 10 | X_0191 | Bacteria | Proteobacteria | Gammaproteobacteria | Pseudomonadales | Pseudomonadaceae | Unidentified | -0.173 | -0.029 | 0.052 | -0.174 |
| 3 | 10 | X_0165 | Bacteria | Bacteroidota | Bacteroidia | Flavobacteriales | Weeksellaceae | Chryseobacterium | -0.295 | -0.026 | 0.259 | -0.296 |
| 3 | 11 | X_0090 | Bacteria | Cyanobacteria | Vampirivibrionia | Obscuribacterales | Obscuribacteraceae | Candidatus Obscuribacter | 0.059 | 0.038 | 0.065 | 0.061 |
| 3 | 12 | X_0099 | Bacteria | Planctomycetota | Planctomycetes | Pirellulales | Pirellulaceae | Unidentified | 0.015 | 0.052 | 0.152 | 0.018 |
| 3 | 13 | X_0124 | Bacteria | Proteobacteria | Gammaproteobacteria | Burkholderiales | Leeiaceae | Leeia | 0.026 | 0.153 | -0.064 | 0.034 |
| 3 | 14 | X_0127 | Bacteria | Firmicutes | Bacilli | Mycoplasmatales | Mycoplasmataceae | Unidentified | 0.067 | -0.010 | -0.059 | 0.066 |

| | | | | | | | | | | | | |
|---|---|---|---|---|---|---|---|---|---|---|---|---|
| 3 | 15 | X_0152 | Bacteria | Proteobacteria | Alphaproteobacteria | Rhizobiales | Beijerinckiaceae | Methylobacterium-Methylorubrum | -0.214 | 0.209 | 0.134 | -0.199 |
| 3 | 16 | X_0154 | Bacteria | Bacteroidota | Bacteroidia | Flavobacteriales | Crocinitomicaceae | Fluviicola | 0.052 | -0.500 | -0.017 | 0.019 |
| 3 | 17 | X_0156 | Bacteria | Nitrospirota | Nitrospiria | Nitrospirales | Nitrospiraceae | Nitrospira | 0.062 | 0.117 | 0.083 | 0.068 |
| 3 | 18 | X_0185 | Bacteria | Bdellovibrionota | Bdellovibrionia | Bdellovibrionales | Bdellovibrionaceae | Bdellovibrio | 0.180 | 0.048 | -0.102 | 0.182 |
| 3 | 19 | X_0186 | Bacteria | Proteobacteria | Gammaproteobacteria | Burkholderiales | Oxalobacteraceae | Undibacterium | 0.279 | 0.088 | -0.077 | 0.283 |
| 3 | 20 | X_0203 | Bacteria | Firmicutes | Bacilli | Erysipelotrichales | Erysipelotrichaceae | Turicibacter | 0.078 | 0.119 | 0.104 | 0.083 |
| 3 | 21 | X_0208 | Bacteria | Verrucomicrobiota | Verrucomicrobiae | Verrucomicrobiales | Verrucomicrobiaceae | Prosthecobacter | -0.059 | 0.065 | 0.062 | -0.055 |
| 3 | 22 | X_0227 | Bacteria | Proteobacteria | Gammaproteobacteria | Pseudomonadales | Moraxellaceae | [Agitococcus] lubricus group | 0.055 | 0.179 | -0.063 | 0.063 |
| 3 | 22 | X_0232 | Bacteria | Firmicutes | Clostridia | Lachnospirales | Lachnospiraceae | Epulopiscium | 0.020 | 0.229 | 0.022 | 0.031 |
| 3 | 23 | X_0228 | Bacteria | Patescibacteria | WWE3 | Unidentified | Unidentified | Unidentified | -0.094 | 0.137 | 0.202 | -0.086 |
| 3 | 24 | X_0229 | Bacteria | Patescibacteria | Parcubacteria | Unidentified | Unidentified | Unidentified | -0.092 | 0.179 | 0.128 | -0.081 |
| 3 | 25 | X_0237 | Bacteria | Proteobacteria | Gammaproteobacteria | Diplorickettsiales | Diplorickettsiaceae | Unidentified | -0.120 | 0.041 | 0.007 | -0.118 |
| 3 | 26 | X_0265 | Bacteria | Patescibacteria | Gracilibacteria | Absconditabacteriales (SR1) | Unidentified | Unidentified | 0.203 | 0.064 | -0.080 | 0.206 |
| 3 | 27 | X_0267 | Bacteria | Proteobacteria | Alphaproteobacteria | Rhizobiales | Amb-16S-1323 | Unidentified | 0.002 | 0.213 | 0.222 | 0.013 |
| 3 | 28 | X_0280 | Bacteria | Patescibacteria | Saccharimonadia | Saccharimonadales | Unidentified | Unidentified | 0.048 | -0.057 | 0.051 | 0.044 |
| 3 | 28 | X_0345 | Bacteria | Patescibacteria | Gracilibacteria | Candidatus Peregrinibacteria | Unidentified | Unidentified | -0.027 | 0.142 | 0.065 | -0.020 |
| 3 | 29 | X_0301 | Bacteria | Unidentified | Unidentified | Unidentified | Unidentified | Unidentified | 0.014 | -0.009 | 0.191 | 0.013 |
| 3 | 30 | X_0313 | Bacteria | Bacteroidota | Bacteroidia | Sphingobacteriales | NS11-12 marine group | Unidentified | -0.068 | -0.108 | -0.089 | -0.074 |
| 3 | 31 | X_0340 | Bacteria | Bdellovibrionota | Oligoflexia | 0319-6G20 | Unidentified | Unidentified | -0.065 | 0.120 | -0.001 | -0.058 |
| 3 | 32 | X_0357 | Bacteria | Bacteroidota | Rhodothermia | Rhodothermales | Rhodothermaceae | Unidentified | 0.060 | 0.106 | 0.012 | 0.065 |

| | | | | | | | | | | | | |
|---|---|---|---|---|---|---|---|---|---|---|---|---|
| 3 | 33 | X_0385 | Bacteria | Verrucomicrobiota | Verrucomicrobiae | Opitutales | Opitutaceae | IMCC26134 | 0.068 | 0.106 | 0.159 | 0.073 |
| 3 | 34 | X_0433 | Bacteria | Proteobacteria | Gammaproteobacteria | Burkholderiales | Nitrosomonadaceae | Nitrosomonas | 0.081 | 0.110 | -0.106 | 0.086 |
| 3 | 35 | X_0507 | Bacteria | Bdellovibrionota | Oligoflexia | 0319-6G20 | Unidentified | Unidentified | -0.040 | 0.005 | 0.038 | -0.039 |
| 3 | 36 | X_0627 | Bacteria | Patescibacteria | Saccharimonadia | Saccharimonadales | Unidentified | Unidentified | -0.057 | -0.123 | 0.066 | -0.063 |
| 3 | 37 | X_0935 | Bacteria | Patescibacteria | Parcubacteria | Unidentified | Unidentified | Unidentified | 0.091 | 0.378 | 0.163 | 0.104 |
| 4 | 1 | X_0119 | Bacteria | Patescibacteria | Parcubacteria | Candidatus Zambryskibacteria | Unidentified | Unidentified | 0.435 | 0.208 | 0.256 | 0.280 |
| 4 | 1 | X_0022 | Bacteria | Proteobacteria | Gammaproteobacteria | Burkholderiales | Comamonadaceae | Ottowia | 0.316 | 0.510 | 0.371 | 0.003 |
| 4 | 1 | X_0143 | Archaea | Nanoarchaeota | Nanoarchaeia | Woesearchaeales | Unidentified | Unidentified | 0.271 | 0.260 | 0.181 | 0.111 |
| 4 | 1 | X_0081 | Bacteria | Patescibacteria | Saccharimonadia | Saccharimonadales | Unidentified | Unidentified | 0.222 | 0.338 | 0.285 | 0.027 |
| 4 | 1 | X_0311 | Archaea | Nanoarchaeota | Nanoarchaeia | Woesearchaeales | GW2011_GWC1_47_15 | Unidentified | 0.180 | 0.224 | 0.208 | 0.051 |
| 4 | 1 | X_0003 | Bacteria | Actinobacteriota | Actinobacteria | Frankiales | Sporichthyaceae | Unidentified | 0.164 | 0.525 | 0.093 | -0.120 |
| 4 | 1 | X_0169 | Bacteria | Patescibacteria | Gracilibacteria | Candidatus Peribacteria | Unidentified | Unidentified | 0.119 | 0.496 | 0.430 | -0.139 |
| 4 | 1 | X_0227 | Bacteria | Proteobacteria | Gammaproteobacteria | Pseudomonadales | Moraxellaceae | [Agitococcus] lubricus group | 0.085 | -0.027 | -0.085 | 0.087 |
| 4 | 1 | X_0140 | Bacteria | Bacteroidota | Bacteroidia | Chitinophagales | Saprospiraceae | Unidentified | 0.083 | 0.349 | 0.240 | -0.093 |
| 4 | 1 | X_0019 | Bacteria | Bacteroidota | Bacteroidia | Chitinophagales | Unidentified | Unidentified | 0.081 | 0.207 | -0.070 | -0.026 |
| 4 | 1 | X_0010 | Bacteria | Actinobacteriota | Actinobacteria | Micrococcales | Microbacteriaceae | Aurantimicrobium | 0.059 | 0.521 | 0.113 | -0.198 |
| 4 | 1 | X_0283 | Bacteria | Verrucomicrobiota | Chlamydiae | Chlamydiales | Simkaniaceae | Unidentified | 0.042 | 0.301 | 0.171 | -0.105 |
| 4 | 1 | X_0174 | Bacteria | Patescibacteria | Saccharimonadia | Saccharimonadales | Unidentified | Unidentified | 0.041 | 0.422 | 0.145 | -0.163 |
| 4 | 1 | X_0051 | Bacteria | Bacteroidota | Bacteroidia | Flavobacteriales | Flavobacteriaceae | Flavobacterium | 0.017 | 0.570 | 0.291 | -0.252 |
| 4 | 1 | X_0105 | Bacteria | Verrucomicrobiota | Verrucomicrobiae | Verrucomicrobiales | Rubritaleaceae | Unidentified | -0.006 | 0.486 | 0.293 | -0.230 |
| 4 | 1 | X_0088 | Bacteria | Patescibacteria | Parcubacteria | Candidatus Campbellbacteria | Unidentified | Unidentified | -0.007 | 0.464 | 0.309 | -0.221 |

| | | | | | | | | | | | | |
|---|---|---|---|---|---|---|---|---|---|---|---|---|
| 4 | 1 | X_0018 | Bacteria | Proteobacteria | Gammaproteobacteria | Burkholderiales | Comamonadaceae | Giesbergeria | -0.028 | 0.647 | 0.310 | -0.320 |
| 4 | 1 | X_0007 | Bacteria | Proteobacteria | Alphaproteobacteria | Sphingomonadales | Sphingomonadaceae | Novosphingobium | -0.034 | 0.270 | 0.166 | -0.155 |
| 4 | 1 | X_0243 | Bacteria | Verrucomicrobiota | Chlamydiae | Chlamydiales | Unidentified | Unidentified | -0.038 | 0.243 | 0.077 | -0.146 |
| 4 | 1 | X_0035 | Bacteria | Verrucomicrobiota | Verrucomicrobiae | Unidentified | Unidentified | Unidentified | -0.055 | 0.389 | 0.138 | -0.225 |
| 4 | 1 | X_0727 | Bacteria | Proteobacteria | Gammaproteobacteria | Salinisphaerales | Solimonadaceae | Unidentified | -0.072 | 0.339 | 0.045 | -0.218 |
| 4 | 1 | X_0058 | Bacteria | Patescibacteria | Parcubacteria | Candidatus Nomurabacteria | Unidentified | Unidentified | -0.138 | 0.164 | 0.030 | -0.197 |
| 4 | 1 | X_0015 | Bacteria | Bacteroidota | Bacteroidia | Chitinophagales | Chitinophagaceae | Unidentified | -0.156 | 0.195 | 0.075 | -0.226 |
| 4 | 1 | X_0001 | Bacteria | Bacteroidota | Bacteroidia | Flavobacteriales | Flavobacteriaceae | Flavobacterium | -0.164 | 0.619 | 0.224 | -0.402 |
| 4 | 1 | X_0042 | Bacteria | Bacteroidota | Bacteroidia | Flavobacteriales | Flavobacteriaceae | Flavobacterium | -0.178 | 0.545 | 0.181 | -0.385 |
| 4 | 1 | X_0013 | Bacteria | Bacteroidota | Bacteroidia | Chitinophagales | Chitinophagaceae | Sediminibacterium | -0.241 | 0.507 | 0.195 | -0.420 |
| 4 | 1 | X_0078 | Bacteria | Proteobacteria | Gammaproteobacteria | Alteromonadales | Alteromonadaceae | Rheinheimera | -0.253 | 0.282 | 0.008 | -0.346 |
| 4 | 1 | X_0142 | Bacteria | Patescibacteria | ABY1 | Candidatus Magasanikbacteria | Unidentified | Unidentified | -0.412 | 0.406 | 0.145 | -0.522 |
| 4 | 2 | X_0028 | Bacteria | Firmicutes | Clostridia | Peptostreptococcales-Tissierellales | Peptostreptococcaceae | Romboutsia | 0.577 | -0.469 | -0.255 | 0.670 |
| 4 | 2 | X_0041 | Bacteria | Firmicutes | Bacilli | Erysipelotrichales | Erysipelotrichaceae | Turicibacter | 0.540 | -0.507 | -0.313 | 0.648 |
| 4 | 2 | X_0014 | Bacteria | Firmicutes | Clostridia | Peptostreptococcales-Tissierellales | Peptostreptococcaceae | Paraclostridium | 0.465 | -0.519 | -0.341 | 0.593 |
| 4 | 2 | X_0005 | Bacteria | Actinobacteriota | Actinobacteria | Micrococcales | Microbacteriaceae | Unidentified | 0.460 | -0.156 | -0.162 | 0.475 |
| 4 | 2 | X_0002 | Bacteria | Fusobacteriota | Fusobacteriia | Fusobacteriales | Fusobacteriaceae | Cetobacterium | 0.410 | -0.250 | -0.272 | 0.468 |
| 4 | 2 | X_0166 | Bacteria | Firmicutes | Clostridia | Clostridiales | Clostridiaceae | Clostridium sensu stricto 1 | 0.393 | -0.154 | -0.029 | 0.415 |
| 4 | 2 | X_0029 | Bacteria | Firmicutes | Clostridia | Clostridiales | Clostridiaceae | Clostridium sensu stricto 1 | 0.322 | -0.072 | -0.109 | 0.318 |

| | | | | | | | | | | | | |
|---|---|---|---|---|---|---|---|---|---|---|---|---|
| 4 | 2 | X_0027 | Bacteria | Proteobacteria | Gammaproteobacteria | Enterobacterales | Hafniaceae | Edwardsiella | 0.296 | -0.169 | -0.236 | 0.336 |
| 4 | 2 | X_0020 | Bacteria | Proteobacteria | Gammaproteobacteria | Enterobacterales | Enterobacteriaceae | Plesiomonas | 0.261 | -0.082 | -0.173 | 0.268 |
| 4 | 2 | X_0183 | Bacteria | Firmicutes | Clostridia | Clostridiales | Clostridiaceae | Unidentified | 0.198 | 0.044 | -0.109 | 0.155 |
| 4 | 3 | X_0006 | Bacteria | Bacteroidota | Bacteroidia | Flavobacteriales | Weeksellaceae | Unidentified | 0.193 | 0.020 | -0.128 | 0.162 |
| 4 | 4 | X_0338 | Bacteria | Planctomycetota | Phycisphaerae | Phycisphaerales | Phycisphaeraceae | CL500-3 | 0.424 | 0.257 | 0.234 | 0.243 |
| 4 | 4 | X_0031 | Bacteria | Proteobacteria | Gammaproteobacteria | Burkholderiales | Burkholderiaceae | Polynucleobacter | 0.422 | 0.144 | 0.307 | 0.303 |
| 4 | 4 | X_0017 | Bacteria | Bacteroidota | Bacteroidia | Chitinophagales | Chitinophagaceae | Edaphobaculum | 0.422 | 0.268 | 0.264 | 0.235 |
| 4 | 4 | X_0052 | Bacteria | Patescibacteria | Parcubacteria | Unidentified | Unidentified | Unidentified | 0.418 | 0.241 | 0.161 | 0.247 |
| 4 | 4 | X_0008 | Bacteria | Proteobacteria | Alphaproteobacteria | Rhizobiales | Rhizobiales Incertae Sedis | Unidentified | 0.385 | 0.421 | 0.285 | 0.114 |
| 4 | 4 | X_0069 | Archaea | Nanoarchaeota | Nanoarchaeia | Woesearchaeales | GW2011_GWC1_47_15 | Unidentified | 0.208 | 0.151 | 0.248 | 0.112 |
| 4 | 4 | X_0049 | Bacteria | Bacteroidota | Bacteroidia | Flavobacteriales | Crocinitomicaceae | Fluviicola | 0.165 | 0.189 | 0.167 | 0.055 |
| 4 | 4 | X_0025 | Bacteria | Actinobacteriota | Actinobacteria | Frankiales | Sporichthyaceae | hgcI clade | 0.139 | 0.112 | 0.216 | 0.070 |
| 4 | 4 | X_0130 | Bacteria | Patescibacteria | Gracilibacteria | Unidentified | Unidentified | Unidentified | 0.066 | 0.441 | 0.364 | -0.153 |
| 4 | 4 | X_0084 | Archaea | Crenarchaeota | Nitrososphaeria | Nitrosopumilales | Nitrosopumilaceae | Candidatus Nitrosotenuis | 0.049 | 0.194 | 0.132 | -0.047 |
| 4 | 4 | X_0113 | Bacteria | Patescibacteria | Gracilibacteria | Candidatus Peribacteria | Unidentified | Unidentified | -0.014 | 0.316 | 0.306 | -0.159 |
| 4 | 4 | X_0045 | Bacteria | Fibrobacterota | Fibrobacteria | Fibrobacterales | Fibrobacteraceae | Unidentified | -0.070 | 0.340 | 0.308 | -0.217 |
| 4 | 4 | X_0160 | Bacteria | Patescibacteria | Gracilibacteria | Candidatus Peregrinibacteria | Unidentified | Unidentified | -0.119 | 0.139 | 0.149 | -0.169 |
| 4 | 4 | X_0075 | Bacteria | Patescibacteria | ABY1 | Candidatus Magasanikbacteria | Unidentified | Unidentified | -0.156 | 0.370 | 0.146 | -0.300 |
| 4 | 4 | X_0055 | Bacteria | Patescibacteria | Microgenomatia | Candidatus Woesebacteria | Unidentified | Unidentified | -0.194 | 0.346 | 0.302 | -0.322 |
| 4 | 4 | X_0332 | Bacteria | Proteobacteria | Gammaproteobacteria | Burkholderiales | Burkholderiaceae | Unidentified | -0.271 | -0.047 | -0.038 | -0.218 |
| 4 | 5 | X_0155 | Bacteria | Proteobacteria | Alphaproteobacteria | Rhizobiales | Rhizobiaceae | Unidentified | 0.285 | -0.364 | -0.293 | 0.404 |

| | | | | | | | | | | | | |
|---|---|---|---|---|---|---|---|---|---|---|---|---|
| 4 | 5 | X_0603 | Bacteria | Proteobacteria | Gammaproteobacteria | Burkholderiales | SC-I-84 | Unidentified | 0.253 | -0.296 | -0.032 | 0.352 |
| 4 | 5 | X_0032 | Bacteria | Bacteroidota | Bacteroidia | Sphingobacteriales | env.OPS 17 | Unidentified | 0.192 | -0.480 | -0.292 | 0.373 |
| 4 | 5 | X_0040 | Bacteria | Patescibacteria | Parcubacteria | Candidatus Nomurabacteria | Unidentified | Unidentified | 0.159 | -0.434 | -0.219 | 0.329 |
| 4 | 5 | X_0103 | Bacteria | Bacteroidota | Bacteroidia | Flavobacteriales | Weeksellaceae | Bergeyella | 0.155 | -0.219 | 0.118 | 0.236 |
| 4 | 5 | X_0059 | Bacteria | Verrucomicrobiota | Verrucomicrobiae | Verrucomicrobiales | Rubritaleaceae | Luteolibacter | 0.143 | -0.428 | -0.175 | 0.313 |
| 4 | 5 | X_0047 | Bacteria | Proteobacteria | Gammaproteobacteria | Xanthomonadales | Xanthomonadaceae | Thermomonas | 0.137 | -0.530 | -0.277 | 0.349 |
| 4 | 5 | X_0193 | Bacteria | Proteobacteria | Alphaproteobacteria | Rhodobacterales | Rhodobacteraceae | Defluviimonas | 0.123 | -0.413 | -0.185 | 0.292 |
| 4 | 5 | X_0235 | Bacteria | Planctomycetota | Planctomycetes | Pirellulales | Pirellulaceae | Unidentified | 0.118 | -0.220 | -0.062 | 0.204 |
| 4 | 5 | X_0023 | Bacteria | Bacteroidota | Bacteroidia | Flavobacteriales | Flavobacteriaceae | Flavobacterium | 0.084 | -0.296 | -0.331 | 0.209 |
| 4 | 5 | X_0250 | Bacteria | Armatimonadota | Fimbriimonadia | Fimbriimonadales | Fimbriimonadaceae | Unidentified | 0.079 | -0.427 | -0.187 | 0.262 |
| 4 | 5 | X_0331 | Bacteria | Verrucomicrobiota | Verrucomicrobiae | Pedosphaerales | Pedosphaeraceae | Unidentified | 0.065 | -0.415 | -0.085 | 0.246 |
| 4 | 5 | X_0200 | Bacteria | Bacteroidota | Bacteroidia | Chitinophagales | Chitinophagaceae | Parafilimonas | 0.009 | -0.382 | -0.094 | 0.185 |
| 4 | 5 | X_0091 | Bacteria | Bacteroidota | Bacteroidia | Chitinophagales | Chitinophagaceae | Sediminibacterium | -0.041 | -0.448 | -0.240 | 0.176 |
| 4 | 5 | X_0330 | Bacteria | Proteobacteria | Alphaproteobacteria | Micavibrionales | Unidentified | Unidentified | -0.049 | -0.295 | -0.234 | 0.095 |
| 4 | 5 | X_0472 | Bacteria | Proteobacteria | Alphaproteobacteria | Reyranellales | Reyranellaceae | Reyranella | -0.108 | -0.244 | -0.097 | 0.020 |
| 4 | 5 | X_0187 | Bacteria | Bacteroidota | Bacteroidia | Sphingobacteriales | AKYH767 | Unidentified | -0.114 | -0.371 | -0.191 | 0.079 |
| 4 | 5 | X_0053 | Bacteria | Bacteroidota | Bacteroidia | Chitinophagales | Chitinophagaceae | Unidentified | -0.125 | -0.202 | -0.005 | -0.015 |
| 4 | 5 | X_0012 | Bacteria | Proteobacteria | Gammaproteobacteria | Burkholderiales | Neisseriaceae | Unidentified | -0.126 | -0.515 | -0.365 | 0.144 |
| 4 | 5 | X_0503 | Bacteria | Proteobacteria | Alphaproteobacteria | Rhodobacterales | Rhodobacteraceae | Unidentified | -0.155 | -0.381 | -0.105 | 0.050 |
| 4 | 5 | X_0112 | Bacteria | Bacteroidota | Bacteroidia | Chitinophagales | Saprospiraceae | Unidentified | -0.173 | -0.465 | -0.240 | 0.081 |
| 4 | 5 | X_0011 | Bacteria | Actinobacteriota | Actinobacteria | Micrococcales | Microbacteriaceae | Unidentified | -0.208 | -0.307 | -0.168 | -0.033 |

| | | | | | | | | | | | | |
|---|---|---|---|---|---|---|---|---|---|---|---|---|
| 4 | 5 | X_0158 | Bacteria | Bdellovibrionota | Oligoflexia | 0319-6G20 | Unidentified | Unidentified | -0.219 | -0.447 | -0.196 | 0.034 |
| 4 | 5 | X_0086 | Bacteria | Patescibacteria | Saccharimonadia | Saccharimonadales | Unidentified | Unidentified | -0.301 | -0.469 | -0.373 | -0.017 |
| 4 | 5 | X_0116 | Bacteria | Bacteroidota | Bacteroidia | Chitinophagales | Chitinophagaceae | Edaphobaculum | -0.326 | -0.391 | -0.202 | -0.083 |
| 4 | 5 | X_0348 | Bacteria | Patescibacteria | Parcubacteria | Unidentified | Unidentified | Unidentified | -0.370 | -0.270 | -0.208 | -0.190 |
| 4 | 5 | X_0050 | Bacteria | Patescibacteria | Gracilibacteria | Absconditabacteriales (SR1) | Unidentified | Unidentified | -0.388 | -0.232 | -0.308 | -0.226 |
| 4 | 5 | X_0009 | Bacteria | Proteobacteria | Gammaproteobacteria | Burkholderiales | Comamonadaceae | Unidentified | -0.501 | -0.427 | -0.425 | -0.202 |
| 4 | 6 | X_0016 | Bacteria | Bacteroidota | Bacteroidia | Flavobacteriales | Weeksellaceae | Cloacibacterium | 0.284 | -0.315 | -0.073 | 0.385 |
| 4 | 7 | X_0021 | Bacteria | Deinococcota | Deinococci | Deinococcales | Deinococcaceae | Deinobacterium | -0.024 | 0.223 | 0.038 | -0.124 |
| 4 | 8 | X_0159 | Bacteria | Proteobacteria | Gammaproteobacteria | Burkholderiales | Comamonadaceae | Rhizobacter | 0.588 | -0.119 | 0.062 | 0.572 |
| 4 | 8 | X_0048 | Bacteria | Proteobacteria | Gammaproteobacteria | Steroidobacterales | Steroidobacteraceae | Unidentified | 0.587 | 0.052 | 0.182 | 0.494 |
| 4 | 8 | X_0192 | Bacteria | Bacteroidota | Bacteroidia | Cytophagales | Spirosomaceae | Unidentified | 0.574 | -0.044 | 0.048 | 0.528 |
| 4 | 8 | X_0222 | Bacteria | Nitrospirota | Nitrospiria | Nitrospirales | Nitrospiraceae | Nitrospira | 0.570 | 0.000 | 0.172 | 0.504 |
| 4 | 8 | X_0120 | Bacteria | Proteobacteria | Gammaproteobacteria | Xanthomonadales | Xanthomonadaceae | Luteimonas | 0.547 | -0.287 | -0.013 | 0.597 |
| 4 | 8 | X_0373 | Bacteria | Acidobacteriota | Holophagae | Holophagales | Holophagaceae | Holophaga | 0.545 | -0.221 | -0.020 | 0.574 |
| 4 | 8 | X_0170 | Bacteria | Planctomycetota | Planctomycetes | Isosphaerales | Isosphaeraceae | Unidentified | 0.540 | -0.154 | 0.063 | 0.544 |
| 4 | 8 | X_0046 | Bacteria | Proteobacteria | Gammaproteobacteria | Burkholderiales | Comamonadaceae | Rhizobacter | 0.537 | -0.405 | -0.087 | 0.623 |
| 4 | 8 | X_0528 | Bacteria | Actinobacteriota | Actinobacteria | Propionibacteriales | Nocardioidaceae | Nocardioides | 0.525 | -0.262 | 0.001 | 0.570 |
| 4 | 8 | X_0068 | Bacteria | Patescibacteria | Saccharimonadia | Saccharimonadales | Saccharimonadaceae | TM7a | 0.512 | 0.033 | 0.182 | 0.438 |
| 4 | 8 | X_0057 | Bacteria | Proteobacteria | Alphaproteobacteria | Rhodobacterales | Rhodobacteraceae | Gemmobacter | 0.512 | -0.122 | -0.024 | 0.506 |
| 4 | 8 | X_0104 | Bacteria | Planctomycetota | Planctomycetes | Gemmatales | Gemmataceae | Gemmata | 0.493 | -0.140 | 0.099 | 0.497 |
| 4 | 8 | X_0030 | Bacteria | Bacteroidota | Bacteroidia | Cytophagales | Spirosomaceae | Flectobacillus | 0.468 | -0.438 | -0.027 | 0.576 |

| | | | | | | | | | | | | |
|---|---|---|---|---|---|---|---|---|---|---|---|---|
| 4 | 8 | X_0071 | Bacteria | Proteobacteria | Gammaproteobacteria | Xanthomonadales | Xanthomonadaceae | Thermomonas | 0.457 | -0.478 | 0.016 | 0.578 |
| 4 | 8 | X_0096 | Bacteria | Bacteroidota | Bacteroidia | Cytophagales | Spirosomaceae | Emticicia | 0.449 | 0.034 | 0.123 | 0.382 |
| 4 | 8 | X_0144 | Bacteria | Bacteroidota | Bacteroidia | Unidentified | Unidentified | Unidentified | 0.442 | 0.049 | 0.007 | 0.368 |
| 4 | 8 | X_0110 | Bacteria | Proteobacteria | Alphaproteobacteria | Rhizobiales | Rhizobiales Incertae Sedis | Unidentified | 0.437 | -0.358 | -0.137 | 0.527 |
| 4 | 8 | X_0181 | Bacteria | Proteobacteria | Alphaproteobacteria | Sphingomonadales | Sphingomonadaceae | Novosphingobium | 0.435 | -0.256 | -0.113 | 0.492 |
| 4 | 8 | X_0324 | Bacteria | Acidobacteriota | Blastocatellia | 11-24 | Unidentified | Unidentified | 0.434 | 0.139 | 0.197 | 0.316 |
| 4 | 8 | X_0400 | Bacteria | Bacteroidota | Bacteroidia | Chitinophagales | Chitinophagaceae | Ferruginibacter | 0.429 | -0.051 | 0.024 | 0.403 |
| 4 | 8 | X_0745 | Bacteria | Gemmatimonadota | Gemmatimonadetes | Gemmatimonadales | Gemmatimonadaceae | Unidentified | 0.411 | -0.150 | 0.046 | 0.430 |
| 4 | 8 | X_0060 | Bacteria | Verrucomicrobiota | Verrucomicrobiae | Unidentified | Unidentified | Unidentified | 0.410 | -0.394 | -0.149 | 0.517 |
| 4 | 8 | X_0151 | Bacteria | Proteobacteria | Alphaproteobacteria | Rhizobiales | Rhizobiales Incertae Sedis | Phreatobacter | 0.399 | -0.136 | -0.024 | 0.413 |
| 4 | 8 | X_0268 | Bacteria | Proteobacteria | Alphaproteobacteria | Micropepsales | Micropepsaceae | Unidentified | 0.392 | -0.384 | -0.130 | 0.499 |
| 4 | 8 | X_0215 | Bacteria | Patescibacteria | Saccharimonadia | Saccharimonadales | Unidentified | Unidentified | 0.383 | -0.414 | -0.142 | 0.501 |
| 4 | 8 | X_0149 | Bacteria | Bacteroidota | Kapabacteria | Kapabacteriales | Unidentified | Unidentified | 0.378 | -0.094 | 0.019 | 0.377 |
| 4 | 8 | X_0062 | Bacteria | Planctomycetota | Planctomycetes | Isosphaerales | Isosphaeraceae | Tundrisphaera | 0.371 | -0.349 | -0.015 | 0.470 |
| 4 | 8 | X_0351 | Bacteria | Nitrospirota | Nitrospiria | Nitrospirales | Nitrospiraceae | Nitrospira | 0.365 | -0.172 | 0.038 | 0.398 |
| 4 | 8 | X_0442 | Bacteria | Proteobacteria | Gammaproteobacteria | Unidentified | Unidentified | Unidentified | 0.358 | -0.165 | -0.161 | 0.389 |
| 4 | 8 | X_0365 | Bacteria | Verrucomicrobiota | Verrucomicrobiae | Pedosphaerales | Pedosphaeraceae | Unidentified | 0.342 | -0.256 | -0.058 | 0.412 |
| 4 | 8 | X_0352 | Bacteria | Bacteroidota | Bacteroidia | Chitinophagales | Chitinophagaceae | Terrimonas | 0.338 | -0.108 | 0.008 | 0.348 |
| 4 | 8 | X_0544 | Bacteria | Proteobacteria | Gammaproteobacteria | Xanthomonadales | Rhodanobacteraceae | Rudaea | 0.276 | -0.293 | 0.000 | 0.370 |
| 4 | 8 | X_0136 | Bacteria | Proteobacteria | Alphaproteobacteria | Sphingomonadales | Sphingomonadaceae | Novosphingobium | 0.272 | 0.013 | -0.094 | 0.235 |
| 4 | 8 | X_0242 | Bacteria | Bacteroidota | Bacteroidia | Sphingobacteriales | env.OPS 17 | Unidentified | 0.254 | -0.251 | -0.092 | 0.334 |

| | | | | | | | | | | | | |
|---|---|---|---|---|---|---|---|---|---|---|---|---|
| 4 | 8 | X_0178 | Bacteria | Planctomycetota | Planctomycetes | Gemmatales | Gemmataceae | Unidentified | 0.237 | -0.080 | 0.170 | 0.247 |
| 4 | 9 | X_0034 | Archaea | Crenarchaeota | Nitrososphaeria | Nitrososphaerales | Nitrososphaeraceae | Candidatus Nitrocosmicus | 0.193 | -0.204 | -0.143 | 0.262 |
| 4 | 10 | X_0121 | Bacteria | Bacteroidota | Bacteroidia | Chitinophagales | Chitinophagaceae | Parasediminibacterium | 0.499 | -0.313 | -0.186 | 0.565 |
| 4 | 10 | X_0073 | Bacteria | Proteobacteria | Gammaproteobacteria | Burkholderiales | Comamonadaceae | Curvibacter | 0.247 | -0.212 | -0.208 | 0.313 |
| 4 | 10 | X_0036 | Bacteria | Bacteroidota | Bacteroidia | Cytophagales | Spirosomaceae | Arsenicibacter | 0.243 | -0.444 | -0.239 | 0.399 |
| 4 | 11 | X_0037 | Bacteria | Proteobacteria | Alphaproteobacteria | Sphingomonadales | Sphingomonadaceae | Novosphingobium | 0.307 | 0.140 | 0.095 | 0.203 |
| 4 | 12 | X_0043 | Bacteria | Proteobacteria | Gammaproteobacteria | Burkholderiales | Oxalobacteraceae | Undibacterium | 0.421 | -0.441 | -0.256 | 0.539 |
| 4 | 13 | X_0044 | Bacteria | Unidentified | Unidentified | Unidentified | Unidentified | Unidentified | 0.197 | -0.493 | -0.155 | 0.381 |
| 4 | 14 | X_0064 | Bacteria | Bacteroidota | Bacteroidia | Bacteroidales | Barnesiellaceae | Unidentified | 0.455 | -0.086 | 0.047 | 0.441 |
| 4 | 15 | X_0066 | Bacteria | Proteobacteria | Gammaproteobacteria | Enterobacterales | Enterobacteriaceae | Citrobacter | 0.240 | -0.159 | -0.065 | 0.283 |
| 4 | 16 | X_0070 | Bacteria | Proteobacteria | Gammaproteobacteria | Burkholderiales | Chromobacteriaceae | Vogesella | 0.219 | -0.049 | 0.092 | 0.216 |
| 4 | 17 | X_0072 | Bacteria | Verrucomicrobiota | Verrucomicrobiae | Unidentified | Unidentified | Unidentified | 0.045 | -0.136 | -0.052 | 0.103 |
| 4 | 18 | X_0077 | Bacteria | Bacteroidota | Bacteroidia | Chitinophagales | Chitinophagaceae | Taibaiella | 0.291 | -0.033 | -0.036 | 0.273 |
| 4 | 19 | X_0080 | Bacteria | Proteobacteria | Gammaproteobacteria | Pseudomonadales | Moraxellaceae | Acinetobacter | 0.066 | 0.058 | -0.047 | 0.032 |
| 4 | 20 | X_0085 | Bacteria | Proteobacteria | Gammaproteobacteria | Pseudomonadales | Pseudomonadaceae | Pseudomonas | 0.261 | 0.014 | -0.179 | 0.224 |
| 4 | 21 | X_0090 | Bacteria | Cyanobacteria | Vampirivibrionia | Obscuribacterales | Obscuribacteraceae | Candidatus Obscuribacter | 0.249 | -0.124 | -0.040 | 0.277 |
| 4 | 22 | X_0095 | Bacteria | Patescibacteria | Saccharimonadia | Saccharimonadales | Unidentified | Unidentified | 0.407 | -0.168 | -0.056 | 0.433 |
| 4 | 23 | X_0101 | Bacteria | Bacteroidota | Bacteroidia | Cytophagales | Microscillaceae | Unidentified | -0.061 | -0.226 | 0.050 | 0.052 |
| 4 | 24 | X_0107 | Bacteria | Proteobacteria | Gammaproteobacteria | Aeromonadales | Aeromonadaceae | Aeromonas | -0.008 | -0.125 | -0.269 | 0.051 |
| 4 | 25 | X_0109 | Bacteria | Proteobacteria | Gammaproteobacteria | Pseudomonadales | Moraxellaceae | Acinetobacter | -0.143 | 0.103 | 0.025 | -0.174 |
| 4 | 26 | X_0114 | Bacteria | Verrucomicrobiota | Verrucomicrobiae | Unidentified | Unidentified | Unidentified | -0.076 | -0.158 | -0.161 | 0.007 |

| | | | | | | | | | | | | |
|---|---|---|---|---|---|---|---|---|---|---|---|---|
| 4 | 27 | X_0115 | Bacteria | Patescibacteria | Gracilibacteria | Absconditabacteriales (SR1) | Unidentified | Unidentified | 0.041 | 0.054 | -0.142 | 0.011 |
| 4 | 28 | X_0123 | Bacteria | Verrucomicrobiota | Chlamydiae | Chlamydiales | cvE6 | Unidentified | -0.019 | 0.006 | 0.031 | -0.020 |
| 4 | 29 | X_0124 | Bacteria | Proteobacteria | Gammaproteobacteria | Burkholderiales | Leeiaceae | Leeia | 0.359 | -0.012 | 0.072 | 0.324 |
| 4 | 30 | X_0127 | Bacteria | Firmicutes | Bacilli | Mycoplasmatales | Mycoplasmataceae | Unidentified | 0.398 | 0.098 | -0.064 | 0.305 |
| 4 | 31 | X_0134 | Bacteria | Verrucomicrobiota | Chlamydiae | Chlamydiales | cvE6 | Unidentified | -0.252 | -0.017 | -0.098 | -0.216 |
| 4 | 32 | X_0152 | Bacteria | Proteobacteria | Alphaproteobacteria | Rhizobiales | Beijerinckiaceae | Methylobacterium-Methylorubrum | -0.257 | -0.059 | -0.001 | -0.200 |
| 4 | 33 | X_0182 | Bacteria | Bacteroidota | Bacteroidia | Chitinophagales | Chitinophagaceae | Unidentified | 0.396 | -0.160 | 0.038 | 0.420 |
| 4 | 34 | X_0185 | Bacteria | Bdellovibrionota | Bdellovibrionia | Bdellovibrionales | Bdellovibrionaceae | Bdellovibrio | 0.175 | 0.120 | -0.064 | 0.098 |
| 4 | 35 | X_0186 | Bacteria | Proteobacteria | Gammaproteobacteria | Burkholderiales | Oxalobacteraceae | Undibacterium | 0.153 | -0.074 | -0.048 | 0.170 |
| 4 | 36 | X_0195 | Bacteria | Bdellovibrionota | Bdellovibrionia | Bdellovibrionales | Bdellovibrionaceae | Bdellovibrio | 0.189 | 0.021 | 0.006 | 0.158 |
| 4 | 37 | X_0199 | Bacteria | Proteobacteria | Alphaproteobacteria | Rhodobacterales | Rhodobacteraceae | Unidentified | 0.165 | 0.160 | -0.063 | 0.070 |
| 4 | 38 | X_0214 | Bacteria | Proteobacteria | Alphaproteobacteria | Sphingomonadales | Sphingomonadaceae | Unidentified | 0.135 | -0.003 | -0.095 | 0.121 |
| 4 | 39 | X_0229 | Bacteria | Patescibacteria | Parcubacteria | Unidentified | Unidentified | Unidentified | 0.246 | 0.038 | 0.071 | 0.200 |
| 4 | 40 | X_0265 | Bacteria | Patescibacteria | Gracilibacteria | Absconditabacteriales (SR1) | Unidentified | Unidentified | 0.143 | -0.049 | -0.136 | 0.150 |
| 4 | 41 | X_0267 | Bacteria | Proteobacteria | Alphaproteobacteria | Rhizobiales | Amb-16S-1323 | Unidentified | 0.051 | -0.147 | 0.029 | 0.113 |
| 4 | 42 | X_0278 | Archaea | Crenarchaeota | Nitrososphaeria | Nitrosotaleales | Nitrosotaleaceae | Candidatus Nitrosotalea | 0.022 | 0.056 | 0.045 | -0.007 |
| 4 | 43 | X_0280 | Bacteria | Patescibacteria | Saccharimonadia | Saccharimonadales | Unidentified | Unidentified | 0.120 | 0.129 | 0.186 | 0.045 |
| 4 | 44 | X_0300 | Bacteria | Planctomycetota | Planctomycetes | Isosphaerales | Isosphaeraceae | Aquisphaera | 0.366 | -0.278 | 0.002 | 0.440 |
| 4 | 45 | X_0301 | Bacteria | Unidentified | Unidentified | Unidentified | Unidentified | Unidentified | 0.170 | -0.124 | -0.108 | 0.207 |
| 4 | 46 | X_0334 | Bacteria | Bacteroidota | Bacteroidia | Chitinophagales | Chitinophagaceae | UTBCD1 | 0.158 | -0.395 | -0.082 | 0.312 |

| | | | | | | | | | | | | |
|---|---|---|---|---|---|---|---|---|---|---|---|---|
| 4 | 47 | X_0362 | Bacteria | Bacteroidota | Bacteroidia | Flavobacteriales | Weeksellaceae | Elizabethkingia | 0.040 | 0.064 | 0.075 | 0.005 |
| 4 | 48 | X_0375 | Bacteria | Verrucomicrobiota | Chlamydiae | Chlamydiales | Simkaniaceae | Unidentified | 0.112 | -0.125 | -0.052 | 0.157 |
| 4 | 49 | X_0455 | Bacteria | Verrucomicrobiota | Chlamydiae | Chlamydiales | Simkaniaceae | Candidatus Rhabdochlamydia | 0.168 | -0.080 | -0.013 | 0.186 |
| 4 | 50 | X_0543 | Bacteria | Proteobacteria | Gammaproteobacteria | Diplorickettsiales | Diplorickettsiaceae | Aquicella | 0.138 | 0.066 | 0.102 | 0.091 |
| 4 | 51 | X_0746 | Bacteria | Patescibacteria | Parcubacteria | Unidentified | Unidentified | Unidentified | 0.085 | -0.254 | -0.092 | 0.191 |
| 4 | 52 | X_0903 | Bacteria | Proteobacteria | Alphaproteobacteria | Rickettsiales | SM2D12 | Unidentified | 0.125 | -0.103 | -0.066 | 0.158 |
| 5 | 1 | X_0097 | Bacteria | Actinobacteriota | Actinobacteria | Corynebacteriales | Mycobacteriaceae | Mycobacterium | 0.335 | 0.681 | 0.052 | 0.358 |
| 5 | 1 | X_0008 | Bacteria | Proteobacteria | Alphaproteobacteria | Rhizobiales | Rhizobiales Incertae Sedis | Unidentified | 0.283 | 0.516 | -0.132 | 0.327 |
| 5 | 1 | X_0035 | Bacteria | Verrucomicrobiota | Verrucomicrobiae | Unidentified | Unidentified | Unidentified | 0.253 | 0.580 | 0.062 | 0.302 |
| 5 | 1 | X_0049 | Bacteria | Bacteroidota | Bacteroidia | Flavobacteriales | Crocinitomicaceae | Fluviicola | 0.216 | 0.386 | 0.028 | 0.263 |
| 5 | 1 | X_0015 | Bacteria | Bacteroidota | Bacteroidia | Chitinophagales | Chitinophagaceae | Unidentified | 0.153 | 0.352 | 0.025 | 0.202 |
| 5 | 1 | X_0411 | Bacteria | Patescibacteria | Saccharimonadia | Saccharimonadales | Unidentified | Unidentified | 0.145 | 0.330 | -0.054 | 0.191 |
| 5 | 1 | X_0563 | Bacteria | Desulfobacterota | Unidentified | Unidentified | Unidentified | Unidentified | 0.134 | 0.423 | 0.093 | 0.192 |
| 5 | 1 | X_0022 | Bacteria | Proteobacteria | Gammaproteobacteria | Burkholderiales | Comamonadaceae | Ottowia | 0.113 | 0.394 | -0.181 | 0.170 |
| 5 | 1 | X_0018 | Bacteria | Proteobacteria | Gammaproteobacteria | Burkholderiales | Comamonadaceae | Giesbergeria | 0.099 | 0.529 | -0.023 | 0.173 |
| 5 | 1 | X_0033 | Bacteria | Proteobacteria | Gammaproteobacteria | Xanthomonadales | Xanthomonadaceae | Arenimonas | 0.098 | 0.532 | 0.026 | 0.173 |
| 5 | 1 | X_0105 | Bacteria | Verrucomicrobiota | Verrucomicrobiae | Verrucomicrobiales | Rubritaleaceae | Unidentified | 0.094 | 0.435 | 0.115 | 0.158 |
| 5 | 1 | X_0174 | Bacteria | Patescibacteria | Saccharimonadia | Saccharimonadales | Unidentified | Unidentified | 0.092 | 0.461 | 0.012 | 0.160 |
| 5 | 1 | X_0141 | Bacteria | Planctomycetota | Planctomycetes | Pirellulales | Pirellulaceae | Unidentified | 0.088 | 0.561 | 0.110 | 0.168 |
| 5 | 1 | X_0547 | Bacteria | Patescibacteria | Saccharimonadia | Saccharimonadales | LWQ8 | Unidentified | 0.088 | 0.536 | 0.107 | 0.165 |

| | | | | | | | | | | | | |
|---|---|---|---|---|---|---|---|---|---|---|---|---|
| 5 | 1 | X_0149 | Bacteria | Bacteroidota | Kapabacteria | Kapabacteriales | Unidentified | Unidentified | 0.067 | 0.487 | 0.041 | 0.142 |
| 5 | 1 | X_0255 | Bacteria | Spirochaetota | Leptospirae | Leptospirales | Leptospiraceae | Turneriella | 0.053 | 0.069 | 0.083 | 0.064 |
| 5 | 1 | X_0537 | Bacteria | Margulisbacteria | Unidentified | Unidentified | Unidentified | Unidentified | 0.047 | 0.481 | 0.106 | 0.123 |
| 5 | 1 | X_0324 | Bacteria | Acidobacteriota | Blastocatellia | 11-24 | Unidentified | Unidentified | 0.038 | 0.109 | -0.045 | 0.056 |
| 5 | 1 | X_0013 | Bacteria | Bacteroidota | Bacteroidia | Chitinophagales | Chitinophagaceae | Sediminibacterium | 0.037 | 0.362 | -0.214 | 0.096 |
| 5 | 1 | X_0001 | Bacteria | Bacteroidota | Bacteroidia | Flavobacteriales | Flavobacteriaceae | Flavobacterium | 0.035 | 0.451 | 0.196 | 0.108 |
| 5 | 1 | X_0019 | Bacteria | Bacteroidota | Bacteroidia | Chitinophagales | Unidentified | Unidentified | 0.035 | 0.156 | 0.015 | 0.061 |
| 5 | 1 | X_0007 | Bacteria | Proteobacteria | Alphaproteobacteria | Sphingomonadales | Sphingomonadaceae | Novosphingobium | -0.024 | 0.363 | -0.114 | 0.040 |
| 5 | 1 | X_0122 | Bacteria | Patescibacteria | Parcubacteria | Candidatus Nomurabacteria | Unidentified | Unidentified | -0.032 | 0.175 | -0.113 | -0.001 |
| 5 | 1 | X_0152 | Bacteria | Proteobacteria | Alphaproteobacteria | Rhizobiales | Beijerinckiaceae | Methylobacterium-Methylorubrum | -0.036 | -0.173 | -0.133 | -0.065 |
| 5 | 1 | X_0017 | Bacteria | Bacteroidota | Bacteroidia | Chitinophagales | Chitinophagaceae | Edaphobaculum | -0.093 | 0.384 | 0.013 | -0.019 |
| 5 | 1 | X_0335 | Bacteria | Bdellovibrionota | Bdellovibrionia | Bdellovibrionales | Bdellovibrionaceae | Bdellovibrio | -0.093 | 0.441 | 0.083 | -0.007 |
| 5 | 1 | X_0045 | Bacteria | Fibrobacterota | Fibrobacteria | Fibrobacterales | Fibrobacteraceae | Unidentified | -0.105 | 0.324 | 0.133 | -0.042 |
| 5 | 1 | X_0208 | Bacteria | Verrucomicrobiota | Verrucomicrobiae | Verrucomicrobiales | Verrucomicrobiaceae | Prosthecobacter | -0.131 | 0.021 | 0.062 | -0.125 |
| 5 | 1 | X_0102 | Bacteria | Proteobacteria | Gammaproteobacteria | Burkholderiales | Burkholderiaceae | Polynucleobacter | -0.151 | 0.322 | 0.206 | -0.086 |
| 5 | 1 | X_0469 | Bacteria | Patescibacteria | Parcubacteria | Unidentified | Unidentified | Unidentified | -0.167 | 0.223 | 0.196 | -0.122 |
| 5 | 1 | X_0292 | Bacteria | Dependentiae | Babeliae | Babeliales | UBA12409 | Unidentified | -0.198 | 0.207 | 0.108 | -0.155 |
| 5 | 1 | X_0723 | Bacteria | Bdellovibrionota | Bdellovibrionia | Bdellovibrionales | Bdellovibrionaceae | Bdellovibrio | -0.240 | 0.343 | 0.236 | -0.163 |
| 5 | 1 | X_0302 | Bacteria | Patescibacteria | Gracilibacteria | Candidatus Peribacteria | Unidentified | Unidentified | -0.366 | 0.128 | 0.210 | -0.336 |
| 5 | 1 | X_0081 | Bacteria | Patescibacteria | Saccharimonadia | Saccharimonadales | Unidentified | Unidentified | -0.376 | 0.325 | 0.129 | -0.294 |

| | | | | | | | | | | | | |
|---|---|---|---|---|---|---|---|---|---|---|---|---|
| 5 | 1 | X_0051 | Bacteria | Bacteroidota | Bacteroidia | Flavobacteriales | Flavobacteriaceae | Flavobacterium | -0.423 | -0.071 | 0.205 | -0.428 |
| 5 | 2 | X_0064 | Bacteria | Bacteroidota | Bacteroidia | Bacteroidales | Barnesiellaceae | Unidentified | 0.468 | 0.272 | -0.059 | 0.490 |
| 5 | 2 | X_0027 | Bacteria | Proteobacteria | Gammaproteobacteria | Enterobacterales | Hafniaceae | Edwardsiella | 0.379 | 0.280 | -0.305 | 0.406 |
| 5 | 2 | X_0002 | Bacteria | Fusobacteriota | Fusobacteriia | Fusobacteriales | Fusobacteriaceae | Cetobacterium | 0.359 | 0.163 | -0.269 | 0.377 |
| 5 | 2 | X_0028 | Bacteria | Firmicutes | Clostridia | Peptostreptococcales-Tissierellales | Peptostreptococcaceae | Romboutsia | 0.323 | 0.100 | -0.162 | 0.334 |
| 5 | 2 | X_0029 | Bacteria | Firmicutes | Clostridia | Clostridiales | Clostridiaceae | Clostridium sensu stricto 1 | 0.288 | 0.196 | -0.212 | 0.312 |
| 5 | 2 | X_0014 | Bacteria | Firmicutes | Clostridia | Peptostreptococcales-Tissierellales | Peptostreptococcaceae | Paraclostridium | 0.278 | 0.008 | -0.243 | 0.275 |
| 5 | 2 | X_0041 | Bacteria | Firmicutes | Bacilli | Erysipelotrichales | Erysipelotrichaceae | Turicibacter | 0.272 | 0.113 | -0.145 | 0.286 |
| 5 | 2 | X_0020 | Bacteria | Proteobacteria | Gammaproteobacteria | Enterobacterales | Enterobacteriaceae | Plesiomonas | 0.259 | 0.109 | -0.189 | 0.272 |
| 5 | 2 | X_0010 | Bacteria | Actinobacteriota | Actinobacteria | Micrococcales | Microbacteriaceae | Aurantimicrobium | 0.209 | 0.219 | 0.024 | 0.239 |
| 5 | 2 | X_0166 | Bacteria | Firmicutes | Clostridia | Clostridiales | Clostridiaceae | Clostridium sensu stricto 1 | 0.175 | 0.233 | 0.019 | 0.207 |
| 5 | 2 | X_0096 | Bacteria | Bacteroidota | Bacteroidia | Cytophagales | Spirosomaceae | Emticicia | 0.167 | 0.051 | -0.179 | 0.173 |
| 5 | 2 | X_0356 | Bacteria | Firmicutes | Clostridia | Lachnospirales | Lachnospiraceae | Epulopiscium | 0.162 | 0.231 | -0.008 | 0.195 |
| 5 | 2 | X_0021 | Bacteria | Deinococcota | Deinococci | Deinococcales | Deinococcaceae | Deinobacterium | 0.000 | -0.014 | 0.097 | -0.003 |
| 5 | 2 | X_0005 | Bacteria | Actinobacteriota | Actinobacteria | Micrococcales | Microbacteriaceae | Unidentified | -0.060 | -0.447 | -0.137 | -0.129 |
| 5 | 3 | X_0042 | Bacteria | Bacteroidota | Bacteroidia | Flavobacteriales | Flavobacteriaceae | Flavobacterium | 0.194 | 0.222 | 0.230 | 0.224 |
| 5 | 3 | X_0003 | Bacteria | Actinobacteriota | Actinobacteria | Frankiales | Sporichthyaceae | Unidentified | 0.103 | 0.393 | 0.204 | 0.161 |
| 5 | 4 | X_0253 | Bacteria | Proteobacteria | Gammaproteobacteria | Unidentified | Unidentified | Unidentified | 0.482 | 0.418 | -0.071 | 0.503 |
| 5 | 4 | X_0293 | Bacteria | Proteobacteria | Gammaproteobacteria | Diplorickettsiales | Diplorickettsiaceae | Aquicella | 0.372 | 0.348 | 0.082 | 0.403 |

| | | | | | | | | | | | | |
|---|---|---|---|---|---|---|---|---|---|---|---|---|
| 5 | 4 | X_0463 | Bacteria | Patescibacteria | Parcubacteria | Unidentified | Unidentified | Unidentified | 0.345 | 0.407 | -0.050 | 0.380 |
| 5 | 4 | X_0119 | Bacteria | Patescibacteria | Parcubacteria | Candidatus Zambryskibacteria | Unidentified | Unidentified | 0.292 | 0.365 | -0.015 | 0.330 |
| 5 | 4 | X_0490 | Bacteria | Patescibacteria | Saccharimonadia | Saccharimonadales | LWQ8 | Unidentified | 0.243 | 0.398 | 0.030 | 0.288 |
| 5 | 4 | X_0074 | Bacteria | Bacteroidota | Bacteroidia | Sphingobacteriales | NS11-12 marine group | Unidentified | 0.216 | 0.532 | 0.059 | 0.272 |
| 5 | 4 | X_0479 | Bacteria | Chloroflexi | Anaerolineae | RBG-13-54-9 | Unidentified | Unidentified | 0.155 | 0.294 | 0.127 | 0.197 |
| 5 | 4 | X_0004 | Bacteria | Bacteroidota | Bacteroidia | Flavobacteriales | Flavobacteriaceae | Flavobacterium | 0.053 | 0.454 | 0.119 | 0.124 |
| 5 | 5 | X_0144 | Bacteria | Bacteroidota | Bacteroidia | Unidentified | Unidentified | Unidentified | 0.171 | -0.072 | -0.030 | 0.155 |
| 5 | 5 | X_0269 | Bacteria | Bacteroidota | Bacteroidia | Chitinophagales | Unidentified | Unidentified | 0.162 | 0.081 | 0.027 | 0.173 |
| 5 | 5 | X_0256 | Bacteria | Planctomycetota | Planctomycetes | Gemmatales | Gemmataceae | Unidentified | 0.107 | 0.094 | 0.009 | 0.121 |
| 5 | 5 | X_0099 | Bacteria | Planctomycetota | Planctomycetes | Pirellulales | Pirellulaceae | Unidentified | 0.104 | 0.165 | 0.095 | 0.129 |
| 5 | 5 | X_0012 | Bacteria | Proteobacteria | Gammaproteobacteria | Burkholderiales | Neisseriaceae | Unidentified | 0.102 | -0.281 | -0.048 | 0.048 |
| 5 | 5 | X_0961 | Bacteria | Bacteroidota | Bacteroidia | Chitinophagales | Chitinophagaceae | Unidentified | 0.097 | 0.172 | 0.235 | 0.124 |
| 5 | 5 | X_0034 | Archaea | Crenarchaeota | Nitrososphaeria | Nitrososphaerales | Nitrososphaeraceae | Candidatus Nitrocosmicus | 0.084 | 0.073 | 0.050 | 0.095 |
| 5 | 5 | X_0151 | Bacteria | Proteobacteria | Alphaproteobacteria | Rhizobiales | Rhizobiales Incertae Sedis | Phreatobacter | 0.080 | 0.095 | -0.023 | 0.094 |
| 5 | 5 | X_0357 | Bacteria | Bacteroidota | Rhodothermia | Rhodothermales | Rhodothermaceae | Unidentified | 0.050 | -0.370 | -0.144 | -0.017 |
| 5 | 5 | X_0140 | Bacteria | Bacteroidota | Bacteroidia | Chitinophagales | Saprospiraceae | Unidentified | 0.043 | -0.276 | 0.061 | -0.007 |
| 5 | 5 | X_0057 | Bacteria | Proteobacteria | Alphaproteobacteria | Rhodobacterales | Rhodobacteraceae | Gemmobacter | 0.039 | -0.346 | -0.116 | -0.023 |
| 5 | 5 | X_0245 | Bacteria | Proteobacteria | Gammaproteobacteria | Burkholderiales | SC-I-84 | Unidentified | 0.023 | -0.182 | 0.097 | -0.009 |
| 5 | 5 | X_0082 | Bacteria | Bacteroidota | Bacteroidia | Chitinophagales | Saprospiraceae | Haliscomenobacter | 0.021 | -0.368 | 0.041 | -0.044 |
| 5 | 5 | X_0283 | Bacteria | Verrucomicrobiota | Chlamydiae | Chlamydiales | Simkaniaceae | Unidentified | -0.015 | -0.074 | 0.142 | -0.028 |
| 5 | 5 | X_0006 | Bacteria | Bacteroidota | Bacteroidia | Flavobacteriales | Weeksellaceae | Unidentified | -0.017 | 0.115 | 0.144 | 0.003 |

| | | | | | | | | | | | | |
|---|---|---|---|---|---|---|---|---|---|---|---|---|
| 5 | 5 | X_0267 | Bacteria | Proteobacteria | Alphaproteobacteria | Rhizobiales | Amb-16S-1323 | Unidentified | -0.018 | -0.041 | 0.071 | -0.025 |
| 5 | 5 | X_0009 | Bacteria | Proteobacteria | Gammaproteobacteria | Burkholderiales | Comamonadaceae | Unidentified | -0.025 | -0.489 | -0.225 | -0.105 |
| 5 | 5 | X_0135 | Bacteria | Bacteroidota | Bacteroidia | Chitinophagales | 37-13 | Unidentified | -0.042 | -0.346 | 0.015 | -0.098 |
| 5 | 5 | X_0026 | Bacteria | Bacteroidota | Bacteroidia | Flavobacteriales | Flavobacteriaceae | Flavobacterium | -0.047 | -0.142 | -0.010 | -0.070 |
| 5 | 5 | X_0053 | Bacteria | Bacteroidota | Bacteroidia | Chitinophagales | Chitinophagaceae | Unidentified | -0.062 | -0.168 | 0.145 | -0.089 |
| 5 | 5 | X_0186 | Bacteria | Proteobacteria | Gammaproteobacteria | Burkholderiales | Oxalobacteraceae | Undibacterium | -0.071 | -0.064 | -0.035 | -0.081 |
| 5 | 5 | X_0101 | Bacteria | Bacteroidota | Bacteroidia | Cytophagales | Microscillaceae | Unidentified | -0.074 | -0.121 | 0.059 | -0.093 |
| 5 | 5 | X_0217 | Bacteria | Patescibacteria | Gracilibacteria | Absconditabacteriales (SR1) | Unidentified | Unidentified | -0.081 | -0.036 | -0.023 | -0.086 |
| 5 | 5 | X_0046 | Bacteria | Proteobacteria | Gammaproteobacteria | Burkholderiales | Comamonadaceae | Rhizobacter | -0.086 | -0.229 | -0.061 | -0.122 |
| 5 | 5 | X_0214 | Bacteria | Proteobacteria | Alphaproteobacteria | Sphingomonadales | Sphingomonadaceae | Unidentified | -0.093 | -0.281 | 0.040 | -0.136 |
| 5 | 5 | X_0114 | Bacteria | Verrucomicrobiota | Verrucomicrobiae | Unidentified | Unidentified | Unidentified | -0.098 | -0.578 | -0.107 | -0.178 |
| 5 | 5 | X_0185 | Bacteria | Bdellovibrionota | Bdellovibrionia | Bdellovibrionales | Bdellovibrionaceae | Bdellovibrio | -0.102 | -0.140 | 0.036 | -0.124 |
| 5 | 5 | X_0016 | Bacteria | Bacteroidota | Bacteroidia | Flavobacteriales | Weeksellaceae | Cloacibacterium | -0.115 | -0.203 | 0.060 | -0.146 |
| 5 | 5 | X_0100 | Bacteria | Bacteroidota | Bacteroidia | Flavobacteriales | Flavobacteriaceae | Flavobacterium | -0.128 | -0.584 | -0.045 | -0.203 |
| 5 | 5 | X_0083 | Bacteria | Planctomycetota | Planctomycetes | Isosphaerales | Isosphaeraceae | Unidentified | -0.130 | -0.367 | -0.033 | -0.182 |
| 5 | 5 | X_0124 | Bacteria | Proteobacteria | Gammaproteobacteria | Burkholderiales | Leeiaceae | Leeia | -0.133 | -0.195 | 0.045 | -0.162 |
| 5 | 5 | X_0158 | Bacteria | Bdellovibrionota | Oligoflexia | 0319-6G20 | Unidentified | Unidentified | -0.149 | -0.460 | -0.100 | -0.209 |
| 5 | 5 | X_0195 | Bacteria | Bdellovibrionota | Bdellovibrionia | Bdellovibrionales | Bdellovibrionaceae | Bdellovibrio | -0.150 | -0.325 | -0.037 | -0.196 |
| 5 | 5 | X_0201 | Bacteria | Proteobacteria | Alphaproteobacteria | Rhodobacterales | Rhodobacteraceae | Unidentified | -0.151 | -0.444 | -0.002 | -0.209 |
| 5 | 5 | X_0072 | Bacteria | Verrucomicrobiota | Verrucomicrobiae | Unidentified | Unidentified | Unidentified | -0.161 | -0.276 | -0.198 | -0.199 |
| 5 | 5 | X_0048 | Bacteria | Proteobacteria | Gammaproteobacteria | Steroidobacterales | Steroidobacteraceae | Unidentified | -0.163 | -0.263 | -0.039 | -0.200 |

| | | | | | | | | | | | | |
|---|---|---|---|---|---|---|---|---|---|---|---|---|
| 5 | 5 | X_0246 | Bacteria | Bacteroidota | Bacteroidia | Chitinophagales | Saprospiraceae | Unidentified | -0.178 | -0.361 | -0.007 | -0.225 |
| 5 | 5 | X_0059 | Bacteria | Verrucomicrobiota | Verrucomicrobiae | Verrucomicrobiales | Rubritaleaceae | Luteolibacter | -0.229 | -0.536 | -0.174 | -0.282 |
| 5 | 5 | X_0032 | Bacteria | Bacteroidota | Bacteroidia | Sphingobacteriales | env.OPS 17 | Unidentified | -0.241 | -0.355 | -0.128 | -0.282 |
| 5 | 5 | X_0306 | Bacteria | Bacteroidota | Bacteroidia | Flavobacteriales | Crocinitomicaceae | Fluviicola | -0.309 | -0.394 | -0.251 | -0.347 |
| 5 | 5 | X_0085 | Bacteria | Proteobacteria | Gammaproteobacteria | Pseudomonadales | Pseudomonadaceae | Pseudomonas | -0.354 | -0.525 | -0.104 | -0.387 |
| 5 | 6 | X_0069 | Archaea | Nanoarchaeota | Nanoarchaeia | Woesearchaeales | GW2011_GWC1_47_15 | Unidentified | 0.173 | 0.538 | 0.055 | 0.236 |
| 5 | 6 | X_0883 | Bacteria | Patescibacteria | Parcubacteria | Candidatus Campbellbacteria | Unidentified | Unidentified | 0.133 | -0.140 | -0.010 | 0.106 |
| 5 | 6 | X_0084 | Archaea | Crenarchaeota | Nitrososphaeria | Nitrosopumilales | Nitrosopumilaceae | Candidatus Nitrosotenuis | 0.110 | 0.499 | 0.071 | 0.179 |
| 5 | 6 | X_0107 | Bacteria | Proteobacteria | Gammaproteobacteria | Aeromonadales | Aeromonadaceae | Aeromonas | 0.012 | -0.001 | -0.078 | 0.012 |
| 5 | 6 | X_0350 | Bacteria | Patescibacteria | Gracilibacteria | Candidatus Peribacteria | Unidentified | Unidentified | -0.028 | 0.373 | 0.102 | 0.039 |
| 5 | 6 | X_0655 | Bacteria | Bdellovibrionota | Oligoflexia | Oligoflexales | Unidentified | Unidentified | -0.031 | 0.026 | 0.002 | -0.026 |
| 5 | 6 | X_0264 | Archaea | Aenigmarchaeota | Aenigmarchaeia | Aenigmarchaeales | Unidentified | Unidentified | -0.032 | 0.356 | -0.121 | 0.031 |
| 5 | 6 | X_0218 | Archaea | Nanoarchaeota | Nanoarchaeia | Woesearchaeales | Unidentified | Unidentified | -0.136 | 0.064 | -0.065 | -0.123 |
| 5 | 6 | X_0163 | Bacteria | Actinobacteriota | Actinobacteria | Corynebacteriales | Mycobacteriaceae | Mycobacterium | -0.158 | 0.080 | -0.104 | -0.141 |
| 5 | 6 | X_0659 | Bacteria | Proteobacteria | Gammaproteobacteria | Coxiellales | Coxiellaceae | Coxiella | -0.171 | 0.135 | 0.061 | -0.144 |
| 5 | 6 | X_0055 | Bacteria | Patescibacteria | Microgenomatia | Candidatus Woesebacteria | Unidentified | Unidentified | -0.172 | 0.358 | 0.106 | -0.097 |
| 5 | 6 | X_0175 | Archaea | Micrarchaeota | Micrarchaeia | Micrarchaeales | Unidentified | Unidentified | -0.200 | 0.410 | 0.150 | -0.110 |
| 5 | 6 | X_0138 | Bacteria | Actinobacteriota | Actinobacteria | Corynebacteriales | Mycobacteriaceae | Mycobacterium | -0.201 | 0.055 | -0.138 | -0.188 |
| 5 | 6 | X_0360 | Bacteria | Patescibacteria | Gracilibacteria | Candidatus Peregrinibacteria | Unidentified | Unidentified | -0.202 | 0.067 | -0.043 | -0.187 |
| 5 | 6 | X_0228 | Bacteria | Patescibacteria | WWE3 | Unidentified | Unidentified | Unidentified | -0.210 | 0.310 | 0.042 | -0.143 |
| 5 | 6 | X_0128 | Bacteria | Verrucomicrobiota | Verrucomicrobiae | Chthoniobacterales | Terrimicrobiaceae | FukuN18 freshwater group | -0.210 | 0.047 | -0.069 | -0.199 |

| | | | | | | | | | | | | |
|---|---|---|---|---|---|---|---|---|---|---|---|---|
| 5 | 6 | X_0011 | Bacteria | Actinobacteriota | Actinobacteria | Micrococcales | Microbacteriaceae | Unidentified | -0.231 | 0.085 | -0.224 | -0.212 |
| 5 | 6 | X_0065 | Bacteria | Patescibacteria | Saccharimonadia | Saccharimonadales | Unidentified | Unidentified | -0.234 | -0.151 | -0.235 | -0.254 |
| 5 | 6 | X_0455 | Bacteria | Verrucomicrobiota | Chlamydiae | Chlamydiales | Simkaniaceae | Candidatus Rhabdochlamydia | -0.242 | -0.354 | -0.110 | -0.284 |
| 5 | 6 | X_0123 | Bacteria | Verrucomicrobiota | Chlamydiae | Chlamydiales | cvE6 | Unidentified | -0.265 | -0.030 | 0.042 | -0.266 |
| 5 | 6 | X_0160 | Bacteria | Patescibacteria | Gracilibacteria | Candidatus Peregrinibacteria | Unidentified | Unidentified | -0.266 | 0.295 | 0.007 | -0.200 |
| 5 | 6 | X_0682 | Bacteria | Patescibacteria | Parcubacteria | Candidatus Yanofskybacteria | Unidentified | Unidentified | -0.271 | -0.391 | -0.053 | -0.313 |
| 5 | 6 | X_0134 | Bacteria | Verrucomicrobiota | Chlamydiae | Chlamydiales | cvE6 | Unidentified | -0.273 | 0.153 | -0.006 | -0.239 |
| 5 | 6 | X_0687 | Bacteria | Proteobacteria | Alphaproteobacteria | Micavibrionales | Unidentified | Unidentified | -0.279 | -0.004 | -0.040 | -0.275 |
| 5 | 6 | X_0023 | Bacteria | Bacteroidota | Bacteroidia | Flavobacteriales | Flavobacteriaceae | Flavobacterium | -0.281 | -0.411 | -0.181 | -0.323 |
| 5 | 6 | X_0429 | Bacteria | Patescibacteria | Saccharimonadia | Saccharimonadales | Unidentified | Unidentified | -0.285 | -0.130 | -0.121 | -0.301 |
| 5 | 6 | X_0106 | Bacteria | Bacteroidota | Bacteroidia | Flavobacteriales | Crocinitomicaceae | Fluviicola | -0.329 | -0.482 | -0.107 | -0.367 |
| 5 | 6 | X_0234 | Bacteria | Patescibacteria | Saccharimonadia | Saccharimonadales | Unidentified | Unidentified | -0.333 | 0.169 | 0.127 | -0.295 |
| 5 | 6 | X_0574 | Bacteria | Patescibacteria | Parcubacteria | Unidentified | Unidentified | Unidentified | -0.347 | -0.312 | -0.096 | -0.378 |
| 5 | 6 | X_0280 | Bacteria | Patescibacteria | Saccharimonadia | Saccharimonadales | Unidentified | Unidentified | -0.354 | -0.025 | 0.164 | -0.353 |
| 5 | 6 | X_0090 | Bacteria | Cyanobacteria | Vampirivibrionia | Obscuribacterales | Obscuribacteraceae | Candidatus Obscuribacter | -0.366 | -0.309 | -0.040 | -0.396 |
| 5 | 6 | X_0440 | Bacteria | Patescibacteria | Saccharimonadia | Saccharimonadales | Unidentified | Unidentified | -0.375 | -0.052 | -0.012 | -0.378 |
| 5 | 6 | X_0086 | Bacteria | Patescibacteria | Saccharimonadia | Saccharimonadales | Unidentified | Unidentified | -0.389 | -0.340 | -0.180 | -0.419 |
| 5 | 6 | X_0050 | Bacteria | Patescibacteria | Gracilibacteria | Absconditabacteriales (SR1) | Unidentified | Unidentified | -0.405 | -0.248 | -0.042 | -0.429 |
| 5 | 6 | X_0187 | Bacteria | Bacteroidota | Bacteroidia | Sphingobacteriales | AKYH767 | Unidentified | -0.414 | -0.283 | -0.026 | -0.440 |
| 5 | 6 | X_0334 | Bacteria | Bacteroidota | Bacteroidia | Chitinophagales | Chitinophagaceae | UTBCD1 | -0.427 | -0.149 | 0.071 | -0.442 |

| | | | | | | | | | | | | |
|---|---|---|---|---|---|---|---|---|---|---|---|---|
| 5 | 6 | X_0142 | Bacteria | Patescibacteria | ABY1 | Candidatus Magasanikbacteria | Unidentified | Unidentified | -0.428 | -0.225 | 0.131 | -0.449 |
| 5 | 6 | X_0262 | Bacteria | Patescibacteria | Microgenomatia | Candidatus Amesbacteria | Unidentified | Unidentified | -0.440 | -0.080 | 0.040 | -0.446 |
| 5 | 6 | X_0297 | Bacteria | Planctomycetota | Planctomycetes | Planctomycetales | Rubinisphaeraceae | Unidentified | -0.453 | -0.007 | -0.070 | -0.448 |
| 5 | 6 | X_0040 | Bacteria | Patescibacteria | Parcubacteria | Candidatus Nomurabacteria | Unidentified | Unidentified | -0.460 | -0.238 | 0.064 | -0.481 |
| 5 | 6 | X_0075 | Bacteria | Patescibacteria | ABY1 | Candidatus Magasanikbacteria | Unidentified | Unidentified | -0.491 | -0.134 | 0.140 | -0.502 |
| 5 | 6 | X_0052 | Bacteria | Patescibacteria | Parcubacteria | Unidentified | Unidentified | Unidentified | -0.501 | 0.181 | 0.073 | -0.454 |
| 5 | 7 | X_0169 | Bacteria | Patescibacteria | Gracilibacteria | Candidatus Peribacteria | Unidentified | Unidentified | 0.579 | 0.545 | 0.055 | 0.572 |
| 5 | 7 | X_0095 | Bacteria | Patescibacteria | Saccharimonadia | Saccharimonadales | Unidentified | Unidentified | 0.574 | 0.570 | 0.037 | 0.562 |
| 5 | 7 | X_0327 | Bacteria | Proteobacteria | Gammaproteobacteria | Xanthomonadales | Xanthomonadaceae | Thermomonas | 0.518 | 0.537 | 0.039 | 0.523 |
| 5 | 7 | X_0031 | Bacteria | Proteobacteria | Gammaproteobacteria | Burkholderiales | Burkholderiaceae | Polynucleobacter | 0.507 | 0.602 | -0.106 | 0.502 |
| 5 | 7 | X_0087 | Bacteria | Bacteroidota | Bacteroidia | Chitinophagales | Chitinophagaceae | Unidentified | 0.498 | 0.504 | 0.004 | 0.510 |
| 5 | 7 | X_0198 | Bacteria | Planctomycetota | Planctomycetes | Pirellulales | Pirellulaceae | Unidentified | 0.494 | 0.594 | 0.009 | 0.493 |
| 5 | 7 | X_0493 | Bacteria | Bacteroidota | Bacteroidia | Chitinophagales | Chitinophagaceae | Sediminibacterium | 0.480 | 0.503 | -0.010 | 0.495 |
| 5 | 7 | X_0230 | Bacteria | Dependentiae | Babeliae | Babeliales | Unidentified | Unidentified | 0.455 | 0.565 | -0.054 | 0.467 |
| 5 | 7 | X_0377 | Bacteria | Proteobacteria | Gammaproteobacteria | Xanthomonadales | Xanthomonadaceae | Pseudoxanthomonas | 0.453 | 0.513 | 0.038 | 0.471 |
| 5 | 7 | X_0421 | Bacteria | Bacteroidota | Bacteroidia | Chitinophagales | Saprospiraceae | Unidentified | 0.438 | 0.554 | -0.024 | 0.454 |
| 5 | 7 | X_1121 | Bacteria | Bacteroidota | Bacteroidia | Chitinophagales | Saprospiraceae | Unidentified | 0.435 | 0.384 | -0.068 | 0.462 |
| 5 | 7 | X_0747 | Bacteria | Patescibacteria | Saccharimonadia | Saccharimonadales | LWQ8 | Unidentified | 0.431 | 0.449 | 0.010 | 0.456 |
| 5 | 7 | X_0259 | Bacteria | Proteobacteria | Gammaproteobacteria | Burkholderiales | Chitinimonadaceae | Chitinimonas | 0.426 | 0.338 | -0.046 | 0.453 |
| 5 | 7 | X_0567 | Bacteria | Patescibacteria | Parcubacteria | Unidentified | Unidentified | Unidentified | 0.407 | 0.457 | 0.029 | 0.435 |
| 5 | 7 | X_0238 | Bacteria | Proteobacteria | Gammaproteobacteria | Pseudomonadales | Pseudomonadaceae | Pseudomonas | 0.404 | 0.555 | 0.031 | 0.426 |

| | | | | | | | | | | | | |
|---|---|---|---|---|---|---|---|---|---|---|---|---|
| 5 | 7 | X_0148 | Bacteria | Patescibacteria | Saccharimonadia | Saccharimonadales | Unidentified | Unidentified | 0.401 | 0.618 | -0.021 | 0.416 |
| 5 | 7 | X_0025 | Bacteria | Actinobacteriota | Actinobacteria | Frankiales | Sporichthyaceae | hgcI clade | 0.400 | 0.587 | 0.096 | 0.419 |
| 5 | 7 | X_0391 | Bacteria | Bacteroidota | Bacteroidia | Chitinophagales | Chitinophagaceae | Unidentified | 0.398 | 0.542 | 0.048 | 0.422 |
| 5 | 7 | X_0094 | Bacteria | Bacteroidota | Bacteroidia | Flavobacteriales | Weeksellaceae | Cloacibacterium | 0.394 | 0.461 | -0.066 | 0.423 |
| 5 | 7 | X_0225 | Bacteria | Bdellovibrionota | Bdellovibrionia | Bdellovibrionales | Bdellovibrionaceae | Bdellovibrio | 0.393 | 0.504 | 0.000 | 0.421 |
| 5 | 7 | X_0476 | Bacteria | Proteobacteria | Gammaproteobacteria | Cellvibrionales | Halieaceae | OM60(NOR5) clade | 0.388 | 0.494 | 0.050 | 0.417 |
| 5 | 7 | X_0167 | Bacteria | Bacteroidota | Bacteroidia | Sphingobacteriales | env.OPS 17 | Unidentified | 0.387 | 0.347 | 0.082 | 0.417 |
| 5 | 7 | X_0154 | Bacteria | Bacteroidota | Bacteroidia | Flavobacteriales | Crocinitomicaceae | Fluviicola | 0.375 | 0.435 | 0.065 | 0.408 |
| 5 | 7 | X_0056 | Bacteria | Verrucomicrobiota | Verrucomicrobiae | Unidentified | Unidentified | Unidentified | 0.375 | 0.443 | 0.090 | 0.407 |
| 5 | 7 | X_0210 | Bacteria | Proteobacteria | Alphaproteobacteria | Rhodobacterales | Rhodobacteraceae | Pseudorhodobacter | 0.373 | 0.457 | -0.050 | 0.405 |
| 5 | 7 | X_0559 | Bacteria | Proteobacteria | Gammaproteobacteria | Burkholderiales | Nitrosomonadaceae | Nitrosomonas | 0.371 | 0.521 | 0.041 | 0.401 |
| 5 | 7 | X_0488 | Bacteria | Proteobacteria | Gammaproteobacteria | Burkholderiales | Chitinibacteraceae | Deefgea | 0.369 | 0.353 | 0.121 | 0.401 |
| 5 | 7 | X_0164 | Bacteria | Proteobacteria | Gammaproteobacteria | Xanthomonadales | Xanthomonadaceae | Thermomonas | 0.364 | 0.322 | -0.016 | 0.395 |
| 5 | 7 | X_0178 | Bacteria | Planctomycetota | Planctomycetes | Gemmatales | Gemmataceae | Unidentified | 0.364 | 0.406 | 0.015 | 0.397 |
| 5 | 7 | X_0038 | Bacteria | Bacteroidota | Bacteroidia | Chitinophagales | Chitinophagaceae | Sediminibacterium | 0.359 | 0.526 | 0.169 | 0.391 |
| 5 | 7 | X_0571 | Bacteria | Unidentified | Unidentified | Unidentified | Unidentified | Unidentified | 0.356 | 0.375 | -0.001 | 0.390 |
| 5 | 7 | X_0252 | Bacteria | Proteobacteria | Alphaproteobacteria | Rhodobacterales | Rhodobacteraceae | Defluviimonas | 0.356 | 0.492 | 0.052 | 0.389 |
| 5 | 7 | X_0716 | Bacteria | Planctomycetota | Planctomycetes | Pirellulales | Pirellulaceae | Blastopirellula | 0.339 | 0.475 | 0.113 | 0.376 |
| 5 | 7 | X_0171 | Bacteria | Planctomycetota | Planctomycetes | Planctomycetales | Rubinisphaeraceae | Rubinisphaera | 0.322 | 0.508 | -0.104 | 0.361 |
| 5 | 7 | X_0631 | Bacteria | Unidentified | Unidentified | Unidentified | Unidentified | Unidentified | 0.318 | 0.531 | 0.085 | 0.357 |
| 5 | 7 | X_0572 | Bacteria | Planctomycetota | Planctomycetes | Pirellulales | Pirellulaceae | Pir4 lineage | 0.314 | 0.446 | -0.026 | 0.353 |

| | | | | | | | | | | | | |
|---|---|---|---|---|---|---|---|---|---|---|---|---|
| 5 | 7 | X_0093 | Bacteria | Proteobacteria | Gammaproteobacteria | Xanthomonadales | Xanthomonadaceae | Thermomonas | 0.312 | 0.219 | -0.288 | 0.338 |
| 5 | 7 | X_0126 | Bacteria | Nitrospirota | Nitrospiria | Nitrospirales | Nitrospiraceae | Nitrospira | 0.297 | 0.510 | -0.047 | 0.339 |
| 5 | 7 | X_0303 | Bacteria | Planctomycetota | Planctomycetes | Planctomycetales | Unidentified | Unidentified | 0.268 | 0.436 | -0.006 | 0.312 |
| 5 | 7 | X_0226 | Bacteria | Firmicutes | Clostridia | Clostridiales | Clostridiaceae | Clostridium sensu stricto 1 | 0.267 | 0.350 | -0.144 | 0.307 |
| 5 | 7 | X_0304 | Bacteria | Bacteroidota | Bacteroidia | Chitinophagales | Unidentified | Unidentified | 0.267 | 0.271 | 0.141 | 0.300 |
| 5 | 7 | X_0037 | Bacteria | Proteobacteria | Alphaproteobacteria | Sphingomonadales | Sphingomonadaceae | Novosphingobium | 0.265 | 0.434 | 0.137 | 0.309 |
| 5 | 7 | X_0111 | Bacteria | Verrucomicrobiota | Verrucomicrobiae | Verrucomicrobiales | Rubritaleaceae | Luteolibacter | 0.260 | 0.634 | 0.264 | 0.307 |
| 5 | 7 | X_0080 | Bacteria | Proteobacteria | Gammaproteobacteria | Pseudomonadales | Moraxellaceae | Acinetobacter | 0.256 | 0.272 | -0.038 | 0.290 |
| 5 | 7 | X_0558 | Bacteria | Proteobacteria | Gammaproteobacteria | Oceanospirillales | Unidentified | Unidentified | 0.252 | 0.484 | 0.023 | 0.301 |
| 5 | 7 | X_0127 | Bacteria | Firmicutes | Bacilli | Mycoplasmatales | Mycoplasmataceae | Unidentified | 0.248 | 0.245 | -0.111 | 0.278 |
| 5 | 7 | X_0120 | Bacteria | Proteobacteria | Gammaproteobacteria | Xanthomonadales | Xanthomonadaceae | Luteimonas | 0.246 | 0.406 | 0.012 | 0.291 |
| 5 | 7 | X_0113 | Bacteria | Patescibacteria | Gracilibacteria | Candidatus Peribacteria | Unidentified | Unidentified | 0.241 | 0.518 | 0.125 | 0.292 |
| 5 | 7 | X_0266 | Bacteria | Verrucomicrobiota | Omnitrophia | Omnitrophales | Omnitrophaceae | Candidatus Omnitrophus | 0.232 | 0.583 | 0.024 | 0.285 |
| 5 | 7 | X_0568 | Bacteria | Verrucomicrobiota | Verrucomicrobiae | Chthoniobacterales | Chthoniobacteraceae | Chthoniobacter | 0.220 | 0.465 | 0.056 | 0.271 |
| 5 | 7 | X_0103 | Bacteria | Bacteroidota | Bacteroidia | Flavobacteriales | Weeksellaceae | Bergeyella | 0.219 | 0.408 | -0.013 | 0.266 |
| 5 | 7 | X_0024 | Bacteria | Bacteroidota | Bacteroidia | Sphingobacteriales | Sphingobacteriaceae | Solitalea | 0.214 | 0.474 | -0.011 | 0.267 |
| 5 | 7 | X_0384 | Bacteria | Bacteroidota | Bacteroidia | Chitinophagales | Saprospiraceae | Unidentified | 0.205 | 0.477 | 0.075 | 0.259 |
| 5 | 7 | X_0223 | Bacteria | Bacteroidota | Bacteroidia | Chitinophagales | Chitinophagaceae | Niabella | 0.182 | 0.346 | -0.122 | 0.227 |
| 5 | 7 | X_0516 | Bacteria | Patescibacteria | Gracilibacteria | Candidatus Peregrinibacteria | Unidentified | Unidentified | 0.168 | 0.521 | 0.019 | 0.231 |
| 5 | 8 | X_0043 | Bacteria | Proteobacteria | Gammaproteobacteria | Burkholderiales | Oxalobacteraceae | Undibacterium | -0.003 | -0.154 | 0.053 | -0.029 |
| 5 | 9 | X_0058 | Bacteria | Patescibacteria | Parcubacteria | Candidatus Nomurabacteria | Unidentified | Unidentified | 0.335 | 0.023 | 0.209 | 0.334 |

| | | | | | | | | | | | | |
|---|---|---|---|---|---|---|---|---|---|---|---|---|
| 5 | 10 | X_0066 | Bacteria | Proteobacteria | Gammaproteobacteria | Enterobacterales | Enterobacteriaceae | Citrobacter | -0.011 | 0.150 | -0.105 | 0.015 |
| 5 | 11 | X_0070 | Bacteria | Proteobacteria | Gammaproteobacteria | Burkholderiales | Chromobacteriaceae | Vogesella | -0.005 | 0.317 | 0.060 | 0.050 |
| 5 | 12 | X_0194 | Bacteria | Verrucomicrobiota | Verrucomicrobiae | Verrucomicrobiales | Rubritaleaceae | Luteolibacter | 0.486 | 0.520 | 0.165 | 0.498 |
| 5 | 12 | X_0224 | Bacteria | Verrucomicrobiota | Verrucomicrobiae | Verrucomicrobiales | Rubritaleaceae | Luteolibacter | 0.379 | 0.388 | 0.096 | 0.411 |
| 5 | 12 | X_0156 | Bacteria | Nitrospirota | Nitrospiria | Nitrospirales | Nitrospiraceae | Nitrospira | 0.379 | 0.412 | 0.058 | 0.411 |
| 5 | 12 | X_0077 | Bacteria | Bacteroidota | Bacteroidia | Chitinophagales | Chitinophagaceae | Taibaiella | 0.291 | 0.316 | -0.055 | 0.326 |
| 5 | 13 | X_0088 | Bacteria | Patescibacteria | Parcubacteria | Candidatus Campbellbacteria | Unidentified | Unidentified | 0.110 | 0.164 | -0.136 | 0.136 |
| 5 | 14 | X_0089 | Bacteria | Bacteroidota | Kapabacteria | Kapabacteriales | Unidentified | Unidentified | -0.039 | 0.091 | 0.088 | -0.022 |
| 5 | 15 | X_0109 | Bacteria | Proteobacteria | Gammaproteobacteria | Pseudomonadales | Moraxellaceae | Acinetobacter | -0.161 | -0.238 | -0.172 | -0.195 |
| 5 | 16 | X_0118 | Bacteria | Proteobacteria | Gammaproteobacteria | Alteromonadales | Alteromonadaceae | Rheinheimera | 0.125 | 0.092 | 0.124 | 0.139 |
| 5 | 17 | X_0130 | Bacteria | Patescibacteria | Gracilibacteria | Unidentified | Unidentified | Unidentified | 0.160 | 0.089 | -0.152 | 0.172 |
| 5 | 18 | X_0170 | Bacteria | Planctomycetota | Planctomycetes | Isosphaerales | Isosphaeraceae | Unidentified | 0.018 | 0.084 | 0.005 | 0.032 |
| 5 | 19 | X_0180 | Archaea | Crenarchaeota | Nitrososphaeria | Nitrosopumilales | Nitrosopumilaceae | Candidatus Nitrosotenuis | -0.018 | 0.455 | 0.047 | 0.062 |
| 5 | 20 | X_0183 | Bacteria | Firmicutes | Clostridia | Clostridiales | Clostridiaceae | Unidentified | 0.117 | 0.064 | -0.039 | 0.126 |
| 5 | 21 | X_0190 | Bacteria | Verrucomicrobiota | Verrucomicrobiae | Verrucomicrobiales | Rubritaleaceae | Luteolibacter | 0.011 | 0.048 | -0.014 | 0.019 |
| 5 | 22 | X_0203 | Bacteria | Firmicutes | Bacilli | Erysipelotrichales | Erysipelotrichaceae | Turicibacter | 0.296 | 0.154 | -0.058 | 0.314 |
| 5 | 23 | X_0227 | Bacteria | Proteobacteria | Gammaproteobacteria | Pseudomonadales | Moraxellaceae | [Agitococcus] lubricus group | 0.107 | 0.070 | -0.056 | 0.117 |
| 5 | 24 | X_0231 | Bacteria | Bacteroidota | Bacteroidia | Cytophagales | Spirosomaceae | Runella | 0.162 | 0.343 | 0.058 | 0.209 |
| 5 | 25 | X_0232 | Bacteria | Firmicutes | Clostridia | Lachnospirales | Lachnospiraceae | Epulopiscium | 0.133 | 0.223 | -0.032 | 0.166 |
| 5 | 26 | X_0247 | Bacteria | Verrucomicrobiota | Verrucomicrobiae | Verrucomicrobiales | Rubritaleaceae | Luteolibacter | 0.165 | 0.240 | -0.017 | 0.199 |
| 5 | 27 | X_0260 | Bacteria | Proteobacteria | Gammaproteobacteria | Burkholderiales | Comamonadaceae | Rubrivivax | 0.026 | 0.189 | -0.016 | 0.057 |

| | | | | | | | | | | | | |
|---|---|---|---|---|---|---|---|---|---|---|---|---|
| 5 | 28 | X_0272 | Bacteria | Spirochaetota | Brevinematia | Brevinematales | Brevinemataceae | Brevinema | 0.222 | 0.276 | -0.205 | 0.257 |
| 5 | 29 | X_0274 | Bacteria | Planctomycetota | Planctomycetes | Gemmatales | Gemmataceae | Fimbriiglobus | 0.114 | 0.163 | -0.061 | 0.139 |
| 5 | 30 | X_0279 | Bacteria | Bacteroidota | Bacteroidia | Cytophagales | Spirosomaceae | Emticicia | 0.062 | 0.362 | 0.089 | 0.119 |
| 5 | 31 | X_0295 | Bacteria | Bacteroidota | Bacteroidia | Chitinophagales | Chitinophagaceae | Terrimonas | 0.109 | 0.190 | 0.034 | 0.138 |
| 5 | 32 | X_0325 | Bacteria | Bacteroidota | Bacteroidia | Cytophagales | Microscillaceae | Unidentified | 0.155 | 0.263 | 0.058 | 0.193 |
| 5 | 33 | X_0336 | Bacteria | Firmicutes | Clostridia | Peptostreptococcales-Tissierellales | Peptostreptococcaceae | Terrisporobacter | 0.140 | 0.116 | -0.031 | 0.157 |
| 5 | 34 | X_0342 | Bacteria | Planctomycetota | Planctomycetes | Pirellulales | Pirellulaceae | Unidentified | -0.027 | 0.065 | 0.109 | -0.015 |
| 5 | 35 | X_0363 | Bacteria | Proteobacteria | Gammaproteobacteria | Xanthomonadales | Rhodanobacteraceae | Dokdonella | 0.202 | 0.248 | 0.044 | 0.236 |
| 5 | 36 | X_0370 | Bacteria | Patescibacteria | Saccharimonadia | Saccharimonadales | Unidentified | Unidentified | -0.126 | 0.173 | -0.020 | -0.093 |
| 5 | 37 | X_0385 | Bacteria | Verrucomicrobiota | Verrucomicrobiae | Opitutales | Opitutaceae | IMCC26134 | -0.016 | 0.089 | -0.011 | 0.000 |
| 5 | 38 | X_0393 | Bacteria | Bacteroidota | Bacteroidia | Chitinophagales | Saprospiraceae | Unidentified | 0.356 | 0.393 | 0.044 | 0.390 |
| 5 | 39 | X_0412 | Bacteria | Firmicutes | Clostridia | Lachnospirales | Lachnospiraceae | Epulopiscium | 0.005 | 0.017 | -0.109 | 0.008 |
| 5 | 40 | X_0419 | Bacteria | Patescibacteria | ABY1 | Candidatus Magasanikbacteria | Unidentified | Unidentified | 0.176 | 0.134 | -0.053 | 0.195 |
| 5 | 41 | X_0441 | Bacteria | Planctomycetota | Planctomycetes | Pirellulales | Pirellulaceae | Pir4 lineage | -0.088 | 0.223 | 0.027 | -0.046 |
| 5 | 42 | X_0458 | Bacteria | Verrucomicrobiota | Verrucomicrobiae | Pedosphaerales | Pedosphaeraceae | DEV008 | -0.153 | 0.064 | -0.168 | -0.139 |
| 5 | 43 | X_0481 | Bacteria | Patescibacteria | Saccharimonadia | Saccharimonadales | Unidentified | Unidentified | -0.108 | 0.291 | 0.163 | -0.052 |
| 5 | 44 | X_0506 | Bacteria | Patescibacteria | Saccharimonadia | Saccharimonadales | Unidentified | Unidentified | 0.238 | 0.060 | 0.004 | 0.245 |
| 5 | 45 | X_0519 | Bacteria | Patescibacteria | ABY1 | Candidatus Uhrbacteria | Unidentified | Unidentified | 0.104 | 0.260 | 0.057 | 0.143 |
| 5 | 46 | X_0562 | Bacteria | Acidobacteriota | Vicinamibacteria | Vicinamibacterales | Vicinamibacteraceae | Unidentified | 0.329 | 0.263 | -0.094 | 0.358 |
| 5 | 47 | X_0927 | Bacteria | Bacteroidota | Bacteroidia | Chitinophagales | Saprospiraceae | OLB8 | 0.197 | 0.298 | 0.055 | 0.236 |
